# Integrated Ising Model with global inhibition for decision making

**DOI:** 10.1101/2024.11.18.624088

**Authors:** Olga Tapinova, Tal Finkelman, Tamar Reitich-Stolero, Rony Paz, Assaf Tal, Nir S. Gov

## Abstract

Humans and other organisms make decisions choosing between different options, with the aim to maximize the reward and minimize the cost. The main theoretical framework for modeling the decision-making process has been based on the highly successful drift-diffusion model, which is a simple tool for explaining many aspects of this process. However, new observations challenge this model. Recently, it was found that inhibitory tone increases during high cognitive load and situations of uncertainty, but the origin of this phenomenon is not understood. Motivated by this observation, we extend a recently developed model for decision making while animals move towards targets in real space. We introduce an integrated Ising-type model, that includes global inhibition, and use it to explore its role in decision-making. This model can explain how the brain may utilize inhibition to improve its decision-making accuracy. Compared to experimental results, this model suggests that the regime of the brain’s decision-making activity is in proximity to a critical transition line between the ordered and disordered. Within the model, the critical region near the transition line has the advantageous property of enabling a significant decrease in error with a small increase in inhibition and also exhibits unique properties with respect to learning and memory decay.

---

Decision making is a dynamic cognitive process that results in the selection of a course of action or formation of a categorical choice [1]. The theoretical description of the decision-making process has been attempted on several levels. There are models that describe neuronal networks that include both excitatory and inhibitory neurons and their dynamics [2, 3]. On a more abstract level there is the successful drift-diffusion model (DDM), which assumes that the difference in evidence that is accumulated for each of the options (mostly in binary decisions) drives the decision process [4, 5]. Within this model, decision making is described by a stochastic diffusion process of a “decision variable” (DV), in addition to a drift which represents the net external evidence (bias) in favor of one of the options. In each decision-making process, the DV moves according to the drift-diffusion dynamics until it reaches one of two thresholds, which encode the two abstract alternatives, and a decision occurs [6]. The DDM describes the main properties of observed decision-making dynamics and explains the principles of the speed-accuracy trade-off [3, 7]. However, due to the simplicity of the DDM, there is only one dynamic parameter which controls many of the results, and it is not able to explain more intricate effects without ad-hoc assumptions, such as asymmetric and time-dependent thresholds, and variable drift [4, 8, 9].

Recent observations found an essential role for global inhibition during the decision-making process, with higher levels of the inhibitory neurotransmitter (GABA) detected in conditions of higher uncertainty [10]. Including the role of inhibition motivated us to develop a new model based on a recently proposed Ising model for animal decision making while moving [11, 12]. In this model, the real-space targets which the animal aims to reach are represented by groups of Ising spins that interact ferromagnetically within each group, but their inter-group interactions become inhibitory for large relative angles. The model successfully predicted the bifurcations during collective motion of animal groups [11], and single animal movement in space towards static targets [12, 13] or moving conspecifics [14]. The Ising model was previously applied to study cognitive processes and the behavior of neural networks during decision making [9] and memory [15, 16].

Here we extend our spin-based spatial decision-making model [11, 12], to include the effects of global inhibition and use it to describe the abstract decision making process in the brain. Our Integrated Ising Model (IIM) gives us an underlying mechanism that drives the random walk process. However, unlike the DDM, which relies on simple (Brownian) random-walk diffusion, our Ising model has an ordered phase in which the random walk changes from simple diffusion to run-and-tumble (RnT) dynamics near the transition line [11]. In particular, we find that the regime close to the phase transition within the ordered phase may have advantageous properties for decision making and can explain the observed role of global inhibition in this process. By comparing our model with two sets of independent experiments, we demonstrate that the IIM in the critical regime can better explain the observations, such as the relation between error and reaction time and the effects of increased global inhibition (related to the measured GABA signal), compared to the DDM.

## THEORETICAL MODEL

The Ising spin model, first elaborated for a system of magnetic spins [17], was previously utilized to describe a decision-making process occurring in a single brain, the dynamics of neural networks, and the brain’s physiological state [15, 18–20], and recently to describe animal movement [11–14]. Considering a two-choice decision task in this paper, we investigate the decision-making process in a single brain in the presence of global inhibition.

We assume that the decision-making circuit can be described in an abstract manner by a network of *N* spins, divided into two equal competing groups of spins (I, II), which encode the goal they refer to (fig. 1A(i)). Each spin represents a single neuron (or a group of neurons) in the brain, which can be in either one of two states: “on” or “off”, *σ*_*i*_ = 1, 0, corresponding to neurons in the firing or resting state, respectively. We assume that interactions between the spins are excitatory within the same group and inhibitory between the two groups, in a fully-connected network, neglecting their spatial organization [20].

**Figure 1:**
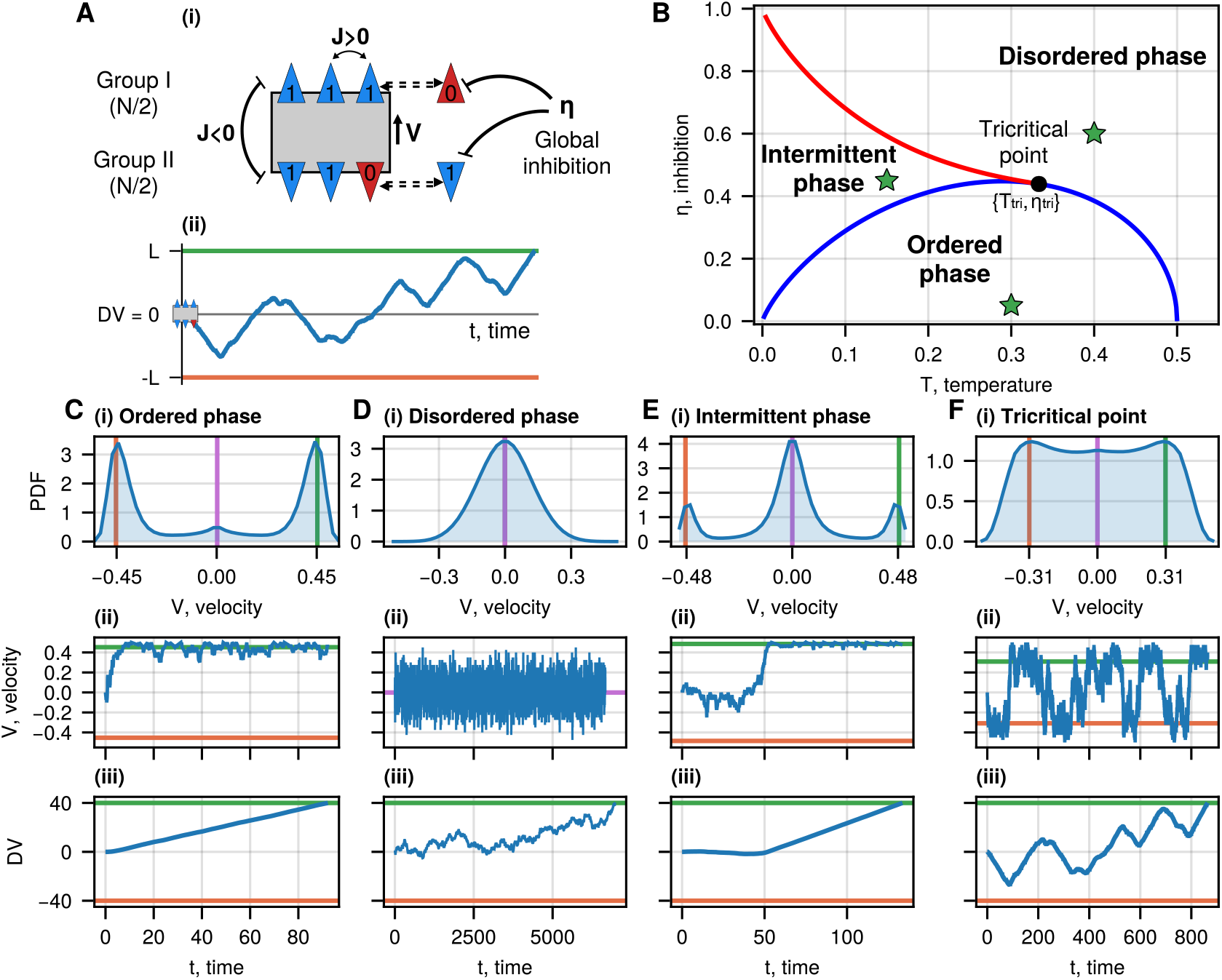
The Integrated Ising model (IIM) for binary decisions (no bias, *ϵ*_1,2_ = 0) [11]. A(i) The spin system is divided into two equal groups, corresponding to the two options of a decision task. The blue triangles represent spins that correspond to firing neurons (*σ* = 1, “on”), while the red ones represent spins that correspond to non-firing neurons (*σ* = 0, “off”). Triangles pointing in the same direction are spins of the same group and have excitatory interactions eq. (1), while spins from one group tend to suppress the other group via cross-inhibition interactions. A(ii) Typical trajectory of the decision variable (DV) in the IIM. The velocity *V* of the DV coordinate is given by the stochastic equations (eq. (3)). The decision occurs when the DV reaches one of the thresholds (green and orange horizontal lines) representing the two options, respectively. The initial conditions are of DV = 0 and all spins in their “off” state. B Phase diagram of the IIM. The blue line denotes the second-order transition (solution of eq. (4)), bounding the ordered phase. The red line denotes the first-order transition, bounding the intermittent phase. Outside these regions, the system is in the disordered phase. The black circle denotes the tricritical point: *η*_tri_ = 0.439, *T*_tri_ = 0.333. The green stars denote the parameters for which the velocity distribution and samples of dynamics are shown in panels C-F. C, D E, F Velocity distributions (i) and examples of the time evolution of the DV’s velocity (ii) and the DV (iii) during the decision-making process. In the ordered phase: *T* = 0.3, *η* = 0.05; the intermittent phase: *T* = 0.15, *η* = 0.45; near the tricritical point: *T* = 0.29, *η* = 0.45; in the disordered phase: *T* = 0.4, *η* = 0.6. The green and orange lines in (i) and indicate the positive and negative MF velocities, respectively (solutions of eq. (4)), while the purple line is the average velocity. The horizontal green and orange lines in (iii) indicate the decision thresholds.

Similar to the regular DDM, we relate the decision-making process to an abstract decision variable (DV) which integrates neuronal firing over time [21, 22], and moves between two fixed and equal thresholds, each encoding one of the options of the two-choice task (fig. 1A(ii)). In the IIM, the DV value either increases or decreases depending on the relative states of the two groups of spins, and when the DV reaches one of the threshold values the decision is reached (fig. 1A(ii)). Our IIM, therefore, is similar to the DDM, but its dynamics can be significantly different, as we show below.

We now introduce the IIM Hamiltonian including a global inhibition a signal from external inhibitory neurons that equally affect all neurons involved in the decision-making process, promoting them to revert to their resting state [10], similar to an external magnetic field applied to a system of magnetic spins

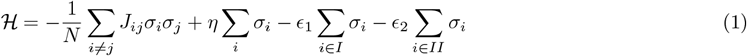

where *J*_*ij*_ is the coupling constant, which equals the multiplication of the preferred directions of spins *i* and *j*, such that *J*_*ij*_ = +1 (ferromagnetic interaction) if they are in the same group, and *J*_*ij*_ = −1 (anti-ferromagnetic interaction) if they are in competing groups [23–25]. We assume here that *J*_*ij*_ is symmetric for simplicity and show the effects of asymmetric interactions in the SI (SI section S1). The first term in eq. (1) gives the summation over all distinct pairs of interacting spins. The second term represents the global inhibition field of strength *η >* 0, which favors the spin state “off”. The last two terms introduce biases in favor of either one of the options (*ϵ*_1*/*2_ *>* 0), applied as external fields.

We use Glauber’s dynamics for the transition rates of the spins [26] (without any biases *ϵ*_1*/*2_ = 0)

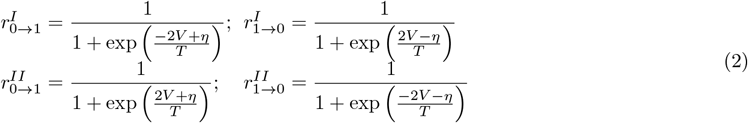

where *T* is the parameter that describes the ratio between the strength of noise in the system and interactions between the spins, playing the role of temperature in the model [27, 28]. The rates in eq. (2) are multiplied by a constant, representing rate units, set to 1.

We define the velocity of the DV as the difference between the fractions of turned “on” spins in the two groups: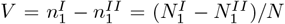. It describes the speed at which the integrator moves along its internal coordinate towards either one of the abstract targets (threshold values, fig. 1A(ii)). The dynamics of 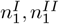 are derived using the rates in eq. (2)

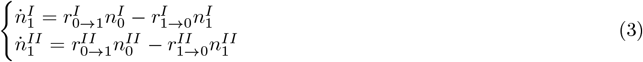

We use these equations and the Gillespie algorithm [29] to numerically simulate the changes in the states of the neural populations. For more details on the simulations and choice of the parameters, see SI section S2. At the steady state, we obtain the mean-field (MF) equation, which solutions give the steady-state values for the DV velocity

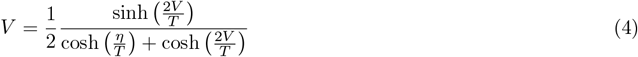

Expanding eq. (4) up to the third order at *V* = 0, we get the condition for the phase transition, at which the zero solution becomes unstable. It gives the second-order transition line on the phase diagram (the blue line in fig. 1B)

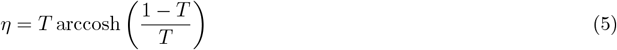

The area under the blue curve (fig. 1B) refers to the ordered phase, where one of the spin groups prevails while the other group is inhibited. It corresponds to two stable non-zero solutions for *V* (fig. 1C). The other transition line, of first-order nature (the red line in fig. 1B, defines a phase (between the red and blue lines) where the zero solution *V* = 0 exists in addition to two stable non-zero solutions. We find the red transition line by solving eq. (4) and 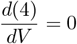 simultaneously. The area above the blue and red transition lines refers to the disordered phase, where only *V* = 0 is stable (fig. 1D(i)). The region between the blue and red curves corresponds to an intermittent phase, where both the zero and non-zero solutions for *V* coexist (fig. 1E(i)).

The tricritical point indicates where the second-order phase transition curve meets the first-order phase transition curve. We find it by solving eq. (4), 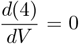, and 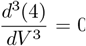 simultaneously, which gives *T*_tri_ ≈ 0.333, *η*_tri_ ≈ 0.439 (the black point in fig. 1B).

In the low temperature and inhibition regime of the ordered phase (below the blue transition line), the DV moves mostly in a ballistic-like trajectory (fig. 1C(ii-iii)), where one group “wins” and inhibits the other. As the system’s temperature and inhibition approach the critical values, the positive and negative solutions for the velocity converge to the zero solution *V* = 0, and the motion transforms into the disordered process (fig. 1D(ii-iii)). In the intermittent regime, the velocity solution can switch between the zero solution to a non-zero value (fig. 1E(ii-iii)). Within the ordered phase, but close to the blue transition line, we find dynamics of the DV to be RnT type, as shown, for example, at the tricritical point (fig. 1F).

The initial conditions for the simulations shown in this paper were for all the spins in their zero configuration. In the SI section S2, we show the results when the initial conditions are such that the spins are in a random initial configuration. The results are not sensitive to this choice of initial conditions unless the system is at very low temperatures.

## PROPERTIES OF DECISION-MAKING PROCESSES IN THE IIM

We now explore the characteristics of the IIM decision-making processes in the presence of bias, which we take here to favor only the option represented by the threshold at *DV* = +*L* (*ϵ*_1_ ≥ 0, *ϵ*_2_ = 0). Therefore, only the Glauber rates of the first group are modified (eq. (2))

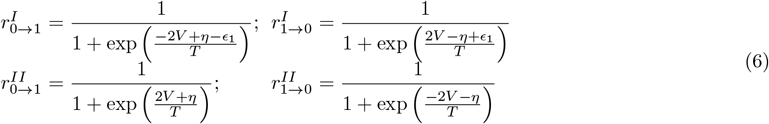

These equations give us the modified MF equation for the velocity of the DV

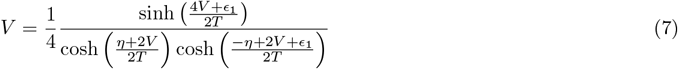

Solving it numerically, we demonstrate the effect of the bias on the velocity distribution fig. 2A(i). The model trajectories of the DV in the presence of the bias (see for example fig. 2A(ii-iii)), allow us to extract the probability of arriving at the biased decision (“correct” decision, when DV hits the threshold at +*L*) and the distribution of “reaction time” (RT), which is the time until a decision is made when the DV reaches either of the two thresholds at ±*L* (see fig. S11 in SI section S3). Throughout the paper, by “RT”, we mean the average reaction time, and the probability of error is the fraction of processes where the DV reached the unfavorable negative threshold at −*L*. Since we want to explore the dependence of the decision-making process on the dynamics of the DV in different parts of the IIM phase diagram, we fix the value of *L* (see also SI section S2 for other values).

**Figure 2:**
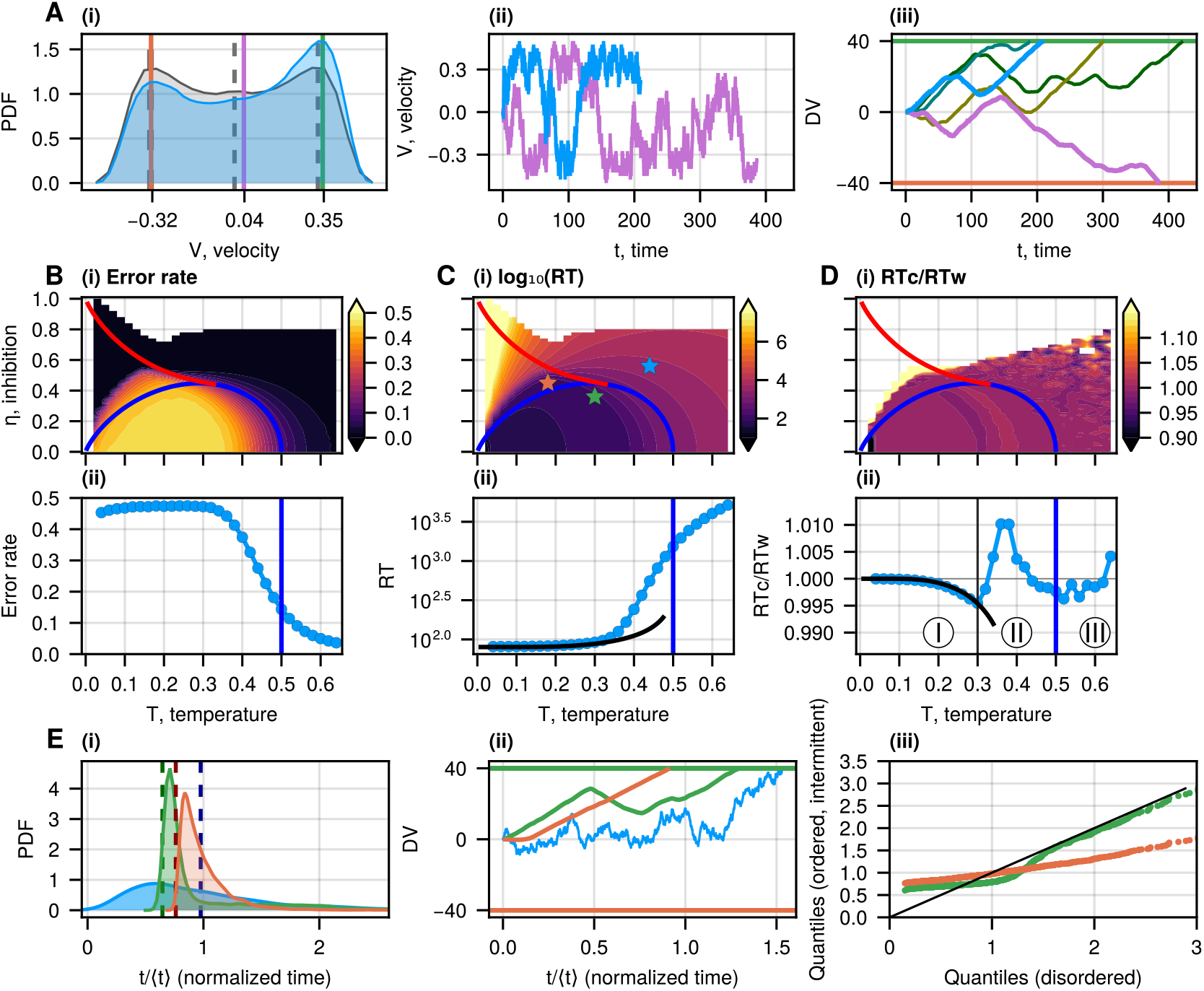
Decision-making dynamics in the IIM. A(i) The probability density of velocity in the IIM near the tricritical point (*T* = 0.29, *η* = 0.44), in the presence of a small constant bias (blue contour, *ϵ*_1_ = 0.01) and without bias (grey contour, *ϵ*_1_ = 0). The purple vertical line indicates the average velocity. The green and orange vertical lines indicate the MF solutions of eq. (7), which coincide with the blue distribution’s peaks. The dashed lines denote the same quantities for the unbiased case. A(ii) Typical velocity dynamics for the trajectories corresponding to the correct (blue) and wrong (purple) choices (same parameters as in the biased case in A(ii)). A(iii) Typical trajectories for the correct choices when the DV reaches the positive threshold (green horizontal line) and the wrong choices when the DV reaches the negative threshold (orange horizontal line). The blue and purple curves correspond to the same colored lines in A(ii). B(i) Error rate, C(i) reaction time (RT), and D(i) the RT ratio in the correct and wrong decisions (RT_c_/RT_w_) as functions of the system’s parameters (*η, T*) at a fixed bias *ϵ*_1_ = 0.01, presented as heatmaps. The red and blue lines on the heatmaps denote the first and second-order transitions, respectively (fig. 1B). B(ii) Error rate, C(ii) RT, and D(ii) the ratio RT_c_/RT_w_ at *η* = 0 as functions of temperature *T* (and same bias as above). The blue vertical line indicates the critical temperature (*T* = 0.5) corresponding to the second-order phase transition. We denote three regimes on the ratio: zone III is above the transition line (the disordered phase). In the ordered phase, we denote a change in the trend of the ratio RT_c_/RT_w_ by the vertical black line. In zone I, the ratio RT_c_/RT_w_ decreases with increasing *T*, while in zone II, it has a non-monotonous dependence on the temperature. The black curve in C(ii), D(ii) gives the theoretical behavior of the RT and the RT_c_/RT_w_ at low temperatures using the ballistic approximation (eq. (8), eq. (9)). E(i) The RT distributions (normalized by the mean RT) for the three regimes (denoted by stars in C(i)), with the vertical dashed lines indicating the theoretical ballistic RT given by 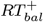 (eq. (8)). E(ii) Typical trajectories during the decision-making process corresponding to the RT distributions shown in E(i). E(iii) Comparison of the RT distributions of the ordered (green) and intermittent (orange) regimes with the disordered regime (black identity line) using a quantile-quantile representation.

For all regions of the phase space, the error rate and the mean RT decrease as the bias grows (see SI section S2). It happens because the positive bias increases the probability of the spin activation in the first group, and hence, the positive MF velocity increases (solution of eq. (7)), compared to the negative solution (see the green and orange vertical lines in fig. 2A(i)).

At fixed bias, the error rate decreases with increasing temperature and inhibition while the RT increases, as shown in fig. 2. Indeed, the error rate is very low in the disordered phase and partly in the intermittent phase (fig. 2B). It happens because, in the disordered phase, the DV moves with low velocity, which approaches zero as the bias diminishes. Therefore, the DV slowly drifts towards the correct threshold, leading to fewer mistakes compared to the ballistic or RnT motion in the ordered phase.

However, the RT grows drastically with temperature and inhibition fig. 2C. This behavior of the model introduces the speed-accuracy trade-off, suggesting that an optimal range of parameters could be where the error is reasonably small while the RT is still not too large. Above the transition line, in the disordered phase, the DV in our model has a low diffusion coefficient [11], thereby giving rise to slow and accurate decisions. Below the transition line, in the ordered phase, our model gives a diffusion coefficient that increases with decreasing temperature (approaching ballistic motion at low *T*), giving rise to fast and inaccurate decisions.

Another property that we can compare to the DDM is the ratio between the RTs of the correct (RT_c_) and wrong (RT_w_) decisions. This ratio (RT_c_/RT_w_) is strictly equal to one for the regular DDM with symmetric thresholds (see the details in SI section S3), but in the IIM we find that there can be deviations from this strict equality (fig. 2D). In fig. 2D(ii), we plot this ratio as a function of temperature for zero inhibition and find three regimes of behavior, depending on the type of motion of the DV: ballistic (I), run-and-tumble (II), and diffusion (III).

At low temperatures (zone I in fig. 2D(ii)), the DV’s motion is ballistic until it reaches the threshold (fig. 1C(iii)). We can estimate the mean RT for the ballistic trajectories that reached the positive or the negative threshold as

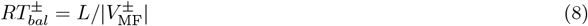

where 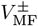 are the solutions of the MF equation (eq. (7)). The ratio of these RTs (marked as the black line in zone I in fig. 2D(ii)) is given by

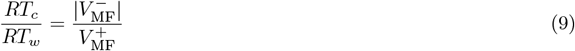

In the biased case, the velocity towards the positive threshold 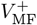 is larger than towards the negative threshold 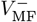 so that the ratio RT_c_/RT_w_ is lower than 1 (the MF ratio is shown by the solid black curve in fig. 2D(ii), see also SI section S3).

In the disordered regime of the IIM (zone III in fig. 2D(ii)), the motion of the DV is identical to the DDM [4, 5] (fig. 1D(iii)), and the RTs can be calculated analytically, giving rise to a ratio equal to one (shown in SI section S3).

In the region of the ordered phase, close to the second-order transition line (zone II in fig. 2D(ii)), the system’s trajectories consist of intervals of movement with almost constant velocity, interrupted by changes in the direction of motion, which can be approximated by the RnT motion (see details in SI section S3, fig. S15). We find that the RnT motion, where the bias is introduced by unequal flipping rates (higher flipping rate towards the correct positive threshold) while keeping the run velocity equal in both directions, results in a RT ratio equal to one (see SI section S3). However, in the IIM, we have a higher run velocity towards the correct decision threshold (fig. 2A), and this makes the correct decision RT shorter than the wrong decisions for a simple asymmetric RnT motion.

On the other hand, the motion in zone II of the IIM is not a simple RnT with instantaneous changes in direction since the spin-flipping events take a finite time to switch their global state. When we analyze a RnT motion with finite time stops during each tumble event, we can obtain a RT that is longer for the correct vs. wrong decisions (see SI section S3). This is due to the fact that trajectories that reach the correct threshold tend to include longer paths with more numerous tumble events that slow down the decision process. Thus, the IIM in the RnT regime (zone II in fig. 2D(ii)) exhibits a RT_c_/RT_w_ that can be both smaller and larger than 1.

The final property of the IIM, which we show in fig. 2E, is the RT distribution in the different regimes, corresponding to the stars indicated in fig. 2C(i). Typical RT distributions are shown in fig. 2E(i) (the time is normalized by the mean RT), and typical trajectories are shown in fig. 2E(ii). We also denote by dashed vertical lines the theoretical ballistic RT as given by 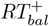 (eq. (8)). The RT distributions of the ordered and intermittent phases show sharp peaks due to the ballistic nature of the trajectories. The differences between the three distributions are quantified using the quantile-quantile plot [30, 31] (fig. 2E(iii)). The distributions for both the ordered and intermittent phases are significantly different from the disordered phase for short times, but the ordered phase (close to the transition) has a long-time tail similar to the disordered phase (see also table S2 for further quantitative measures of these distributions). Overall, our IIM exhibits decision-making properties that are similar to the DDM with different regimes of effective diffusion coefficient as a function of our model parameters (temperature and inhibition). However, the transition from simple diffusion to the ballistic or RnT motion in the ordered phase leads to qualitative deviations from the DDM-like behavior.

## COMPARISON OF THE IIM WITH OTHER DECISION-MAKING MODELS

We now briefly compare the IIM with a few similar or commonly used models for decision making (fig. 3). In the IIM, *n*_1_ and *n*_2_ indicate the instantaneous firing states of the two spin populations that refer to the two neuronal populations (fig. 3A). The spins interact via self-excitation within each group (*J*_*in*_) and cross-inhibition between the groups (*J*_*out*_). The spins tend to fire with higher rates in the presence of bias (*ϵ*_1_, *ϵ*_2_), representing learning based on external information. The global inhibition affects both groups, promoting the “off”-state. *y*_1_ and *y*_2_ represent the integrated quantities, where *y*_1_ = DV = *n*_1_ −*n*_2_*dt* and *y*_2_ = −DV. The decision is made when either *y*_1_ or *y*_2_ reaches the fixed threshold *L*.

**Figure 3:**
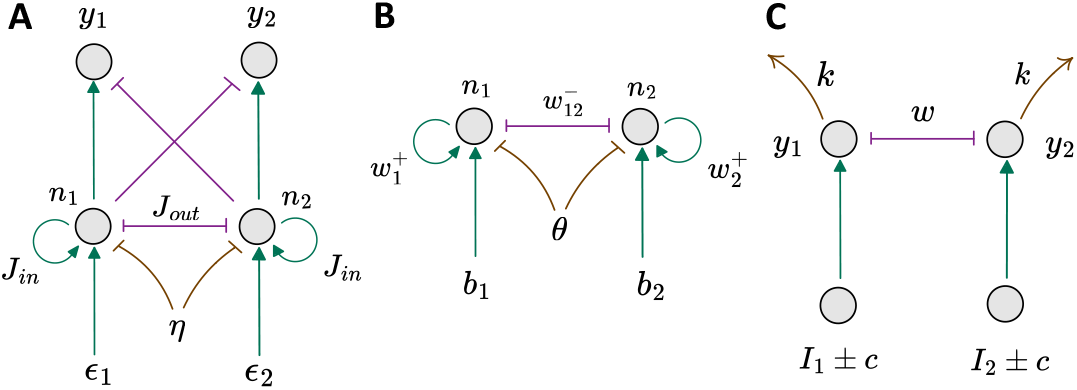
Architectures of different decisionmaking models. A IIM model. B IDM model [9]. C LCA model [2]. The green arrows indicate excitation, the purple lines with flat ends indicate cross-inhibition, and the brown arrows or lines with flat ends indicate global inhibition (IIM), activation threshold (IDM), and leakage (LCA).

The comparison between the IIM and the DDM was mentioned above (see also SI section S3). We note again that in the disordered regime, the IIM recovers the DDM behavior. In this respect, the IIM extends the DDM by introducing correlations in the dynamics arising from the underlying spin interactions. A crucial difference is that in the DDM, the dynamics of the firing that the DV integrates are purely Gaussian white noise (with an additional drift) with no temporal correlations. In contrast, in the IIM, the dynamics have temporal correlations induced by the Ising coupling between the spins. The temporal correlations appear most clearly in the RnT trajectories of the DV for the ordered and intermittent phases (fig. 2A(ii-iii)).

Another theoretical model for decision making that is based on the Ising model in a similar spirit to our work is the Ising Decision Maker (IDM) [9]. In the IDM (fig. 3B), the neural network consists of two pools of neurons (represented by their instantaneous firing states *n*_1_ and *n*_2_) with pairwise excitatory (inside the group, 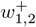) and inhibitory (between the groups, 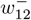) interactions, and activation threshold *θ*. The external fields *b*_1,2_ represent the sensory evidence, and initially all the spins “off”. In the ordered phase of the Ising model, there are two minima that correspond to the states with one of the two spin groups having a large activity, while the other group is inhibited (see SI section S4, fig. S17(a)). The decision in the IDM is made when the system reaches the region around one of these minima, corresponding to one group reaching a high-activity state (SI fig. S17(a)) [32].

The major difference between the IIM and the IDM is that in the IDM, there is no integration of the firing activity of the spins over time (fig. 3A, B). Due to this crucial difference, the IDM can not mimic the DDM model, and in the disordered phase of the Ising model, the IDM becomes locked in indecision (see SI section S4 fig. S17(b)). More details on the comparison between the IIM and IDM in the ordered regime are given in the SI section S4.

Another common class of decision-making models is the Leaky Competing Accumulator (LCA) model [2]. The overall structure of the model is shown in fig. 3C. In this model, there are two accumulator units (*y*_1,2_) that integrate the noisy evidence from two firing neuronal populations (*I*_1,2_ ±*c*). The accumulators interact via cross-inhibition (*w*). The model also allows the decay of the accumulator’s activity (“leakage”, *k*). The decision is made once the activity of one of the integrated quantities reaches a positive threshold *L*. The main difference between the LCA and the IIM is that in the LCA, the cross-inhibition appears only at the level of the integrated firing rates, and there is no cross-inhibition at the level of the underlying firing elements, as in the IIM. The lack of cross-inhibition at the underlying firing signal that enters the integrator means that in the LCA there is no sharp phase transition associated with the decision-making process.

Our model, therefore, contains the phase-transition property of Ising-based models at the neuronal firing level (as in the IDM), while the decision is made at the level of an integrated quantity, similar to the DDM and LCA models. By combining these properties our model extends previous models and exhibits novel dynamical regimes and decision-making properties.

## SPECIAL PROPERTIES OF THE IIM NEAR THE TRICRITICAL POINT

In fig. 2, we demonstrated that the IIM predicts a trade-off between accuracy and speed, which suggests that the region around the phase transition line allows a compromise between these conflicting traits. The system can adjust its parameters with respect to the transition line by varying *T* and/or *η*. Since the temperature (*T*) represents the noise in the neural network it may be less amenable to easy control and adjustment. On the other hand, the global inhibition (*η*) can be readily adjusted by the activity of inhibitory neurons. Motivated by this observation, we explore the role of inhibition as the control parameter that the brain adjusts in order to tune its accuracy, as indicated by recent experiments [10].

In fig. 4A we plot points (blue) in the *T, η* parameter space that have the same accuracy (30% errors) for a given constant (and small) bias (as in fig. 2B(i)). We then consider shifting these points by increasing the inhibition by a small fixed increment *dη* (fig. 4A) and analyze how the error rate and the RT change due to this shift (the green dots in fig. 4A,B). In response to the small increase in inhibition, the error rate decreases (fig. 4B), while the RT increases as expected (fig. 4C). We find that the decrease in error is the most significant for lower temperatures, where the shift in inhibition moves the IIM along the sharp gradient of the error contours (fig. 2B(i)). At higher temperatures, the shift in inhibition has a vanishing effect on the accuracy, as it corresponds to moving the IIM along the error contour. This analysis suggests that the region close to the tricritical point may be advantageous with respect to allowing the brain to gain in accuracy per minimal increase in inhibitory activity. Note, however, that the exact location of the error rate minimum depends on the choice of bias.

**Figure 4:**
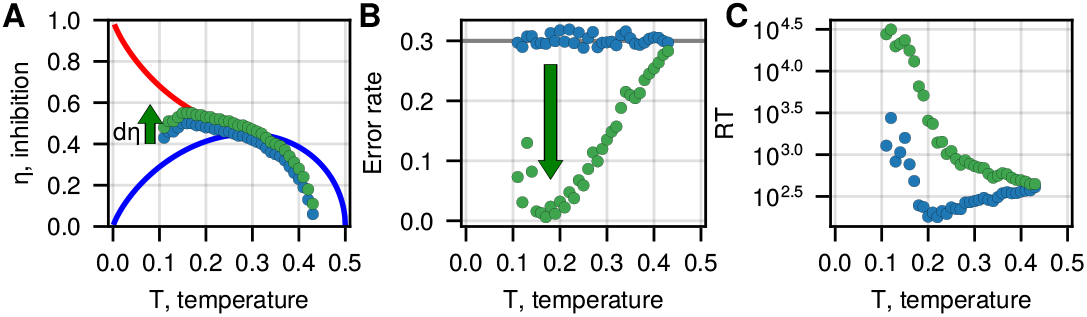
Inhibition controls accuracy. A At fixed bias *ϵ*_1_ = 0.01, we find the points on the phase diagram that have a fixed error level of 0.3 (the blue circles). The green circles denote a shift of the blue circles by increasing the global inhibition by *dη* = 0.05. B The error rate and C the RT are shown for the blue and green points from A. We find that the small increase in inhibition at low temperatures leads to a significant error reduction while the RT drastically increases. At high temperatures, both the error and the RT are less affected by the increase in inhibition.

In the IIM, we can relate the spin states to the neuronal firing activity and see how it depends on the global inhibition. Therefore, we can make some predictions using our model with respect to the neuronal activity during decision making, which is measurable [32, 33]. We show in the SI (section S5) that the learning process in the region near the tricritical point is most “costly”, involving the largest relative increase in bias and neuronal activity (SI fig. S19(a)2, (d)2), but on the other hand, has the advantage of giving the largest relative increase in the decision speed (SI fig. S19(c)2), while the accuracy of the decisions is least sensitive to a constant rate of bias decay (SI fig. S19(c)1).

Note that in the IIM the dynamics, such as the tumble rate in the RnT regime, depend on the system size (number of spins *N*), as shown in SI fig. S4. However, close to the transition line the dependence on *N* diminishes [11], making the behavior close to the transition line insensitive to fluctuations in the number of participating neurons. This is also the region which does not change its dynamics when there are fluctuations in the interaction strength between the spins (SI section S1).

To summarize, we find that near the transition line, close to the tricritical point (in the ordered or intermittent phases), there are special properties of the IIM which may be advantageous for the decision-making process. In this region, the dynamics are most robust to fluctuations in the network size and connectivity and may be an optimal compromise between speed and accuracy, with the ability to most significantly improve accuracy with a small increase in global inhibition. These properties suggest that the decision-making circuit in the brain may correspond in our model to the region in the vicinity of the tricritical point. In the next section, we analyze new experimental data, which we compare to the IIM, and find support for this hypothesis.

Experimental results, calculated for 20 volunteers, who participated in a game of 60 gain and 60 loss trials of two-choice tasks under uncertainty, where the symbols in each pair encode a monetary gain or loss with 70% and 30% probabilities. The error rate indicates the proportion of the wrong choices in trials 34-60, after the learning period. RT_gain_ and RT_loss_ are the mean reaction times in the gain and loss trials after the learning period, and RT_c_ and RT_w_ are the mean RTs in the correct and wrong decisions. Note that the ratios are calculated per participant and then averaged.

## COMPARING THE IIM TO EXPERIMENTS

To test the IIM, we analyze recent experimental data obtained from volunteers playing a two-armed bandit game [34]. In all the games, the subject chooses one option (in the form of a special character) per trial, and this character either gives monetary gain (0 or +1) or loss (0 or −1) as a reward, with some fixed probabilities (which are unknown to the subject). The goal is to maximize the total score (fig. 5A). Each pair of symbols refers to either gain or loss trials. We define the correct option as the option that increases the total score with a higher probability in the gain condition and decreases the total score with a lower probability in the loss condition. The gain and loss trials can be either separated, so the participants learn the hidden probabilities of the rewards for one pair in a game, or alternatively, the gain and loss trials can be intermixed, and the two pairs alternate randomly in the same game. Since the probabilities encoded by the symbols are unknown to the participants, the initial trials give rise to a learning process during which the participants form a bias towards one of the options in each pair. During these experiments, the choices of the participants and the corresponding decision time (reaction time, RT) were registered.

**Figure 5:**
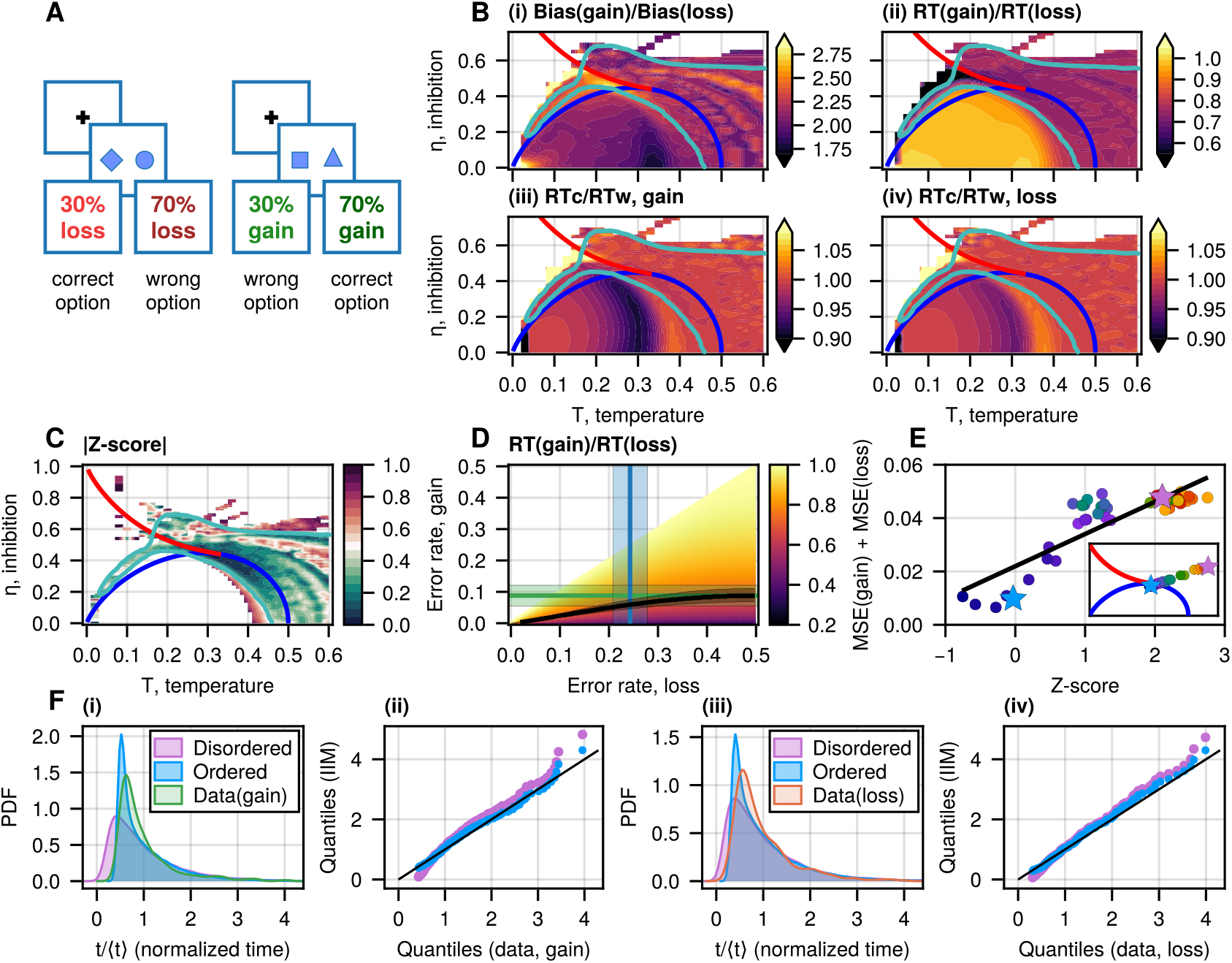
Experimental setup I compared to the IIM. A Task design in experimental setup I. The gain and loss trials are intermixed. Each symbol encodes a monetary gain (0 or 1) under the gain conditions or a monetary loss (0 or −1) under the loss conditions, with fixed probabilities 70% and 30%. The participants learn these differences during 120 trials. B Heatmaps of various quantities. The red and blue lines on the heatmap denote the first and second-order transitions, respectively. (i) The ratio of the biases that fit the measured errors in the gain and loss conditions (table I) for each point of the phase space. B(ii) The RT ratio in the gain and loss conditions calculated using the biases obtained to give the observed error rates. B(iii-iv) The ratio RT_c_/RT_w_ for the gain and loss conditions. C Phase diagram with a heatmap denoting the absolute value of the Z-score, which measures the deviation between the calculated and experimentally observed ratio RT_gain_/RT_loss_ (see main text). The turquoise contour is a guide to the eye, indicating the region of the heatmap with |Z|-score ≤0.3, where agreement is high. D Heatmap of the analytical ratio of the mean RTs in the gain and loss conditions for the DDM model as a function of the errors in the two conditions (eq. (11)). The green and blue lines and the shaded rectangles indicate the error rates in the gain and loss conditions from the experiment (table I). The black line and the shaded area indicate the mean RT ratio in the DDM that satisfies the observed ratio RT_gain_/RT_loss_ of 0.7± 0.04. E Quantification of the deviations between the simulated and experimental RT distributions (normalized by the mean RT) by the mean squared error (MSE) summed over the distributions, adding up both the gain and loss cases. The MSE is plotted as a function of the Z-score (C). The points are taken along a line shown in the inset. The region with the lowest Z-score corresponds to the lowest MSE. F(i,iii) Examples of RT distributions for two values in the RnT and diffusion regimes (marked by stars of the corresponding colors in E) for both gain and loss conditions, compared with the experimental data. F(ii,iv) A quantile-quantile plot for this comparison of the RT distributions.

### Setup I: intermixed gain and loss trials

In this version of the experiment, 20 volunteers played a game of 60 gain and 60 loss trials, which were intermixed randomly. In each trial, a pair of symbols represented a probability of monetary gain (with 70% and 30% probability, respectively) or monetary loss (70% and 30%), fig. 5A (see SI section S6 for the detailed explanation and analysis). The experiment was approved by the Weizmann Institutional Review Board.

The results are shown in table I. The error rate indicates the proportion of wrong choices in trials 34-90, where the errors seem to be saturated following the initial learning period (see SI section S6). We give the ratio between the mean reaction times RT_gain_ and RT_loss_ in the gain and loss trials, respectively, and the ratio between the mean reaction times for the correct and wrong decisions (RT_c_ and RT_w_) for the gain and loss trials separately. Note that the ratios are calculated per participant and then averaged. The mean ratio of the RTs for the correct and wrong decisions (RT_c_/RT_w_) is very close to but slightly smaller than 1 (t-test, gain: *p*_8_ = 0.11; loss: *p*_15_ = 0.25), which might indicate slow errors [8, 9].

**Table I:** Experimental setup I: intermixed gain/loss trials.

| Parameter | mean ± SE |
| --- | --- |
| Error rate, gain | $0.09 \pm 0.03$ |
| Error rate, loss | $0.24 \pm 0.04$ |
| $RT_{\text{gain}}/RT_{\text{loss}}$ | $0.70 \pm 0.04$ |
| $RT_c/RT_w$, gain | $0.82 \pm 0.10$ |
| $RT_c/RT_w$, loss | $0.93 \pm 0.06$ |

The most outstanding feature that we find in this experiment is the significant difference between the error rates in the gain and loss trials. This starkly differs from the behavior observed in the next experiment, where the gain and loss trials were conducted in separate games (see the following subsection). This observation is surprising since the reward probabilities of the two options are the same in both gain and loss trials. We also observe a large difference in the mean RTs between the gain and loss trials, with the gain decisions occurring significantly faster.

Within our model, bias is the only parameter which, when increased, decreases both the error rate and the RT simultaneously (see SI section S2, section S7), in agreement with the differences between the gain and loss trials in the experiment (table I). In addition, since both gain and loss trials interchange during the game and have the same reward probability, we assume that *T* and *η* are fixed during the game, and we use bias as the only free parameter that changes between the gain and loss cases, which means that the participants have formed different biases for the gain and loss cases during the learning period. We identify two biases *ϵ*_1,gain_, *ϵ*_1,loss_ per each point (*T, η*), such that it gives the error rate of both the gain or loss conditions (as given in table I). As expected, the bias for the gain conditions is larger, leading to lower error (fig. 5B(i)).

We then calculate the RT for the two cases and the corresponding ratio RT_gain_/RT_loss_ (fig. 5B(ii)). In fig. 5C, we show the area on the phase diagram where the ratio RT_gain_/RT_loss_ (as represented by the different biases in the model) is in agreement with the experimental observation within the error bars (table I). Within this area, we indicate by green line a region where the agreement with the experiment is strongest, as defined by having low| *Z* |-score: |RT_gain_/RT_loss_ −*µ*| */σ* (here *µ* = 0.7, *σ* = 0.04 are taken from table I). This region was chosen such that the ratio |RT_gain_/RT_loss_ predicted by the IIM is within 0.3*σ* from the mean observed value *µ*. While the entire disordered phase satisfies the experimental observation (within the error bars), the most optimal region in our model that fits the experimental data lies near the tricritical point (see more details in SI section S8, fig. S24, S25).

We now demonstrate that the regular DDM, equivalent to the disordered phase of the IIM, gives only a marginal fit to the experimental data. We calculate the analytical expression for the ratio of the RTs at the given error rates in terms of the dimensionless Péclet number Pe = *vL/D*, which characterizes the ratio between the diffusion and drift
time scales

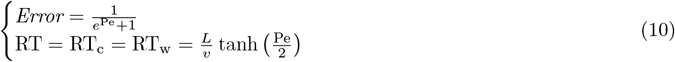

where the *Error* is the probability of reaching the negative decision threshold −*L, v* is a constant drift, and *D* is the diffusion coefficient. As we only varied the bias in the IIM, we keep the drift velocity *v* as a control parameter and fix the diffusion coefficient *D* and the threshold *L*. Then, we express the ratio RT_gain_/RT_loss_ as a function of the error rates (*Error*_gain_, *Error*_loss_):

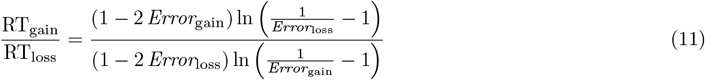

Plugging the experimental values from table I into the analytical solution of the DDM (eq. (11)), we find that the DDM’s RT ratio is: RT_gain_/RT_loss_ = 0.78 ± 0.11. This is in good agreement with the average numerical result in the disordered region of the IIM: RT_gain_/RT_loss_ = 0.76 ± 0.05 (fig. 5B(ii), see also SI section S8, fig. S24). We can, therefore, use the analytical calculation for the DDM to demonstrate that it only gives a marginal fit to the experimental data of the RT ratio, fig. 5D (hypothesis: RT_gain_/RT_loss_(data) ≠ 0.78; t-test: *df* = 15, *t* = −1.676, *p* = 0.1144; Wilcoxon-test: *W* = 39, *p* = 0.1439). This analysis shows that while our IIM near the tricritical point (the RnT regime) fits the experimental data very well, it can also be marginally fitted by the disordered phase, which corresponds to the DDM behavior (SI, fig. S24). By comparing to an analytical RnT model for the DV dynamics, we show that this is the crucial property that allows the IIM to fit the experimental data so well near the tricritical point (see the analysis in SI, fig. S25).

We can also compare the experimental ratios of correct vs. wrong decision RT (table I) to the theoretical expectation. In the DDM, this ratio is strictly 1, as in the disordered phase of the IIM. The experimental data gives ratios that are smaller than 1 on average, but these differences are within the error bars. We note that in the best-fit region of the IIM (fig. 5B(iii-iv)), just below the transition line, there are small deviations of this ratio from 1.

Finally, we compare the RT distributions (as a function of the normalized time divided by the mean RT) extracted from the experiment and those predicted by the model. We limit this comparison to a line within the region of the theoretical parameter space that fits the mean RT ratio of the gain and loss experiments (shown in the inset of fig. 5E). We chose this line to span the behavior from RnT to pure diffusion and cover the range of Z-score values (see also fig. 5C). We quantify the deviations between the simulated and experimental RT distributions by the mean squared error (MSE) summed over the RT distributions. We show in fig. 5E that the region near the transition line, with the lowest Z-score, also gives the best fit for the shape of the RT distribution. This is demonstrated for two values in the RnT and diffusion regimes in fig. 5F, for both gain and loss conditions. This result further supports our conclusion that the RnT dynamics near the transition line provide a more accurate description of the experimental data compared to the purely DDM description.

### A. Setup II: separate gain and loss trials

In the second set of experiments [35], the volunteers played separate games of gain and loss. The main novelty was the ability to monitor the concentration of inhibitory neurotransmitter (*γ*-aminobutyric-acid, GABA) during the decision-making process. The GABA concentration was quantified from the dorsal anterior cingulate cortex (dACC), using Proton Magnetic Resonance Spectroscopy (^1^H-MRS) at 7T (see more details in SI section S9 and [35]). In the experiment, 107 volunteers played four separate games (of 50 trials), in each of the following combinations: gain and loss with probabilities of 65/35 and 50/50.

Under the unbiased conditions, when the probabilities of zero and non-zero rewards for both options were 50% (no correct option), the RT was measured as the baseline (labeled RT_50/50_). Following the initial learning period (28 trials), trials 29-50 are used for the data analysis (see the details in SI section S9). The gain and loss trials did not show significant differences in their error rates and RT, so their data was combined (SI section S9).

For a further analysis, we divided the participants into three groups according to their error rates and RT normalized by the unbiased RT (i.e., RT_65/35_/RT_50/50_, fig. 6A(i)). The orange group indicates the volunteers who did not learn the correct choice very well and had an error rate larger than 0.2 (an arbitrary threshold, but the analysis is insensitive to the value of this threshold as we demonstrate in the SI section S9 C).

**Figure 6:**
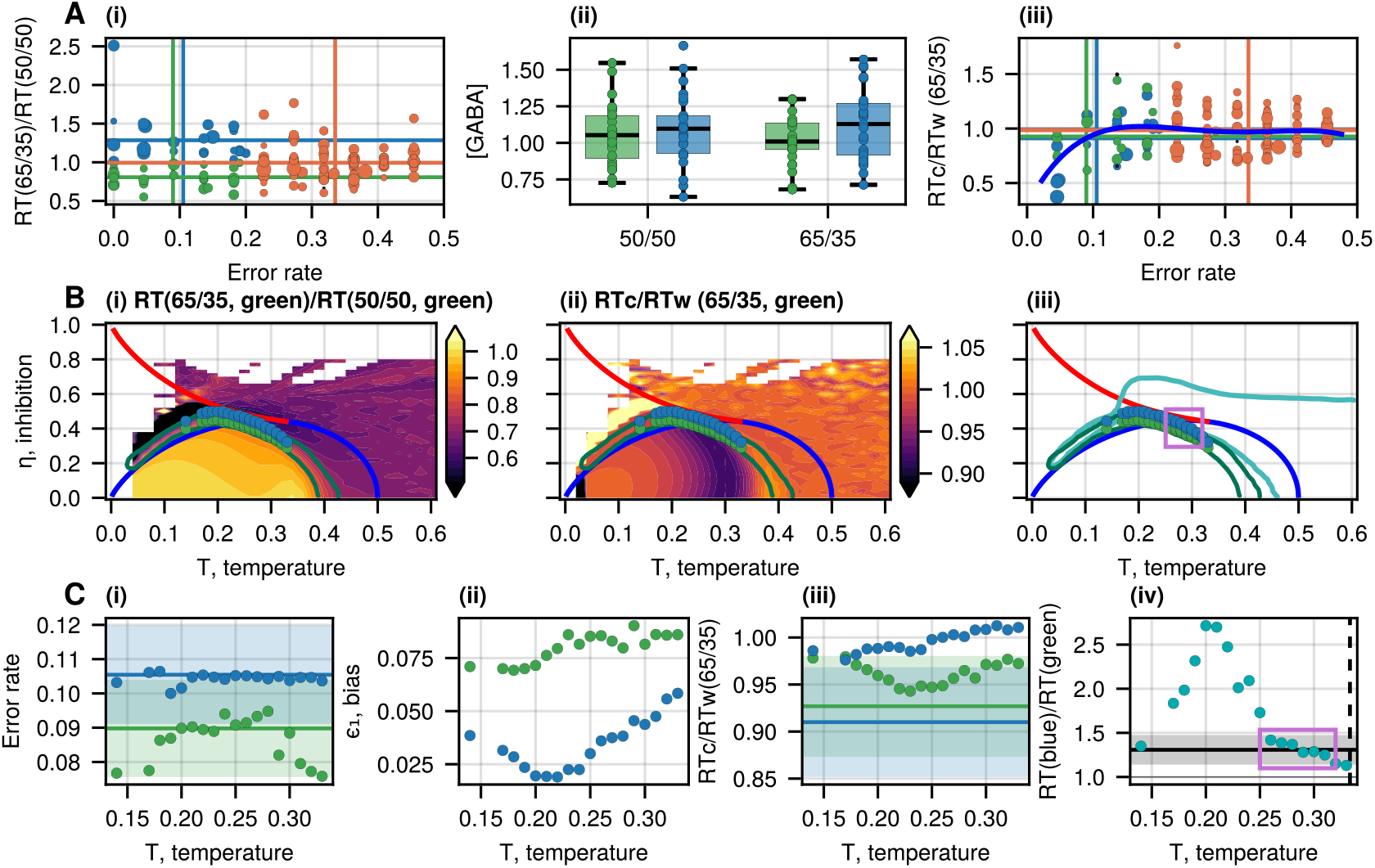
Experimental Setup II compared to the IIM. A(i) Normalized RT (RT_65/35_/RT_50/50_) as a function of the error rate. Each RT for the gain or loss trials (with the probabilities of 65%-35% per choice) is normalized by the participant’s RT in the unbiased condition (with the reward probability of 50% per choice). Each point represents a result from a single participant, for gain and loss separately. The size of the points is related to the average GABA concentration measured for each participant during the task. The data is divided into three groups: the green group has error rate ≤ 0.2 and normalized RT ≤1, the blue group has error rate ≤ 0.2 and RT *>* 1, while the orange group has error rate *>* 0.2. The vertical and horizontal lines denote the average error rate and normalized RT for each group. A(ii) The GABA concentration, quantified from the dorsal anterior cingulate cortex (dACC) during the task for the unbiased and biased trials for the green and blue groups. We find that the concentration of GABA_50/50_ (green) is not different from GABA_50/50_ (blue) (unequal variance t-test: p = 0.591), though for the biased conditions, the concentration GABA_65/35/_ (green) shows a marginal difference with GABA_65/35_ (blue) (unequal variance t-test: p = 0.0725, see also SI section S9, table S9). A(iii) The ratio RT_c_/RT_w_ for the same groups of A(i) at the biased conditions (both gain and loss). Each point represents a result from a single volunteer. The size of the points is related to the average GABA concentration of each participant during the task. The blue line indicates a 4th-degree polynomial fitted to the data as a guide to the eye. B(i) The normalized RT ratio (between the biased and unbiased conditions) for the average error rate of the green group (table II), as given by the IIM. For each point of the phase diagram, we find the bias that satisfies the error rate of the green group (0.09 ± 0.01), and this gives us the biased RT_65/35_. The dark green contour denotes the region that fits the green group’s normalized RT ratio (RT_65/35_/RT_50/50_, without constraining the ratio RT_c_/RT_w_). The green circles correspond to values of the parameters that also fit the green group’s RT_c_/RT_w_ ratio. The blue circles denote a shift of the green circles by increasing the global inhibition by a factor of 1.17, which is the ratio of the measured average GABA concentrations in the two groups for the biased conditions (fig. 6A(ii)). The red and blue lines on the heatmaps denote the first and second-order transitions, respectively. B(ii) Heatmap of the RT_c_/RT_w_ given by the IIM for the average error rate of the green group under the biased conditions. B(iii) Comparison of the phase space regions that fit the experimental data in setups I and II. The turquoise contour indicates the area of the phase space that best matched the experimental data of setup I (fig. 5C). C(i-ii) The error rate and biases of the green and blue circles from B, as a function of temperature *T*. The calculated error rates agree with the mean values of the experimental observations (denoted by the horizontal lines and shading). The higher error rate for the blue circles corresponds to lower biases. C(iii) The ratio RT_c_/RT_w_ for the green and blue circles in B. The green circles agree well with the experimental observation (denoted by the horizontal lines and shading), while the blue circles indicate a lower agreement. C(iv) The ratio of the mean RTs for the green and blue circles in B as a function of temperature *T*. The black line and the shaded area indicate the ratio of the average biased RTs between the blue and green groups in the experiment: 1.31 ±0.17. The purple box indicates the region of best agreement, near the tricritical point (black vertical line denotes *T*_tri_).

The error rate of the green group is below the 0.2 threshold, and the ratio RT_65/35_/RT_50/50_ ≤1. This group of participants makes accurate and fast decisions, as expected in our model since a strong bias gives rise to accurate decisions and decreases the RT (fig. S7). This general property also appears in the DDM.

Surprisingly, we find another group of participants (blue group, fig. 6A(i)) that make accurate decisions (error rate ≤ 0.2), but slower than in the unbiased case: the ratio RT_65/35_/RT_50/50_ *>* 1. The low error rate indicates a significant bias, and it is, therefore, surprising that the bias does not manifest in faster decisions. An important observation that may explain this puzzling group of volunteers is given in fig. 6A(ii). For the green and blue groups, we compare the GABA concentration during the tasks in the biased and unbiased conditions. We find that the GABA concentrations in both groups are not different in the unbiased case, while in the biased conditions, the blue group exhibits larger concentrations compared to the green group.

Finally, we plot the ratio of correct vs. wrong RTs for the three groups (RT_c_/RT_w_, fig. 6A(iii)). The experimental data shows a significant decrease in this ratio for the green and blue groups (at low error rate values), which indicates a deviation from the DDM prediction (where this ratio is strictly 1).

We now systematically compare all of the experimental data described above and summarized in table II to our IIM. We start by fitting the data of the green group (see SI section S7). For each point (*T, η*), we find the bias that corresponds to the measured average error rate of 0.09± 0.01. Using this bias, we derive the normalized RT (RT_65/35_/RT_50/50_, fig. 6B(i)) and the ratio RT_c_/RT_w_ (fig. 6B(ii)). Then, we first select the region that fits the observed normalized RT of 0.81 ± 0.02, denoted by the dark-green contour. Next, we also fit to the observed ratio RT_c_/RT_w_ = 0.93 ± 0.05 and find a narrow region near the tricritical point, denoted by the green points in fig. 6B. It is satisfying to find that this narrow region of parameters lies close to the edge of the region that fits the experiments of the previous section (fig. 6B(iii)).

**Table II:** Experimental setup II: separated gain and loss trials.

| mean ± SE | Green | Blue | Orange |
| --- | --- | --- | --- |
| Size | 27 | 26 | 91 |
| Error rate | $0.09 \pm 0.01$ | $0.11 \pm 0.01$ | $0.34 \pm 0.01$ |
| $RT_{65/35}$ (sec) | $0.61 \pm 0.04$ | $0.79 \pm 0.05$ | $0.67 \pm 0.02$ |
| $RT_{65/35}/RT_{50/50}$ | $0.81 \pm 0.02$ | $1.28 \pm 0.06$ | $0.99 \pm 0.02$ |
| $RT_c/RT_w$, 65/35 | $0.93 \pm 0.05$ | $0.91 \pm 0.06$ | $0.98 \pm 0.02$ |
| GABA<sub>65/35</sub> | $1.02 \pm 0.03$ | $1.13 \pm 0.05$ | $1.03 \pm 0.02$ |
| GABA<sub>50/50</sub> | $1.06 \pm 0.04$ | $1.09 \pm 0.05$ | $1.03 \pm 0.02$ |

Next, we wish to explain the behavior of the blue group using our model. Guided by the observation of larger inhibitory signals for these volunteers compared to the green group (fig. 6A(ii)), we assume a simple linear relation between the GABA concentration and the level of global inhibition in the IIM. We, therefore, use the measured ratio GABA_65/35_(blue)/GABA_65/35_(green) = 1.17 *±* 0.11 (table II) and simply shift with respect to the green group to larger values of *η* (the blue points in fig. 6B). Note that an increase in the cross-inhibition between the spin groups, which will also manifest in an increase in the GABA concentration, has the opposite effect of lowering the accuracy and decreasing the RT (see SI section S2, fig. S8).

Experimental results, calculated for 107 volunteers participated in four games of 50 trials of two-choice tasks under uncertainty, where in two games, the symbols in each pair encode a monetary gain or loss with 65% and 35% probabilities, and in the other two games, the symbols give a reward with equal probabilities 50%. The results of the gain and loss trials are combined, as they do not show any significant differences (SI section S9). The group names relate to the color in fig. 6A(i). The error rate indicates the proportion of the wrong choices in trials 29-50, after the learning period. The mean RTs during the trials after the learning period, and the RT_c_ and RT_w_ are the mean RTs in the correct and wrong decisions. The GABA concentration is quantified from the dorsal anterior cingulate cortex (dACC) during the task for the biased and unbiased conditions.

We now wish to test if our interpretation of the blue group as having higher global inhibition is consistent with the observed increase of the RT_65/35_ between the green and blue groups (table II). We start by finding the values of the bias in the IIM for the blue points (fig. 6C(ii)) which gives us the observed error rate of 0.11± 0.01 (fig. 6C(i)). Using these values, we extract the ratio RT_c_/RT_w_ (fig. 6C(iii)), and we see that the blue points are slightly outside the experimental data. We next calculate the RTs (RT_65/35_) and the ratio of the biased RT_65/35_(blue)*/* RT_65/35_(green) (fig. 6C(iv)). We find that the RT ratio calculated from our IIM agrees very well with the observed value (black line and shading) at temperatures close to the tricritical point (fig. 6B(iii)).

The comparison between the experimental data and the analytic relation between error and normalized RT for the DDM model shows that this model does not agree with the data (SI section S9 E).

The good agreement between the IIM and the data suggests the following interpretation of the differences between the two groups of volunteers. While the green group learns the correct choice with a strong bias (therefore having fast and accurate decisions), the blue group does not learn so well (weaker bias, fig. 6C(ii)) but instead compensates with an increase in inhibition (exhibiting higher GABA concentrations) to improve the accuracy of the decisions. The price that the blue group pays is slower decisions compared to the green group. The self-consistency and robustness of our analysis and agreement with the IIM are demonstrated using different threshold values to divide the data into the three groups (section S9 C).

## DISCUSSION

We have presented here a new theoretical framework for describing the decision-making process in the brain, the Integrated Ising Model (IIM). It is based on Ising spins whose state represents the firing of neurons, arranged in groups that represent each one of the available options, and interact in an excitatory manner within the group, while cross-inhibiting the spins in the other group. The states of these spins drive the state of an integrator, which acts as the Decision Variable (DV), and upon reaching one of two threshold values, a decision is made. This last property is identical to the highly successful drift-diffusion model (DDM) [5], which the IIM recovers in its disordered regime (high levels of noise and inhibition).

The IIM model goes beyond the DDM, and in its ordered phase it displays run-and-tumble dynamics (RnT) with significant deviations from the DDM. In this phase, we find faster and less accurate decisions with a speed-accuracy trade-off, which may be optimized near the 2nd-order phase transition line. Just below this transition line, near the tricritical point, the model predicts maximal gain in accuracy as a function of an increase in global inhibition, suggesting a mechanism for explaining the observed increased inhibition when uncertainty is high [10]. In the same region below the transition line, we find a minimum in the rate of accuracy decay per loss of learned bias, which is an advantage for maintaining accuracy for a longer time. This region is also where the behavior is insensitive to fluctuations in the size of the spin groups and the strength of their interactions. Therefore, we find that the IIM suggests that it can be advantageous for the brain to be in this critical region.

We compare our IIM to two new experimental data sets, which support the model’s prediction regarding the importance of the critical regime. The first experiment allows us to map the data to a region of the IIM phase space (*T, η*) around the tricritical point and the 2nd-order transition line. The second experiment also contained measurements about changes in the inhibition strength within the decision-making region of the brain. This data set again localizes the data to an area of the IIM phase space that is in the vicinity of the tricritical point. Notably, both experimental data sets do not fit well within the regular DDM. Both experimental results therefore suggest that the brain utilizes the special properties of the critical region near the phase transition line. This result is different from the criticality that was proposed to exist in the brain with respect to the structure and the spatial connectivity of the neural network [36–39], including the effects of global inhibition [40], or in collective animal systems [41]. It is another form of criticality that is not manifested in the spatial interactions (which are all-to-all in our model) but rather in the dynamics of the decision-making process.

Using our model, we interpret the experimental data as indicative of two ways that the brain can achieve accurate decisions. The first is based on developing a strong bias to the correct choice, leading to fast decisions, while the second is based on weaker bias, compensated by higher inhibition that leads to slow and accurate decisions. These two processes may be reminiscent of the fast-and-slow decision-making processes described by Daniel Kahneman’s “Thinking, Fast and Slow” [42].

Note that our spin-based model for decision making was motivated by the success of this approach in describing the decision making of individual animals and animal groups while navigating through space [12–14]. More theoretical work is planned to further elucidate the properties of the IIM, such as extending it to describe more than binary choices [2, 30, 43, 44]. In the future, we intend to explore the IIM in the context of perception tasks as well as in connection with spatial navigation. In addition, future experimental work is needed to further test the proposal made in this work regarding the criticality of the decision-making process.

## ACKNOWLEDGMENTS

We acknowledge financial support from the Israeli Science Foundation (ISF) personal grant 416/20 and the National Institutes of Health (NIH) grant R01-AG080672 to Assaf Tal. This work was partially supported by Nella and Leon Benoziyo Center for Neurosciences (Weizmann Institute).

## Notes

### Competing Interest Statement

The authors have declared no competing interest.

