## Supplementary Information for "Integrated Ising Model with global inhibition for decision making"


### S1. IIM PHASE DIAGRAM: ASYMMETRIC INTERACTIONS

Denote:
$k = 2$ – number of abstract options
$N$ – total number of spins in both groups.
$N_{\text{on}}^{I,II}$ – number of active spins in groups I, II.
$n_{1,2} = N_{\text{on}}^{I,II}/N$ – fraction of active spins in groups I, II.
$\vec{p_{1,2}}$ – vectors from the abstract integrator toward two abstract targets in the two-dimensional abstract space
$\vec{V} = n_1\vec{p}_1 + n_2\vec{p}_2$ – solution of the MF master equation
$\vec{V}_0 = \frac{1}{2}V_0(\vec{p}_1 + \vec{p}_2)$ – compromise solution

In this section, we explore the compromise and non-compromise solutions for the fractions of active spins and their stability in the binary case ($k = 2$ is the number of options), where the cross-inhibition $\text{J}_{\text{out}}$ between two competing groups of spins is lower than the self-excitation $\text{J}_{\text{in}}$ within each group. Assuming constant $\text{J}_{\text{in}}$ and $\text{J}_{\text{out}}$, we derive the dynamical equations:

$$\begin{cases} \dfrac{dn_1}{dt} = \dfrac{1/k}{1+\exp\left(\frac{-k(\text{J}_{\text{in}}\,n_1+\text{J}_{\text{out}}\,n_2)+\eta-\epsilon_1}{T}\right)} - n_1 \\ \dfrac{dn_2}{dt} = \dfrac{1/k}{1+\exp\left(\frac{-k(\text{J}_{\text{in}}\,n_2+\text{J}_{\text{out}}\,n_1)+\eta-\epsilon_2}{T}\right)} - n_2 \end{cases} \tag{S1}$$

*a. Introduce V*

Previously, in the symmetric case ($\text{J}_{\text{out}} = -\text{J}_{\text{in}} = -1$), we assumed that an abstract agent integrates the firing activity of the spins in the system, and its coordinate DV moves along a one-dimensional line until it reaches either positive or negative threshold $L$. We now place the integrator and the abstract targets in a two-dimensional space in the following way. The vectors $\vec{p}_i$, directed from the agent to the abstract targets, are defined according to the values of the coupling constants so that

$$\begin{cases} \vec{p}_1\vec{p}_1 = \vec{p}_2\vec{p}_2 = \text{J}_{\text{in}} = 1 \\ \vec{p}_1\vec{p}_2 = \text{J}_{\text{out}} < 0 \end{cases} \tag{S2}$$

We introduce a new parameter $\vec{V}$, which refers to the agent's velocity towards the abstract targets

$$\vec{V} = \sum_i n_i\vec{p}_i = n_1\vec{p}_1 + n_2\vec{p}_2 \tag{S3}$$

where $n_i$ is the fraction of active spins in group $i$. This approach resembles the modeling of the animal motion towards two stationary targets in the real space [1]. Then, the projections of $\vec{V}$ on each direction $p_i$ is

$$\begin{cases} \vec{V}\vec{p}_1 = \text{J}_{\text{in}}\,n_1 + \text{J}_{\text{out}}\,n_2 \\ \vec{V}\vec{p}_2 = \text{J}_{\text{in}}\,n_2 + \text{J}_{\text{out}}\,n_1 \end{cases} \tag{S4}$$

As a result, we can re-write the dynamical equations for the fractions of active spins as follows:

$$\begin{cases} \dfrac{dn_1}{dt} = \dfrac{1/k}{1+\exp\left(\frac{-k\vec{V}\vec{p}_1+\eta-\epsilon_1}{T}\right)} - n_1 \\ \dfrac{dn_2}{dt} = \dfrac{1/k}{1+\exp\left(\frac{-k\vec{V}\vec{p}_2+\eta-\epsilon_2}{T}\right)} - n_2 \end{cases} \tag{S5}$$

Then, we obtain a single closed master equation for the system's dynamics with respect to $\vec{V}$:

$$\frac{d\vec{V}}{dt} = \sum_i \frac{1/k}{1+\exp\left(\frac{-k\vec{V}\vec{p}_i+\eta-\epsilon_i}{T}\right)}\vec{p}_i - \vec{V} \tag{S6}$$

*b. Compromise solutions*

We define the compromise solution as a solution with equal activity in the two groups: $n_1 = n_2$. Therefore, the velocity is $\vec{V_0} = V_0(\vec{p}_1+\vec{p}_2)/k$, where $V_0$ is the amplitude of the compromise solution which describes the total activity in the system. Then, we get the fractions of the active spins and the projection of $\vec{V_0}$

$$\begin{cases} n_1 = n_2 = V_0/k \\ \vec{V_0}\vec{p}_1 = \vec{V_0}\vec{p}_2 = V_0(\mathrm{J_{in}}+\mathrm{J_{out}})/k \end{cases} \tag{S7}$$

Here $V_0 \le 1$ because $n_1, n_2 \le 1/k$. In the unbiased case $\epsilon_i = 0$, we get the following master equation for the compromise solution at the steady state:

$$F(V_0) = \frac{dV_0}{dt} = \frac{1}{1+\exp\left(\frac{-V_0(\mathrm{J_{in}}+\mathrm{J_{out}})+\eta}{T}\right)} - V_0 = 0 \tag{S8}$$

In the limit of high temperatures $T \to \infty$, there is only one solution $V_0 = 1/2$. At low temperatures, two solutions can co-exist: the high-activity state $V_0 > 1/2$ and the low-activity state $V_0 < 1/2$. The compromise solutions appear (or disappear) once $F(V_0)$ crosses 0 at its peak, so we can find the area where one of two compromise solutions vanishes by solving $dF(V_0)/dV_0 = 0$:

$$\frac{dF(V_0)}{dV_0} = -1 + \frac{(\mathrm{J_{in}}+\mathrm{J_{out}})\exp\left(\frac{V_0(\mathrm{J_{in}}+\mathrm{J_{out}})+\eta}{T}\right)}{T\left[\exp\left(\frac{V_0(\mathrm{J_{in}}+\mathrm{J_{out}})}{T}\right)+\exp\left(\frac{\eta}{T}\right)\right]^2} = 0 \tag{S9}$$

In the limit $\mathrm{J_{in}} = 1$, $\mathrm{J_{out}} \to -1$, we get $\frac{dF(V_0)}{dV_0} = -1$, meaning that only one compromise solution exists.

*c. Stability of the compromise solutions*

For each compromise solution, we aim to check its stability. For this, we perturb $\vec{V_0}$ along the perpendicular direction $\vec{l} = \vec{p}_1 - \vec{p}_2$:

$$\begin{cases} \vec{V} = \vec{V_0} + \vec{\epsilon} \\ \vec{\epsilon} = \epsilon(\vec{p}_1 - \vec{p}_2)/k \\ (\vec{p}_1+\vec{p}_2)(\vec{p}_1-\vec{p}_2) = 0 \end{cases} \tag{S10}$$

We plug $\vec{V}$ into eq. (S6) and expand it to the 1st order in $\epsilon$:

$$\frac{d\vec{\epsilon}}{dt} = \sum_i \frac{(\vec{p}_1-\vec{p}_2)\cdot\vec{p}_i}{4T\left[1+\cosh\left(\frac{-k\vec{V_0}\vec{p}_i+\eta}{T}\right)\right]}\epsilon\vec{p}_i - \vec{\epsilon} \tag{S11}$$

Then, we project it on $\vec{p}_1 - \vec{p}_2$:

$$\frac{d\epsilon}{dt}(\mathrm{J_{in}}-\mathrm{J_{out}}) = \sum_i \frac{\left[(\vec{p}_1-\vec{p}_2)\cdot\vec{p}_i\right]^2}{4T\left[1+\cosh\left(\frac{-k\vec{V_0}\vec{p}_i+\eta}{T}\right)\right]}\epsilon - \epsilon(\mathrm{J_{in}}-\mathrm{J_{out}}) \tag{S12}$$

where $\left[(\vec{p}_1-\vec{p}_2)\cdot\vec{p}_i\right]^2 = (\mathrm{J_{in}}-\mathrm{J_{out}})^2$.

Assuming that $|\,\mathrm{J_{out}}\,| < |\,\mathrm{J_{in}}\,|$, we get the stability equation, which we solve together with eq. (S8) to find the second-order transition line:

$$\frac{d\epsilon}{dt} = \epsilon\left(-1 + \frac{\mathrm{J_{in}}-\mathrm{J_{out}}}{2T\left[1+\cosh\left(\frac{-V_0(\mathrm{J_{in}}+\mathrm{J_{out}})+\eta}{T}\right)\right]}\right) \tag{S13}$$

In the limit $\mathrm{J_{in}} = 1$, $\mathrm{J_{out}} \to -1$, we get

$$\frac{d\epsilon}{dt} = \epsilon\left(-1+\frac{1}{T\left[1+\cosh\left(\frac{\eta}{T}\right)\right]}\right) = 0 \Rightarrow \eta = T\,\mathrm{arccosh}\left(\frac{1-T}{T}\right) \tag{S14}$$

which transforms into the second-order transition line in the symmetric case.

The constraint on the right edge of the second-order transition appears where the compromise solutions exist irrespective of inhibition. At $T > T_{\mathrm{crit}}$, the peak of $d\epsilon/\epsilon dt$ stops crossing 0 and is always negative. Plugging one possible peak of $d\epsilon/\epsilon dt$ at $V_0 = 0, \eta = 0$ into $d\epsilon/\epsilon dt \ge 0$, we obtain

$$\left.\frac{d\epsilon}{\epsilon dt}\right|_{V_0=0,\ \eta=0} \ge 0 \Rightarrow T_{\mathrm{crit}} = \frac{\mathrm{J_{in}}-\mathrm{J_{out}}}{4} \tag{S15}$$

We can also find inhibition $\eta$, at which $V_0 = 1/2$ ($n_0 = 1/4$) is the solution at $T_{\mathrm{crit}}$ (the right edge of the ordered phase):

$$\eta_{\mathrm{crit}} = \frac{\mathrm{J_{in}}+\mathrm{J_{out}}}{2} \tag{S16}$$

*d. Non-compromise solutions*

We introduce the difference in the activity of the two groups $v = n_1 - n_2$, which can take zero and non-zero values:

$$f(v) = \frac{dv}{dt} = -v + \frac{1/k}{1 + \exp\left(\frac{-k(\mathrm{J_{in}}\, n_1 + \mathrm{J_{out}}\, n_2) + \eta}{T}\right)} - \frac{1/k}{1 + \exp\left(\frac{-k(\mathrm{J_{in}}\, n_2 + \mathrm{J_{out}}\, n_1) + \eta}{T}\right)} \tag{S17}$$

We replace $n_1 = v + n_2$ to re-write the following expressions:

$$\begin{cases} \mathrm{J_{in}}\, n_1 + \mathrm{J_{out}}\, n_2 = \mathrm{J_{in}}\, n_1 - \mathrm{J_{in}}\, n_2 + \mathrm{J_{in}}\, n_2 + \mathrm{J_{out}}\, n_2 = \mathrm{J_{in}}\, v + n_2(\mathrm{J_{in}} + \mathrm{J_{out}}) \\ \mathrm{J_{out}}\, n_1 + \mathrm{J_{in}}\, n_2 = \mathrm{J_{out}}\, n_1 - \mathrm{J_{out}}\, n_2 + \mathrm{J_{out}}\, n_2 + \mathrm{J_{in}}\, n_2 = \mathrm{J_{out}}\, v + n_2(\mathrm{J_{in}} + \mathrm{J_{out}}) \end{cases} \tag{S18}$$

We plug them into eq. (S17):

$$f(v) = \frac{dv}{dt} = -v + \frac{1/k}{1 + \exp\left(\frac{-k(\mathrm{J_{in}}\, v + n_2(\mathrm{J_{in}} + \mathrm{J_{out}})) + \eta}{T}\right)} - \frac{1/k}{1 + \exp\left(\frac{-k(\mathrm{J_{out}}\, v + n_2(\mathrm{J_{in}} + \mathrm{J_{out}})) + \eta}{T}\right)} \tag{S19}$$

Similarly to the symmetric case, we take $df(v)/dv$:

$$\frac{df(v)}{dv} = -1 + \frac{\mathrm{J_{in}}\left[\operatorname{sech}\left(\frac{k(\mathrm{J_{out}}\, n_2 + \mathrm{J_{in}}(v + n_2)) - \eta}{2T}\right)\right]^2 - \mathrm{J_{out}}\left[\operatorname{sech}\left(\frac{k(\mathrm{J_{in}}\, n_2 + \mathrm{J_{out}}(v + n_2)) - \eta}{2T}\right)\right]^2}{4T} \tag{S20}$$

We get back to $n_1$ and $n_2$:

$$\frac{df(v)}{dv} = -1 + \frac{\mathrm{J_{in}}\left[\operatorname{sech}\left(\frac{k(\mathrm{J_{in}}\, n_1 + \mathrm{J_{out}}\, n_2) - \eta}{2T}\right)\right]^2 - \mathrm{J_{out}}\left[\operatorname{sech}\left(\frac{k(\mathrm{J_{out}}\, n_1 + \mathrm{J_{in}}\, n_2) - \eta}{2T}\right)\right]^2}{4T} \tag{S21}$$

Therefore, to get the intermittent phase, we should solve

$$\begin{cases} n_1 = \dfrac{1/k}{1 + \exp\left(\frac{-k(\mathrm{J_{in}}\, n_1 + \mathrm{J_{out}}\, n_2) + \eta}{T}\right)} \\ n_2 = \dfrac{1/k}{1 + \exp\left(\frac{-k(\mathrm{J_{in}}\, n_2 + \mathrm{J_{out}}\, n_1) + \eta}{T}\right)} \\ \dfrac{df(v)}{dv} = 0 = -1 + \dfrac{\mathrm{J_{in}}\left[\operatorname{sech}\left(\frac{k(\mathrm{J_{in}}\, n_1 + \mathrm{J_{out}}\, n_2) - \eta}{2T}\right)\right]^2 - \mathrm{J_{out}}\left[\operatorname{sech}\left(\frac{k(\mathrm{J_{out}}\, n_1 + \mathrm{J_{in}}\, n_2) - \eta}{2T}\right)\right]^2}{4T} \end{cases} \tag{S22}$$

with respect to $\eta$, $n_1$, $n_2$ for each set $T$, $\mathrm{J_{in}}$, $\mathrm{J_{out}}$. Here, we seek for $n_1 \neq n_2$.

It turns out that the numerical solving is more stable if we solve the equations above with respect to $v$ together with $n_2$:

$$\begin{cases} \dfrac{dv}{dt} = 0 = -v + \dfrac{1/k}{1 + \exp\left(\frac{-k(\mathrm{J_{in}}\, v + n_2(\mathrm{J_{in}} + \mathrm{J_{out}})) + \eta}{T}\right)} - \dfrac{1/k}{1 + \exp\left(\frac{-k(\mathrm{J_{out}}\, v + n_2(\mathrm{J_{in}} + \mathrm{J_{out}})) + \eta}{T}\right)} \\ n_2 = \dfrac{1/k}{1 + \exp\left(\frac{-k(\mathrm{J_{in}}\, n_2 + \mathrm{J_{out}}(v + n_2)) + \eta}{T}\right)} \\ \dfrac{df(v)}{dv} = 0 = -1 + \dfrac{\mathrm{J_{in}}\left[\operatorname{sech}\left(\frac{k(\mathrm{J_{in}}(v + n_2) + \mathrm{J_{out}}\, n_2) - \eta}{2T}\right)\right]^2 - \mathrm{J_{out}}\left[\operatorname{sech}\left(\frac{k(\mathrm{J_{out}}(v + n_2) + \mathrm{J_{in}}\, n_2) - \eta}{2T}\right)\right]^2}{4T} \end{cases} \tag{S23}$$

*e. Phase diagram*

We modify the IIM's phase diagram to incorporate the asymmetric interaction within the spin groups and between the groups (fig. S1(a)). We take $\mathrm{J_{out}}$ as an example.

By solving eq. (S8), we obtain the second-order transition line (blue lines in fig. S1(b)). The darker blue line corresponds to the low-activity state $V_0 \leq 1/2$, where the fractions of active spins $n_1 = n_2 \leq 0.25$ (the solutions along the line are shown in fig. S1(c)), while the light blue line corresponds to the high-activity state $V_0 \geq 1/2$, where the fractions $n_1 = n_2 \geq 0.25$. In both cases, the velocity of the DV is zero because at this transition lines, the compromise solution becomes stable: $V = n_1 - n_2 = 0$ (fig. S1(d)).

We obtain the second-order transition line by solving eq. (S23) (purple and magenta lines in fig. S1(b)). The purple line bounds (together with the blue lines) the phase, where the low-activity compromise state co-exist with the decision state (two non-zero solutions for $V$; purple lines in fig. S1(c), (d)), while the magenta line bounds the area, where the high-activity compromise state co-exist with the decision state (two non-zero solutions for $V$).

In fig. S2, we demonstrate how the phase diagram and the solutions for the spin activity in both groups and the velocity along the transition lines change if we vary $\mathrm{J_{out}}$. It turns out that the size of the ordered phase increases when we increase $|\,\mathrm{J_{out}}\,|$, and the ordered phase completely disappears for $\mathrm{J_{out}} \geq 0$. The transition lines bound the intermittent phase at low temperatures, while the entire are around it is the disordered phase (fig. S2(d)).

Finally, in fig. S3, we show the solutions as functions of inhibition at constant temperature (along the black vertical line). The compromise solutions are represented by the blue lines (darker blue for the low-activity solution and the light blue for the high-activity solution) and the non-compromise solutions are represented by the purple and magenta lines (for the activities in the first and second spin groups). The blue and purple vertical lines indicate the transition lines at the fixed temperature.

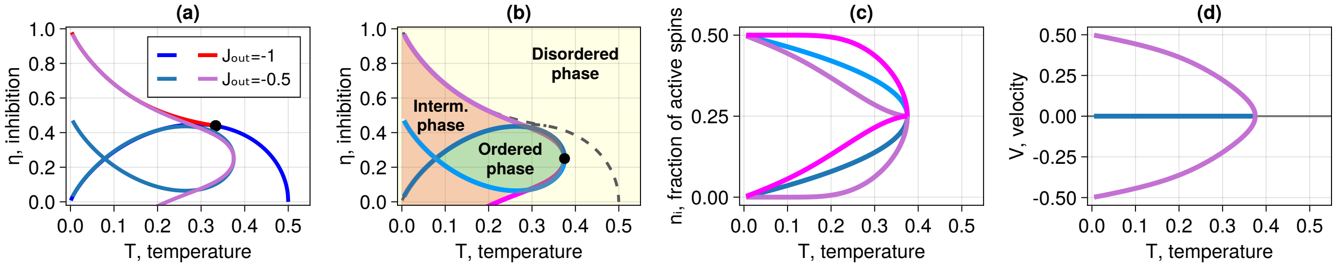


Figure S1: (a) Phase diagram of the IIM with asymmetric interactions ($J_{in} = 1$, $J_{out} = -0.5$) vs. symmetric interactions ($J_{in} = 1$, $J_{out} = -1$). The pale cornflower-blue line and the purple line indicate the second and the first-order transitions for $J_{out} = -0.5$, respectively. The bright blue line and the red line indicate the second and the first-order transitions for $J_{out} = -1$, respectively. The black circle is the tricritical point in the symmetric case. (b) Phases on the phase diagram for the IIM with asymmetric interactions: self-excitation within each group $J_{in} = 1$, cross-inhibition between different groups $J_{out} = -0.5$. The two blue lines indicate the second-order phase transitions (darker blue for the low-activity MF compromise solution $n_1 = n_2 \leq 0.25$, lighter blue for the high-activity compromise solution $n_1 = n_2 \geq 0.25$). The purple and magenta lines are the first-order transition lines. The black circle is the tricritical point in the asymmetric case (given by eq. (S15), eq. (S16)). The grey dashed lines refer to the symmetric case ($J_{out} = -1$). (c) MF solutions for the mean activities in the two groups (fractions of active spins). The blue lines indicate the MF solutions along the second-order transition lines of the corresponding color (compromise). The purple and magenta lines show the MF activities in the two groups for the non-compromise regime along the first-order transition lines of the corresponding color. (d) MF velocity ($V = n_1 - n_2$). The blue line indicates the MF velocity along the second-order transition lines (compromise). The purple line shows the MF velocity for the non-compromise regime along the first-order transition lines.

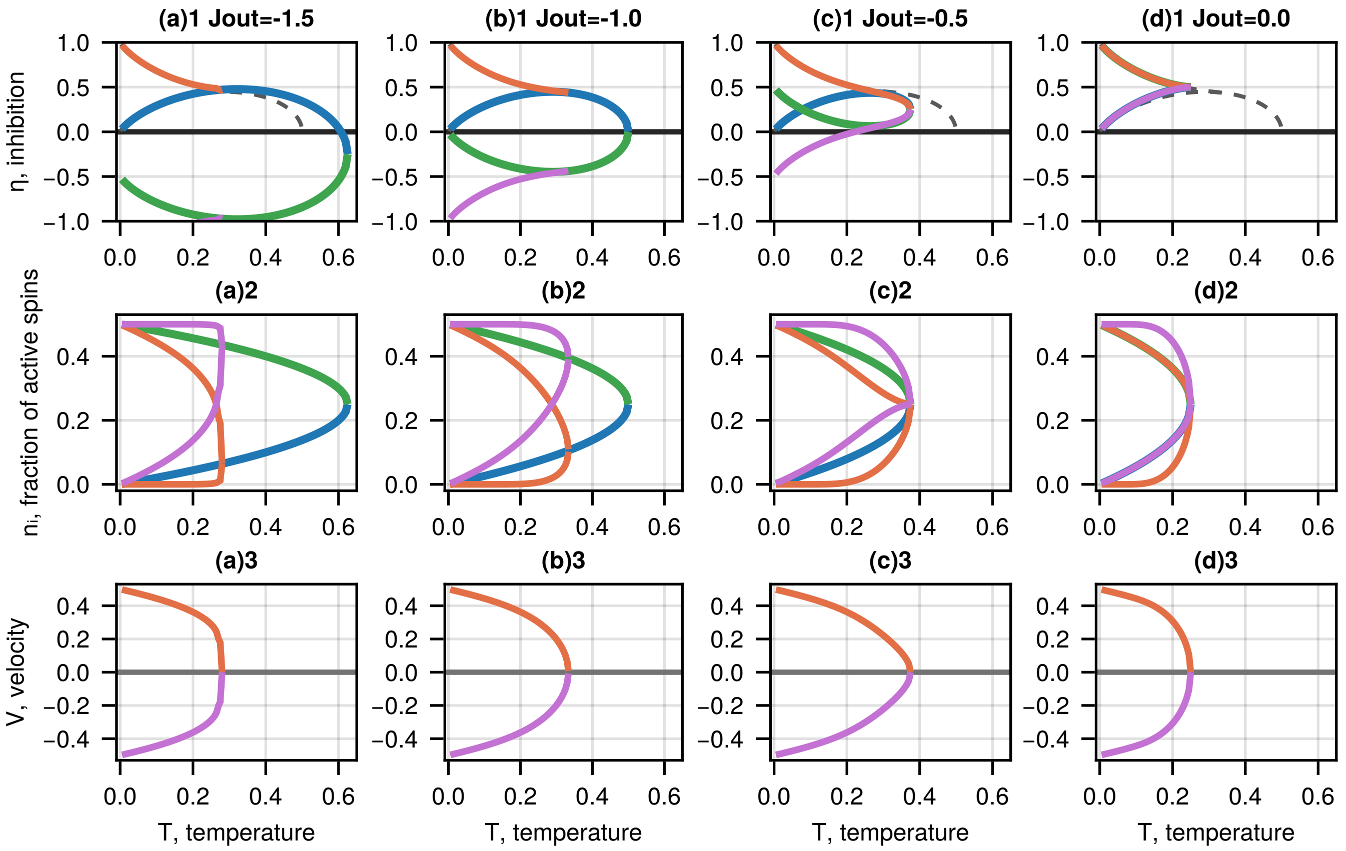


Figure S2: (a)1 Phase diagram of the IIM with asymmetric interactions ($J_{in} = 1$, $J_{out} = -1.5$). The blue and green lines indicate the second-order transition. The purple and orange lines indicate the first-order transition. The dashed lines indicate the phase diagram for $J_{out} = -1$. (a)2 MF solutions for the mean activities in the two groups (fractions of active spins). The blue and green lines indicate the MF solutions along the second-order transition lines of the corresponding color (compromise). The purple and orange lines show the MF activities in the two groups for the non-compromise regime along the first-order transition lines of the corresponding color. (a)3 MF velocity ($V = n_1 - n_2$) for the non-compromise regime along the first-order transition lines. (b) $J_{out} = -1$. (c) $J_{out} = -0.5$. (d) $J_{out} = 0$.

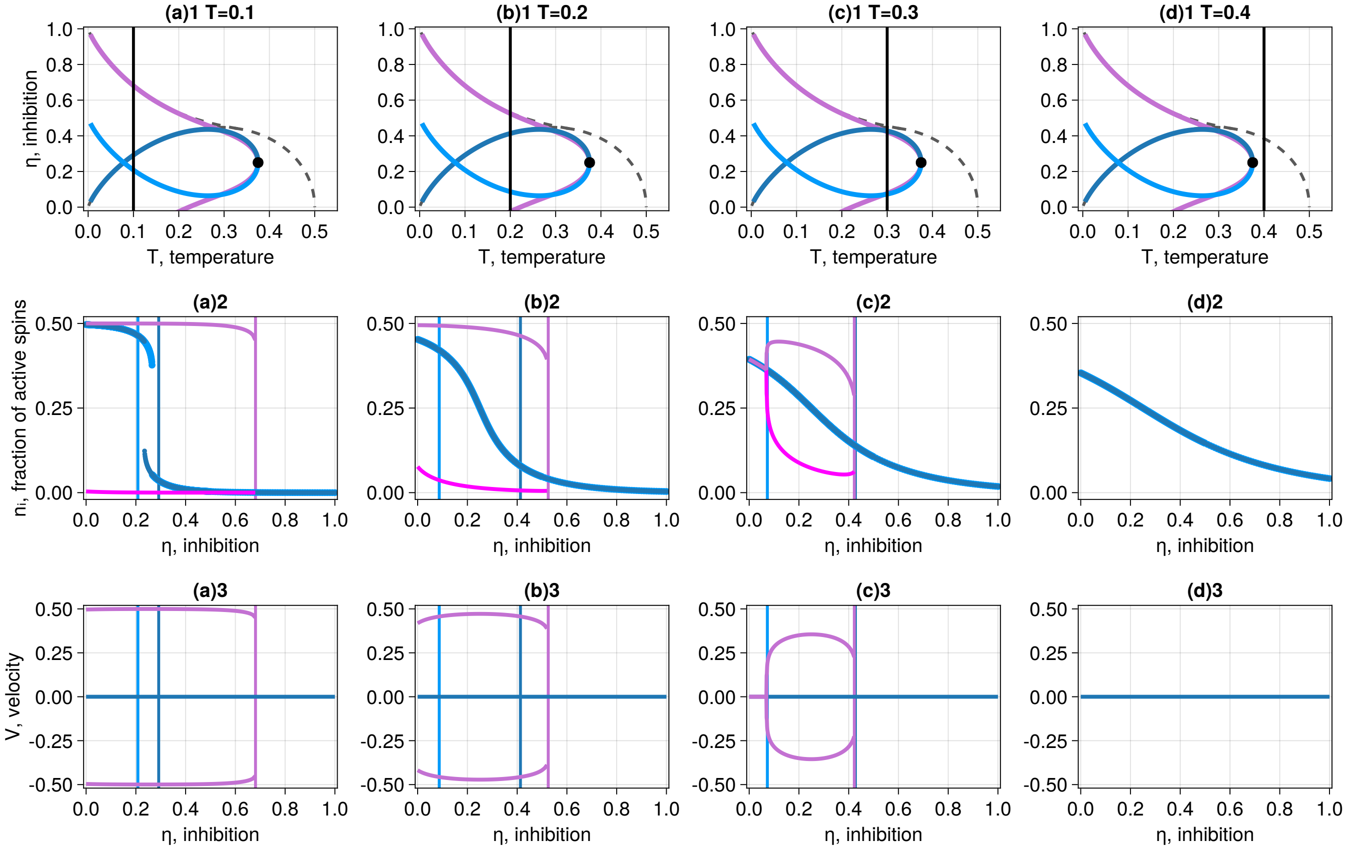


Figure S3: (a)1 Phase diagram of the IIM with asymmetric interactions with self-excitation within each group $\mathrm{J_{in}} = 1$ and cross-inhibition between different groups $\mathrm{J_{out}} = -0.5$ (blue and purple lines) and $\mathrm{J_{out}} = -1$ (grey dashed lines). The pale cornflower-blue line indicates the area above which the high-activity compromise solution becomes unstable (second-order transition). The light blue line shows the area below which the low-activity compromise solution gets unstable (another second-order transition). The purple lines are the first-order transition lines. The black circle is the tricritical point in the asymmetric case (given by eq. (S15), eq. (S16)). The black vertical line indicates $T = 0.1$. (a)2 MF solutions for $n_1$, $n_2$ as functions of the global inhibition $\eta$ along a fixed temperature $T = 0.1$. The blue curves show the compromise solutions (high and low activity if they exist). The purple and magenta lines indicate $n_1$ and $n_2$, respectively, for the non-compromise solution. The vertical lines denote the corresponding transition lines at a fixed temperature $T = 0.1$. (a)3 MF solutions for velocity $V$ as a function of the global inhibition $\eta$ along a fixed temperature $T = 0.1$. The blue line indicates the compromise solution, and the purple lines indicate the non-compromise solutions. (b) $T = 0.2$, (c) $T = 0.3$, (d) $T = 0.4$.

### S2. IIM PARAMETERS AND NUMERICAL SIMULATIONS

#### A. IIM parameters

All numerical simulations and analyses of the experimental data were implemented in Julia [2]. The values of the IIM's parameters used in this work are given in table S1.

Table S1: Parameters, used in the numerical simulations of the IIM

| Parameter | Symbol | Range |
|---|---|---|
| Temperature | $T$ | $0\dots1$ |
| Global inhibition | $\eta$ | $0\dots1$ |
| Bias | $\epsilon_1$ | $0\dots1$ |
| Self-excitation | $J_{\text{in}}$ | 1 |
| Cross-inhibition | $J_{\text{out}}$ | -1 |
| Decision threshold | $L$ | 40 |
| Total number of spins | $N$ | 50 |
| Number of spins in group I | $N^I$ | 25 |
| Number of spins in group II | $N^{II}$ | 25 |
| Initial number of active spins, IC: ZERO | $N_1^{I,II}$ | 0 |
| Initial number of active spins, IC: RAND | $N_1^{I,II}$ | $0\dots25$ |

#### B. Number of spins

The spin system consists of two equal subgroups, which refer to the given alternatives in a decision task. We investigate how the decision properties depend on the number of spins in the subgroups.

It turns out that in the case of small groups, the system cannot be approximated by the mean-field (MF) theory and behaves differently compared to a larger system. For large systems, the gradient of both error and reaction time (RT) becomes sharper near the phase transition, and it also requires more computational power for simulations (fig. S4).

For our simulations, we chose the size of the subgroups of 25 spins. As a result, the model's behavior is similar to the MF prediction, while numerical simulations remain relatively fast (black line in fig. S4).

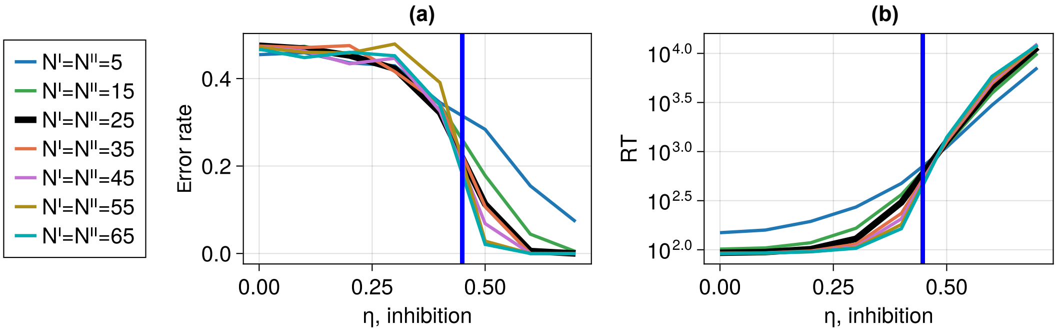


Figure S4: (a) Error rate, (b) Reaction time (RT) as functions of the global inhibition $\eta$ at fixed temperature $T = 0.3$, bias $\epsilon_1 = 0.01$, and variable numbers of spins in each subgroup $N^{I,II}$ (denoted by different colors). The thick black line indicates the value, used in all simulations in the main part ($N^{I,II} = 25$). The blue vertical lines indicate the second-order transition at $\eta = 0.447$.

#### C. Decision threshold

The properties of the IIM depend on the threshold value $L$, which determines when the stochastic processes end. At fixed bias, the probability of errors decreases with $L$, while the RT increases with $L$ (fig. S5). This paragraph explores the dependence of the decision properties in the IIM on temperature and inhibition in different regions of the phase space for lower and higher thresholds ($L = 10,\ 100$) than we present in the main text ($L = 40$, fig. 2).

It turns out that the dependencies of the decision properties (error rate, RT, the RT ratio in the correct and wrong decisions $RT_c/RT_w$) on the system's parameters $T$, $\eta$ remain similar across different threshold values (fig. S6). It allows us to fix a certain threshold and adjust the corresponding bias to achieve the desired error rate for each point of the phase space while fitting the IIM to the experimental data.

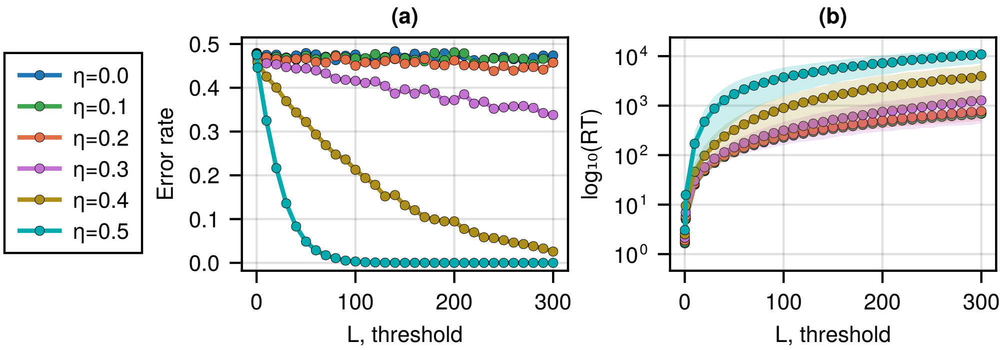


Figure S5: Error rate, reaction time (RT), and the RT ratio in the correct and wrong decisions as functions of the system's parameters ($\eta$, $T$) at a fixed bias $\epsilon_1 = 0.01$ for the zero IC for different thresholds, presented as the heatmaps (the color bars indicate the values). (a) Error rate, (b) and RT (mean $\pm$ STD) as functions of the threshold value $L$ at fixed temperature $T = 0.3$, cross-inhibition $\mathrm{J_{out}} = 1$, and bias $\epsilon_1 = 0.01$, while the global inhibition varies (denoted by color).

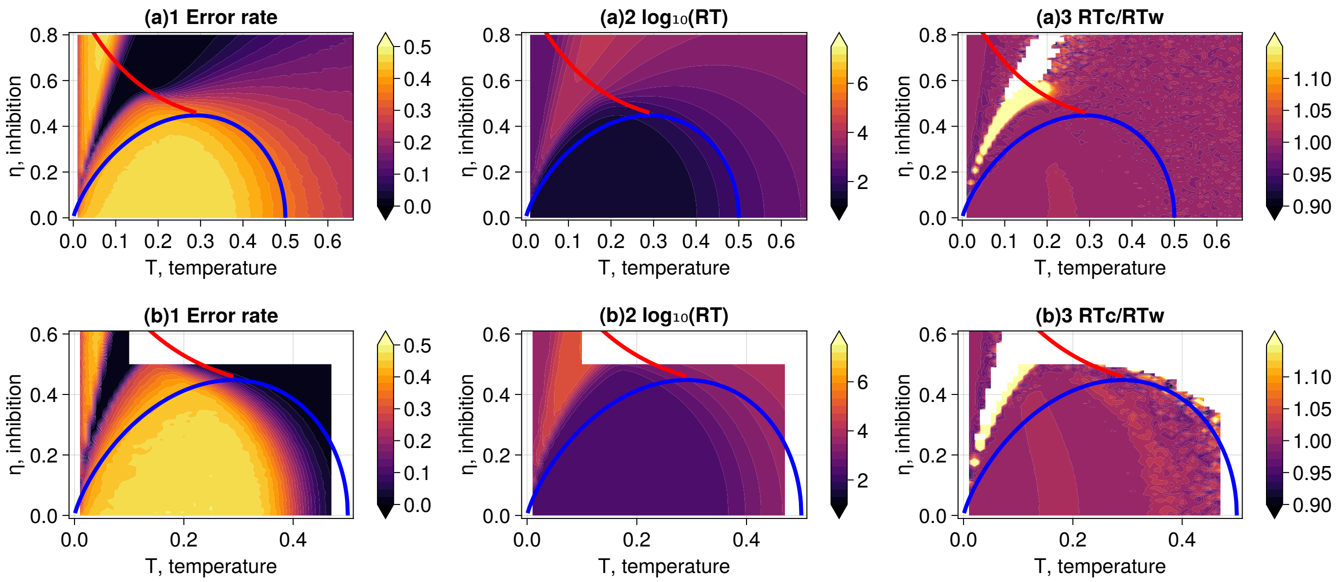


Figure S6: Error rate, reaction time (RT), and the RT ratio in the correct and wrong decisions as functions of the system's parameters ($\eta$, $T$) at a fixed bias $\epsilon_1 = 0.01$ for the zero IC for different thresholds, presented as the heatmaps (the color bars indicate the values). The red and blue lines on the heatmaps denote the first and second-order transitions, respectively. (a) $L = 10$. (b) $L = 100$.

#### D. Temperature, global inhibition, bias, cross-inhibition

In this paragraph, we focus on the dependence of the decision's properties on the system's parameters (temperature $T$, global inhibition $\eta$, bias $\epsilon_1$, and cross-inhibition $\mathrm{J_{out}}$) at a fixed threshold $L = 40$ and the zero initial conditions, where all the spins are initially turned "off" ($\sigma_i = 0$). We find that both error rate and RT decrease as the bias increases (fig. S7(a)), which means that if one option is distinctly more favorable than the other option, the decisions are faster and more accurate.

If the global inhibition or temperature (noise) increase (fig. S7(b), (c)), the dynamics of the decision processes changes from the ordered to the disordered regime, and therefore, the error rate decreases, while the RT increases. In contrast, if we increase cross-inhibition, the error rate increases (fig. S8(a)), while the RT decreases (fig. S8(b)).

#### E. Random initial conditions

In this section, we demonstrate the results of the IIM with the random IC, where the initial state of each spin is random ($\sigma_i = 0$ or 1). The dependencies of the IIM properties on the system's parameters $T$, $\eta$ (fig. S9) resemble the results for the zero IC (fig. 2 in the main text).

For the random IC, the error rate in the intermittent phase at lower inhibition tends to be larger than for the zero IC (between the blue and red lines in fig. S9(a)1). It happens because for the zero IC, where all the spins are initially turned "off" ($\sigma_i = 0$), the initial velocity is always zero, and it is stuck at this value for an extended period of time

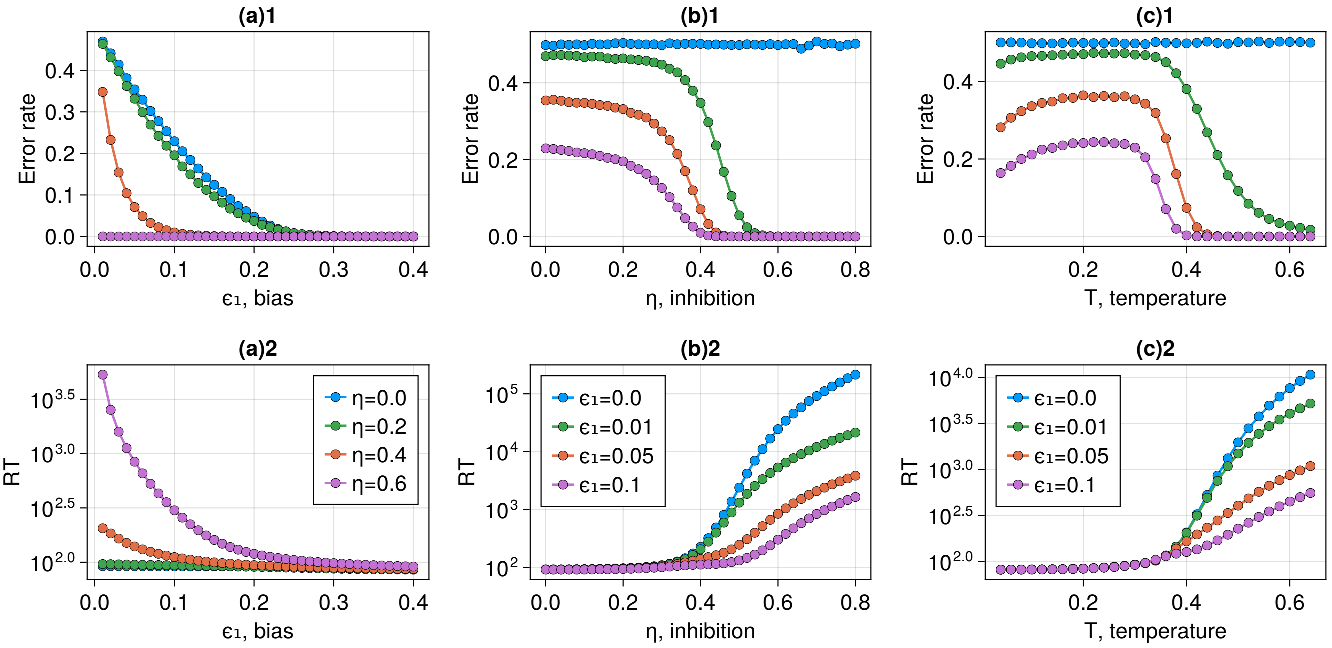


Figure S7: Dependence of the decision properties in the IIM on the model's parameters (global inhibition $\eta$, temperature $T$, bias $\epsilon_1$) for the zero IC. (a) Error rate and reaction time (RT) as functions of bias at fixed temperature $T = 0.3$. (b) Error rate and reaction time (RT) as functions of the global inhibition at fixed temperature $T = 0.3$. (c) Error rate and reaction time (RT) as functions of temperature at fixed inhibition $\eta = 0$.

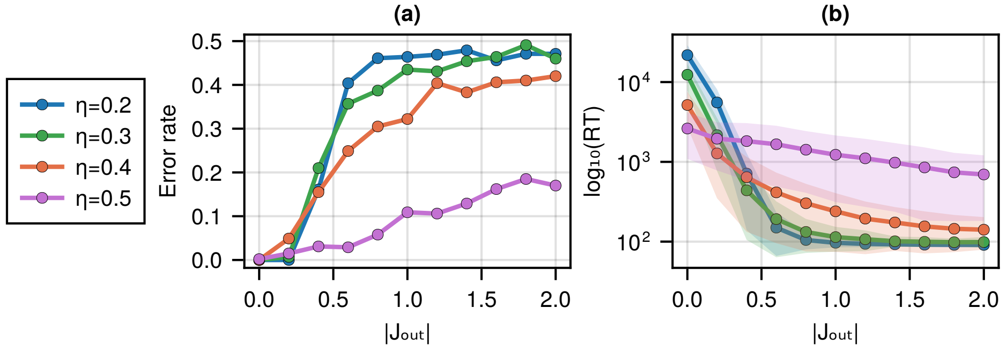


Figure S8: (a) Error rate, (b) and RT (mean ± STD) as functions of the cross-inhibition $J_{out}$ (its absolute value) at fixed temperature $T = 0.3$ and bias $\epsilon_1 = 0.01$, while the global inhibition varies (denoted by color).

until the velocity spontaneously changes its value to one of the non-zero MF solutions and continues to move to a threshold ballistically. At the same time, at the random IC, the probability of starting with a negative velocity and the rate of reaching the negative MF velocity is higher. Similarly, the near RT in the intermittent phase is higher for the random IC (fig. S9(b)1).

The RT ratio in the correct and wrong decisions in the intermittent phase (yellow area in fig. S9(c)1) is above 1, because in simulations, if the initial velocity is negative, it quickly reaches the negative MF solution and continues to move to the threshold. If the initial velocity is zero, it will likely get stuck and switch to the positive value with a higher probability than to the negative solution. Therefore, in the simulations, we sample more long trajectories that reach the positive threshold than the negative one, and the mean RT in the correct decisions appears to be higher.

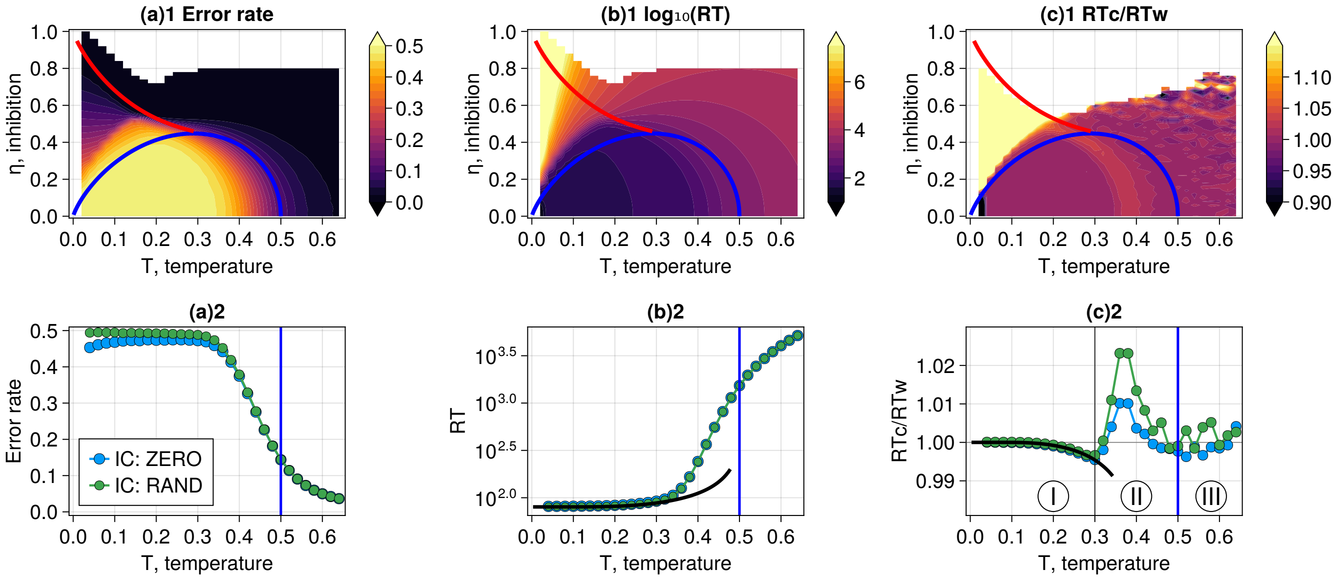


Figure S9: (a)1 Error rate, (b)1 RT, (c)1 the RT ratio in the correct and wrong decisions as functions of the system's parameters ($\eta$, $T$) at fixed bias $\epsilon_1 = 0.01$ for the random IC, presented as the heatmaps (the color bars indicate the values). The red and blue lines on the heatmaps denote the first and second-order transitions, respectively. (a)2 Error rate, (b)2 RT, (c)2 the RT ratio in the correct and wrong decisions as functions of temperature at fixed bias $\epsilon_1 = 0.01$ for the random IC (green line) and the zero IC (blue line).

### S3. DIFFERENT REGIMES OF THE IIM

In this section, we explore the properties of the IIM dynamics in different areas of the phase diagram that correspond to different regimes of motion: ballistic (I), run-and-tumble (II), and diffusion (III), marked in fig. S10(c)2 (similar to fig. 2 in the main text). We derive the analytical expressions to estimate the error rate and the mean RT in the IIM for each regime separately.

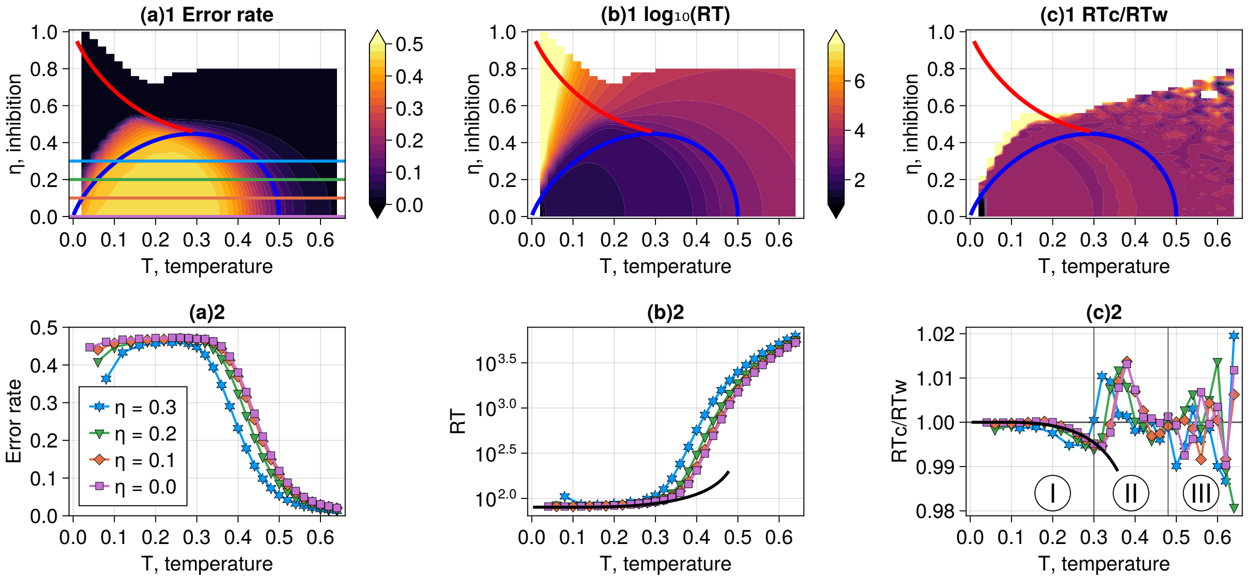


Figure S10: (a)1 Error rate, (b)1 reaction time (RT), (c)1 the RT ratio in the correct and wrong decisions as functions of the system's parameters ($\eta$, $T$) at fixed bias $\epsilon_1 = 0.01$ for the zero initial conditions (IC: ZERO), presented as the heatmaps (the color bars indicate the values). The red and blue lines on the heatmaps denote the first and second-order transitions, respectively. (a)2 Error rate, (b)2 reaction time (RT), (c)2 the RT ratio in the correct and wrong decisions as functions of temperature for variable levels of inhibition, denoted by different colors in (a)1. The RT ratio in the correct and wrong decisions depends on the region of the phase space. In zone I, the movement is ballistic. In zone II, the dynamics can be described as the run-and-tumble process. In zone III, the process is drift-diffusion. The black theoretical line indicates the behavior of the RT and the RT ratio in the correct and wrong decisions at low temperatures using the ballistic approximation (eq. (9)).

#### A. RT distributions

We explore the RT distributions in each zone in numerical simulations and estimate the average duration of decision trajectories that reach the correct (positive) and wrong (negative) thresholds (fig. S11). It turns out that the RT distributions are heavily right-skewed [3], and they become wider as we increase temperature $T$. Depending on the location of the parameters in the phase space, the ratio $\mathrm{RT_c/RT_w}$ can be both below 1 ("slow errors") or above 1 ("fast errors").

To quantitatively compare the shapes of the RT distributions, we determine the skewness and kurtosis of the RT distributions for different areas of the phase space (fig. S12). We find that the skewness of the RT distributions reaches its maximal values in the ordered phase (yellow area), where the ballistic trajectories co-exist together with non-linear trajectories (run-and-tumble), and decreases in the disordered phase at higher temperature and inhibition.

For each phase, we choose one point (marked as stars in fig. 2E in the main text and fig. S12) and estimate the asymmetry of the RT distributions. For the direct comparison, we show the exact parameters that are used to describe skewed distributions (table S2).

Table S2: Comparison of the parameters of the normalized RT distributions, given in different phases.

| **IIM's Parameters** | **Mean** | **Median** | **STD** | **Skewness** | **Kurtosis** | $L/V^+_{\mathbf{MF}}$ | **# simulations** |
|---|---|---|---|---|---|---|---|
| Intermittent: $T = 0.18, \eta = 0.45, \epsilon_1 = 0.01$ | 1 | 0.93 | 0.24 | 3.24 | 16.69 | 0.76 | $10^5$ |
| Ordered: $T = 0.3, \eta = 0.36, \epsilon_1 = 0.01$ | 1 | 0.75 | 0.56 | 2.71 | 9.55 | 0.65 | $2 \times 10^4$ |
| Disordered: $T = 0.44, \eta = 0.56, \epsilon_1 = 0.01$ | 1 | 0.83 | 0.67 | 2.08 | 6.67 | 0.96 | $10^3$ |

The parameters of IIM ($\eta$, $T$) are given in fig. 2C(ii) in the main text and marked in fig. S12. $RT^+_{bal} = L/V^+_{\mathrm{MF}}$ (eq. (8)) indicates the theoretical prediction for the mean RT using the MF velocity.

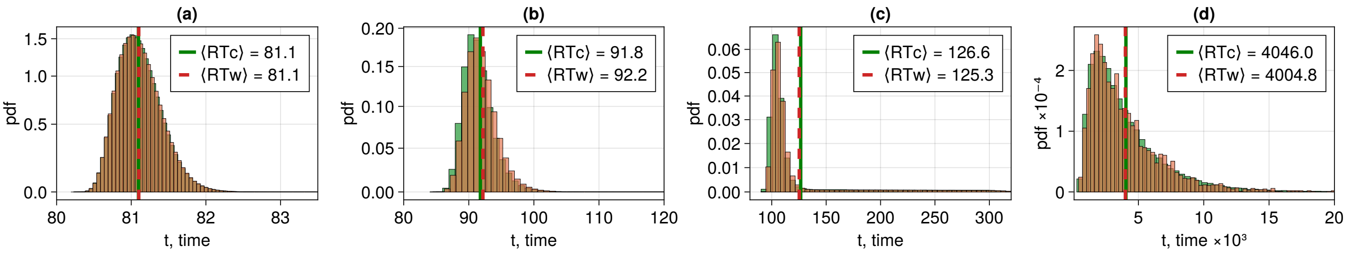


Figure S11: RT distributions for the correct (green) and wrong (orange) decisions in the absence of the global inhibition ($\eta = 0$) at fixed bias $\epsilon_1 = 0.01$. The green and red vertical lines indicate the mean RT in the correct and wrong decisions, respectively. (a) Zone I: $T = 0.06$. The error rate is 0.4608, $\mathrm{RT_c}/\mathrm{RT_w}$ is 0.999958. (b) Zone II: $T = 0.3$. The error rate is 0.4726, $\mathrm{RT_c}/\mathrm{RT_w}$ is 0.9958. (c) Zone II: $T = 0.36$. The error rate is 0.4414, $\mathrm{RT_c}/\mathrm{RT_w}$ is 1.0108. (d) Zone III: $T = 0.6$. The error rate is 0.0516, $\mathrm{RT_c}/\mathrm{RT_w}$ is 1.0103.

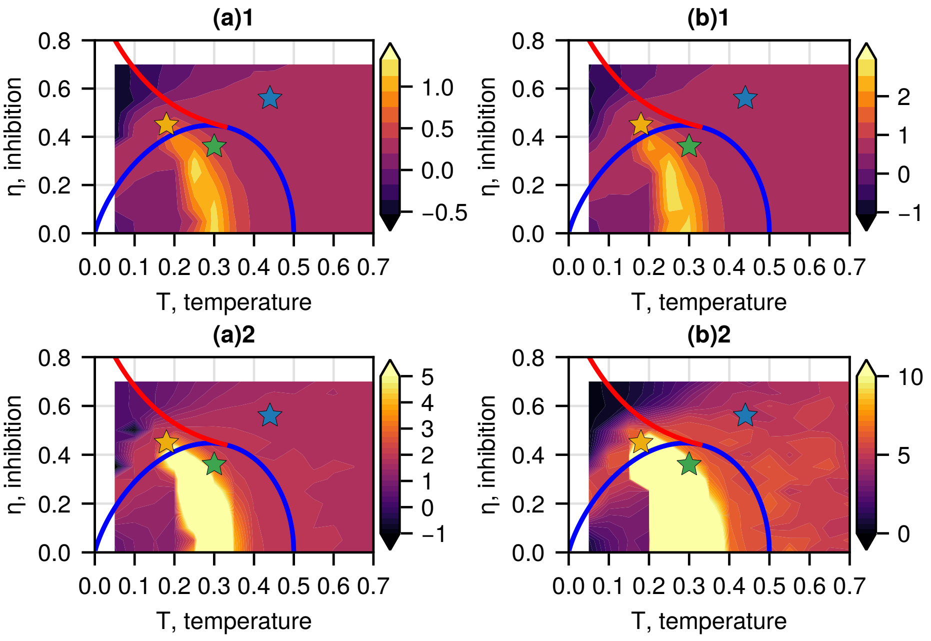


Figure S12: (a) Skewness of the RT distributions: (a)1 log-scale, (a)2 linear scale with the maximal value of 5 on the colorbar. (b) Kurtosis of the RT distributions: (b)1 log-scale, (b)2 linear scale with the maximal value of 10 on the colorbar. The quantities and measured at a constant bias $\epsilon_1 = 0.01$ in $10^5$ simulations per point in the ordered phase and $5 \times 10^3$ simulations per point in the disordered and intermittent phases. The stars indicate the location of the points, shown in table S2.

#### B. Zone I: ballistic motion (low temperature, low inhibition)

At low temperature and inhibition (zone I in fig. S10(c)2), the IIM dynamics can be described by ballistic motion. At the zero initial conditions (IC: ZERO), all the spins are initially turned "off" ($\sigma_i = 0$), and the initial velocity is zero. Therefore, in the limit of very low temperatures and at fixed bias, the initial spin flip determines the direction of the ballistic trajectory. Then, the probability of reaching the negative threshold (error rate) is analytically predicted as the ratio of the Glauber rates at the beginning of the process (eq. (6))

$$Error\Big|_{\substack{\text{ballistic regime}\\ \text{IC: ZERO}}} = \frac{r^{II}_{0\to1}}{r^{I}_{0\to1} + r^{II}_{0\to1}}\Big|_{V=0} = \frac{1}{1 + \frac{1+e^{\frac{\eta}{T}}}{1+e^{\frac{\eta-\epsilon_1}{T}}}} = \frac{e^{-\frac{\eta}{T}} + e^{-\frac{\epsilon_1}{T}}}{2e^{-\frac{\eta}{T}} + e^{-\frac{\epsilon_1}{T}} + 1} \tag{S24}$$

Therefore, we can predict the error rate in the limit of low temperatures (solid lines in fig. S13(a)). Similarly, we can find the corresponding bias for a fixed error (solid lines in fig. S13(b))

$$\epsilon_1 = \eta - T \ln\left[\frac{1 - Error\left(2 + e^{\frac{\eta}{T}}\right)}{-1 + Error}\right] \tag{S25}$$

where the solution exists if $\eta \geq 0$ and $1/3 < Error < 1/2$. Otherwise, $0 < Error \leq 1/3$ and $\eta > T \ln\left(\frac{1}{Error} - 2\right)$.

As temperature increases, more flips might happen at the initial stage of the trajectory, and the first spin flip no longer determines the final decision. The trajectories in this regime are almost linear with a slight delay at the beginning. The RT distribution in the simulations with the fixed threshold is narrow and skewed towards longer times for both correct and wrong choices (fig. S11(a)).

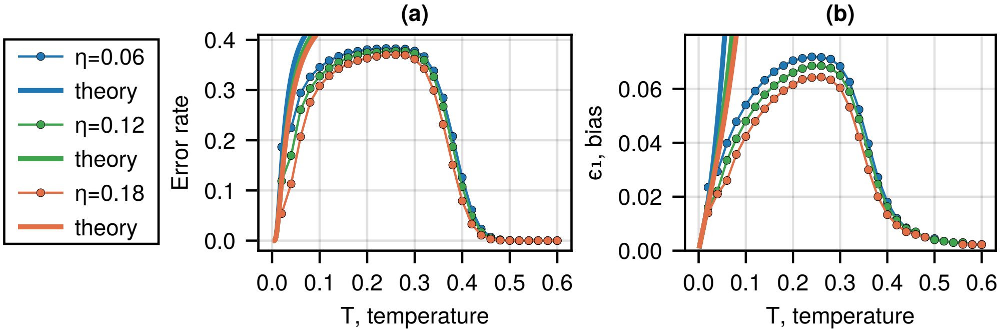


Figure S13: (a) The error rate as a function of temperature and global inhibition (denoted by color) at fixed bias $\epsilon_1 = 0.04$ (the lines with circles). The solid lines indicate the theoretical predictions at different inhibition levels (denoted by color) for the ballistic motion at low temperatures (eq. (S24)). (b) Bias as a function of temperature and global inhibition (denoted by color) at a fixed error rate of $0.3 \pm 0.01$. The solid lines indicate the theoretical predictions at different inhibition levels (denoted by color) for the ballistic motion at low temperatures (eq. (S25), eq. (8)).

#### C. Zone III: drift-diffusion (high temperature, high inhibition)

This section describes the high-temperature and high-inhibition disordered regime of the IIM (zone III in fig. S10(c)2) using the drift-diffusion model (DDM) and investigates the main properties of the diffusion process with a constant drift in an interval. The advantage of the DDM is that it is able to fully estimate the outcomes analytically, while the MF theory gives only an approximation for the IIM.

We consider a diffusing particle in the interval $[-L; L]$ with absorbing boundaries, which means that eventually, the particle touches one of the thresholds and leaves the system immediately. Then, the stochastic Langevin equation of a freely accelerated particle starting at point $x_0 \in [-L; L]$ at $t_0 = 0$ looks as follows:

$$\frac{dx}{dt} = v + \sqrt{2D}\eta(t) \tag{S26}$$

where $\eta$ represents the Gaussian white noise: $\langle \eta(t) \rangle = 0$ and $\langle \eta(t)\eta(t') \rangle = \delta(t - t')$. The white noise is almost everywhere discontinuous and has infinite variation [4], thus, we use the finite difference simulations to approximate the continuous solution of eq. (S26). We can estimate the DDM parameters from the IIM as follows:

$$v = \frac{\langle x(t) \rangle}{t}; \quad D = \frac{\langle x^2(t) \rangle - \langle x(t) \rangle^2}{2t} \tag{S27}$$

The diffusion process and its first-passage properties are determined by the forward Fokker–Planck equation (also called convection-diffusion equation), which describes the evolution of the particle's concentration $c(x, t|x_0)$ in space and time. The concentration represents the probability of finding the particle at position $x$ at time $t$ if it started at $x_0$ at $t_0 = 0$ (see the derivation in chapter 5.2 in [5], and chapter 2 in [6]):

$$\frac{\partial c(x, t|x_0)}{\partial t} + v\frac{\partial c(x, t|x_0)}{\partial x} = D\frac{\partial^2 c(x, t|x_0)}{\partial x^2} \tag{S28}$$

The particle starts at $x_0$, then, the initial condition is $c(x, t = 0|x_0) = \delta(x - x_0)$. The boundary conditions are $c(x = -L, t|x_0) = c(x = L, t|x_0) = 0$, showing that the particle leaves the system immediately, once it touches the boundaries. The constant velocity $v$ in this model introduces a bias in the system.

In the following paragraphs, we find analytically the proportion of the correct and wrong choices and the mean first-passage time (MFPT) to reach the ends of the interval to estimate the error rate and the RT in the decision processes.

The splitting probability $\epsilon^{\pm}$ at $x_0 \in [-L; L]$ is the probability for a particle to hit a threshold and leave the system through either $x = +L$ or $x = -L$ without touching the other boundary, and starting at $x_0$ at $t = 0$. It obeys the backward Fokker–Planck equations (see section 5.2.8 in [5] and section 1.6 in [6]):

$$\begin{cases} D\nabla^2_{x_0}\epsilon^{\pm}(x_0) + v \cdot \nabla_{x_0}\epsilon^{\pm}(x_0) = 0 \\ \epsilon^{+}(-L) = 0, \quad \epsilon^{+}(L) = 1 \\ \epsilon^{-}(-L) = 1, \quad \epsilon^{-}(L) = 0 \end{cases} \tag{S29}$$

where the boundary conditions indicate that the particle is immediately absorbed when it hits a threshold. For a positive bias $v > 0$, the positive threshold $x = +L$ indicates the correct option, while the negative threshold $x = -L$ implies the wrong alternative. Therefore, we use $\epsilon^-$ to assess the error rate in this process.

The mean first-passage time $T$ (MFPT) is the unconditioned mean exit time for a particle to leave the interval through any end starting at point $x_0$ (see section 5.2.7 in [5] and sections 1.6, 2.3 in [6]). By definition,

$$T(x_0) = \int_0^{\infty} \int_{-L}^{L} c(x, t|x_0) dx dt \tag{S30}$$

The MFPT in the drift-diffusion process obeys the backward equation:

$$\begin{cases} D\nabla^2_{x_0}T(x_0) + v\cdot\nabla_{x_0}T(x_0) = -1 \\ T(-L)=0, \quad T(L)=0 \end{cases} \tag{S31}$$

where the boundary conditions indicate that the particle is immediately absorbed when it starts at a threshold.

The conditioned mean exit time $T^{\pm}$ is the mean exit time for a particle to leave the interval through a specific site $x=\pm L$ without touching the other boundary, which obeys

$$\begin{cases} D\nabla^2_{x_0}\left[\epsilon^{\pm}(x_0)T^{\pm}(x_0)\right] + v\cdot\nabla_{x_0}\left[\epsilon^{\pm}(x_0)T^{\pm}(x_0)\right] = -\epsilon^{\pm}(x_0) \\ \epsilon^{\pm}(-L)T^{\pm}(-L)=0, \quad \epsilon^{\pm}(L)T^{\pm}(L)=0 \end{cases} \tag{S32}$$

where we use the fact that $T^{+}(x=L)=0$ and $T^{-}(x=-L)=0$ for the initial conditions to find the analytical solutions. We use $T$ to assess the mean RT in the IIM, while $T^{\pm}$ correspond to the RTs in the correct and wrong decisions.

Our goal is to predict the error rate, the mean RT, and the RT ratio in the correct and wrong decisions $\mathrm{RT_c}/\mathrm{RT_w}$ using the symmetric DDM (with equal thresholds $\pm L$ for both options and $x_0=0$). We derive the analytical solutions (fig. S14(a)):

$$\begin{cases} \epsilon^{-}(x_0=0) = \frac{1}{e^{\frac{Lv}{D}}+1} \\ T(x_0=0) = T^{+}(x_0=0) = T^{-}(x_0=0) = \frac{L}{v}\tanh\left(\frac{Lv}{2D}\right) \end{cases} \tag{S33}$$

We introduce the dimensionless Péclet number $\mathrm{Pe} = vL/D$ characterizing the ratio between the diffusion and convection time scales. In the case of pure diffusion, the Pectlet number is much smaller than 1 ($\mathrm{Pe} \ll 1$). We write the decision properties in terms of the Péclet number:

$$\begin{cases} \epsilon^{-}(x_0=0) = \frac{1}{e^{\mathrm{Pe}}+1} \\ T(x_0=0) = T^{+}(x_0=0) = T^{-}(x_0=0) = \frac{L}{v}\tanh\left(\frac{\mathrm{Pe}}{2}\right) \end{cases} \tag{S34}$$

As a result, for the initial position in the middle of the interval, the conditioned times are equal [7] ($T^{+}=T^{-}$, fig. S14(a)3), meaning that the RT ratio $T^{+}/T^{-}$ is independent of the system's parameter Pe, and the DDM with the equal thresholds and the constant drift does not explain the cases where the RT ratio in the correct and wrong decisions differs from 1 (so-called fast or slow errors [8, 9]). Also, the single parameter Pe fully characterizes the proportion of the wrong choices (error rate).

### D. Zone II: run-and-tumble

In the region of the ordered phase space, close to the second-order transition line (zone II in fig. S10(c)2), the system's trajectories consist of the intervals of movement with constant velocity, interrupted by the changes in the direction of motion (fig. S15). We use the simpler run-and-tumble (RnT) process to describe the decision dynamics in the IIM in zone II and derive its first-passage properties to estimate the properties of the IIM analytically.

In the general RnT process without stops [10], a particle moves along a one-dimensional line according to the following stochastic equation :

$$\frac{dx}{dt} = v_{\sigma}\sigma(t) \tag{S35}$$

where $\sigma(t)$ is the random variable which switches between $\pm 1$ at the Poisson rates $\alpha_{R,L}$ [11], whereas $v_{\sigma}$ takes values $v_L$ or $v_R$ (left-oriented (L) if $\sigma(t)=-1$, and right-oriented (R) if $\sigma(t)=1$). Let $\alpha_R$ be the rate of change of velocity from $-v_L$ to $v_R$ (from left to right), and $\alpha_L$ is the rate of change from $v_R$ to $-v_L$ (from right to left). The particle starts at the initial position $x_0=0$ with either $v_R$ or $-v_L$ and moves inside the interval $[-L;L]$ with the absorbing boundaries, meaning that the particle disappears as soon as it hits the boundary.

The differences in the tumble rates $\alpha_{R,L}$ and velocities $v_{R,L}$ introduce a bias in the system. If $\alpha_R > \alpha_L$ and/or $v_R > v_L$, the positive threshold $+L$ denotes the correct option.

#### 1. *Parameter extraction for the run-and-tumble process*

In the IIM, at low temperatures and inhibition, the tumbles are fast relative to the running durations, and we can neglect the time it takes to make a flip compared to the characteristic time of movement with a constant velocity. However, at higher temperatures and inhibition, the process of acceleration and deceleration is considerable, so we add a fixed pause at each flip to compensate for that (fig. S15). This section aims to explain how we extract parameters for the RnT process with and without stops from the IIM. We consider these two cases separately.

We consider a large threshold $L$ such that the trajectory contains both kinds of tumbles (from left to right and from right to left): $L \gg v_{R,L}/\left(\alpha_R+\alpha_L\right)$ where $v_R$, $v_L$, $\alpha_R$, and $\alpha_L$ are the velocities and the tumble rates in the

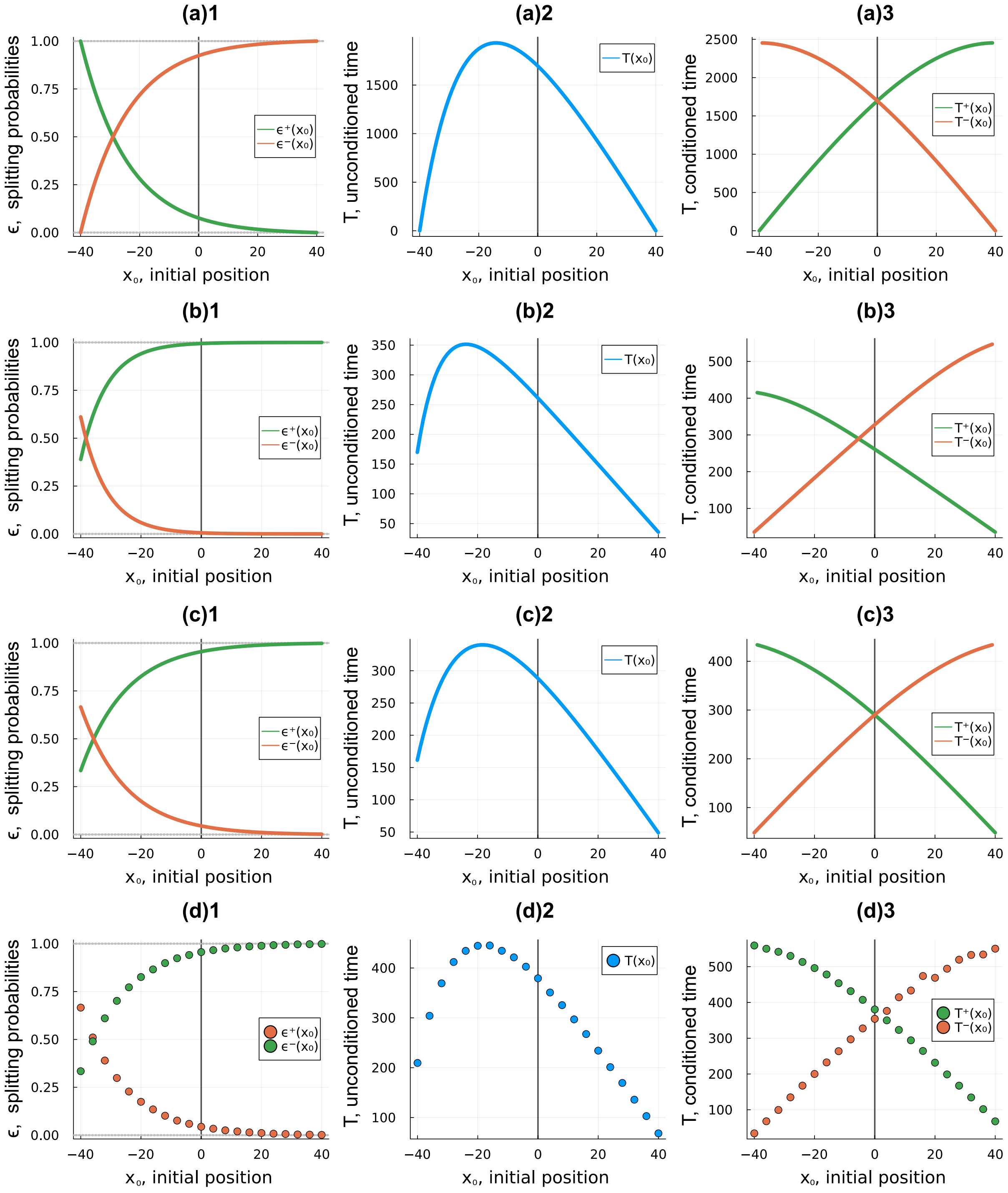


Figure S14: Splitting probabilities, unconditioned and conditioned mean first-passage times (MFPT) as functions of the particle's initial position for simple stochastic models. The solid lines are the analytical solutions. The circles are the results of the numerical simulations. The grey vertical lines indicate the particle's initial position in the IIM ($x_0 = 0$). The thresholds are symmetric ($L = \pm 40$). The models' parameters are estimated in the ordered and disordered phases of the IIM. (a) Drift-diffusion model with $v = 0.02$, $D = 0.32$ (Pe $= 2.5$). (b) Asymmetric RnT process with $\alpha_R = 0.03$, $\alpha_L = 0.01$, $v_R = 0.3$, $v_L = 0.2$. (c) Symmetric RnT process with $\alpha_R = 0.03$, $\alpha_L = 0.01$, $v_R = v_L = 0.3$. (d) Symmetric RnT process with stops with $\alpha_R = 0.03$, $\alpha_L = 0.01$, $v_R = v_L = 0.3$, $t_{\text{stop}} = 20$.

RnT process (fig. S16(a), (c)). In the case of very low tumble rates, when the trajectories contain only one tumble, we measure the mean time at which the first tumble occurs.

We average the velocity along the trajectory in a sliding window to reduce the noise and determine the zeros (see the purple curve and green dots in fig. S16(b)). Then, we divide the smoothed curve into regions corresponding to the motion with a constant velocity $v_R$ (positive direction) or $v_L$ (negative direction) in the RnT process, while the zeros indicate the tumbles. The following analysis depends on whether we estimate the RnT process with or without stops.

In the case of the regular RnT process without stops, at each fragment $i$ of the smooth curve between two zeros, we estimate the median velocity $v_{R/L,i}$ and the time $\tau_{R/L,i}$ between two tumbles, where $\tau_R$ is the time in the right-oriented

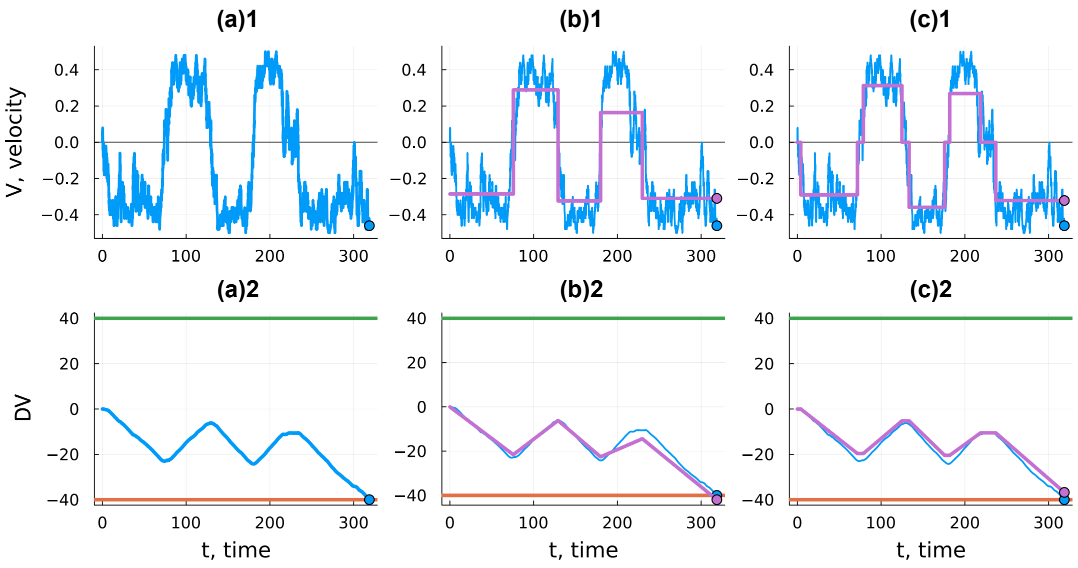


Figure S15: (a) Typical velocity dynamics and trajectory in the IIM at $\eta = 0.2$, $T = 0.36$, $\epsilon_1 = 0.026$ (zone II: the ordered phase near the critical line). The green and orange horizontal lines indicate the positive and negative thresholds. The blue circle indicates the finite state of the system as it reaches the threshold. (b) Approximation of the IIM using the RnT process without stops. (c) Approximation of the IIM using the RnT process with stops. By this mapping, we can extract the corresponding RnT parameters. The purple circle indicates the finite state of the RnT process as the IIM model reaches a threshold. The RnT finite coordinate can be both inside and outside the interval.

state ($v_R$), $\tau_L$ is the time in the left-oriented state ($v_L$). Thus, we obtain a RnT process with different velocities (the orange line in fig. S16(c)). We neglect the last region because it finishes as the agent reaches the boundaries and does not correspond to a tumble.

Then, we average the times $\tau_{R/L,i}$ and calculate the tumble rates $\alpha_{R,L} = \langle \tau_{R/L,i} \rangle^{-1}$. Also, we average the positive and negative median velocities with weights, which correspond to the time $\tau_{R,L}$ between tumbles (the green lines in fig. S16(c)):

$$v_R = \frac{\sum_i v_{R,i}\tau_{R,i}}{\sum_j \tau_{R,j}}; \qquad v_L = \frac{\sum_i v_{L,i}\tau_{L,i}}{\sum_j \tau_{L,j}}$$

In the case of the RnT process with stops, we introduce additional pauses at the tumble events. We divide each fragment of the smoothed velocity curve between the zeros into three regions (fig. S15(c)1). The first region refers to acceleration, the second region is approximated by linear motion, and the third region refers to deceleration. We repeat this procedure for each fragment of trajectory excluding the last one where the DV reaches the threshold.

We consider one fragment between zeros. In the first region, we estimate the mean time of acceleration $\tau_a$, where the absolute value of velocity $V$ increases from zero to a certain value $V^*$, which depends on the median velocity in this fragment: $V^* = 0.9 \times \text{median}(V)$. In the second, we assume a persistent motion, where V oscillates around the MF solution. In the third part, we calculate the mean time of deceleration $\tau_d$, where the absolute value of $V$ decreases from $V^*$ to zero.

The pauses in the RnT process correspond to the regions of acceleration and deceleration. Thus, we determine the average duration of stops: $t_{\text{stop}} = \tau_a + \tau_b$. Then, we estimate the RnT velocities $v_{R,L}$ in each interval between the pauses, as was described above for the regular RnT without stops.

From the analysis of the trajectories at different points of the IIM phase diagram within the ordered phase, we find that the IIM dynamics can be approximated by different versions of the RnT motion: symmetric ($v_R \approx v_L$) or asymmetric ($v_R \neq v_L$), without stops ($t_{\text{stop}} \ll \tau_{R,L}$) or with stops. We also noticed that in the biased case ($\epsilon_1 > 0$), both the tumbling rates and the velocities are different: $\alpha_R > \alpha_L$, $v_R > v_L$.

#### 2. *Run-and-tumble without stops*

We aim to analyze the decision properties in the IIM using the RnT model. Therefore, we find the analytical expressions for the proportion of correct and wrong choices and the mean first-passage time (MFPT) to reach the ends of the interval in the RnT motion without stops.

We consider a particle that starts with $v_R$ or $-v_L$ with equal probabilities and moves as given in eq. (S35). To determine the first-passage properties of the RnT process, we first construct the forward master equations for the

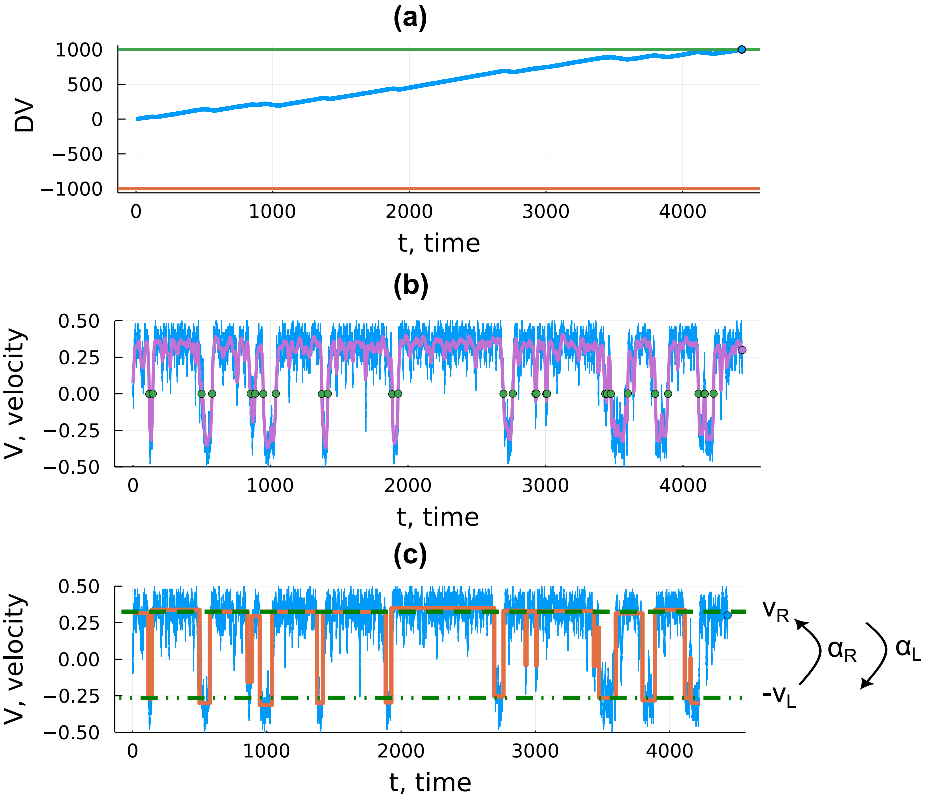


Figure S16: (a) The typical trajectory of the decision variable (DV) and the velocity dynamics in the IIM at $T = 0.36$, $\eta = 0.33$, $\epsilon_1 = 0.035$ (the ordered phase) for a high threshold value: $L = 1000$, same for both options. Initially, all spins are in the resting state (IC: ZERO). (b) Velocity dynamics in the IIM. The purple line indicates the averaged velocity in a sliding window of the length of 250 (time steps). The green circles denote the zeros at which a tumble occurs. (c) Velocity dynamics in the IIM. The orange line indicates the approximated RnT process. The green dashed lines denote the average velocities $v_R$ (positive direction) and $v_L$ (negative direction). $\alpha_{R,L}$ are the tumble rates.

probability densities $P_{L,R}(x, t|x_0, 0)$ to find the particle at position $x$ at time $t$ if it started at $x_0$ at time $t_0 = 0$ with velocity $v_R$ or $-v_L$:

$$\begin{cases} \partial_t P_R(x,t|x_0,0) + v_R \partial_x P_R(x,t|x_0,0) = +\alpha_R P_L(x,t|x_0,0) - \alpha_L P_R(x,t|x_0,0) \\ \partial_t P_L(x,t|x_0,0) + (-v_L)\partial_x P_L(x,t|x_0,0) = +\alpha_L P_R(x,t|x_0,0) - \alpha_R P_L(x,t|x_0,0) \end{cases} \tag{S36}$$

with the initial condition $P(x, t = 0|x_0, 0) = \delta(x - x_0)$, where $P = [P_R + P_L]/2$ is the total probability of finding the particle.

To estimate the error rate in the IIM, we first find the analytical expressions for the splitting probabilities $\epsilon^{\pm}_{L,R}$, which indicate the probability for the particle to leave the system through the left end $x = -L$ or the right end $x = L$ of the interval starting at $x_0$ with velocities $v_{L,R}$ (L, R: moving to the left or right). These quantities obey the following coupled backward equations [5, 10, 12–15]:

$$\begin{cases} v_R \cdot \nabla_{x_0} \epsilon^{\pm}_R(x) + \alpha_L(\epsilon^{\pm}_L - \epsilon^{\pm}_R) = 0 \\ -v_L \cdot \nabla_{x_0} \epsilon^{\pm}_L(x) + \alpha_R(\epsilon^{\pm}_R - \epsilon^{\pm}_L) = 0 \\ \epsilon^{+}_L(-L) = 0, \quad \epsilon^{+}_R(L) = 1 \\ \epsilon^{-}_L(-L) = 1, \quad \epsilon^{-}_R(L) = 0 \end{cases} \tag{S37}$$

where the boundary conditions indicate that the particle eventually leaves the system through the left ('–') end if it started at $x_0 = -L$ with $-v_L$ or through the right ('+') end if it started at $x_0 = L$ with $v_R$.

For a positive bias, $\alpha_R > \alpha_L$ or $v_R > v_L$, the positive threshold $x = +L$ denotes the correct option, while the negative threshold $x = -L$ implies the wrong alternative. Therefore, we use $\epsilon^- = [\epsilon^-_R(x_0 = 0) + \epsilon^-_L(x_0 = 0)]/2$ to assess the error rate in the decision process.

To calculate the mean RT in the IIM, we first derive the unconditioned MFPT $T_{L,R}$ for the particle to leave the interval at any end (right/left) starting at point $x_0$ with velocity $\pm v_{L,R}$ (left-oriented or right-oriented), which obeys the backward master equations:

$$\begin{cases} v_R \cdot \nabla_{x_0} T_R(x_0) + \alpha_L(T_L(x_0) - T_R(x_0)) = -1 \\ -v_L \cdot \nabla_{x_0} T_L(x_0) + \alpha_R(T_R(x_0) - T_L(x_0)) = -1 \\ T_L(-L) = 0, \quad T_R(L) = 0 \end{cases} \tag{S38}$$

where the boundary conditions indicate that the particle is absorbed if it starts at either end of the interval with the appropriate orientation.

The RTs in the correct and wrong decisions can be estimated by using the conditioned MFPTs $T^{\pm}_{L,R}$, which indicate the mean exit time for a particle to leave the interval through the specific site ('+/–': $x = \pm L$) starting at point $x_0$ with velocity $v_R$ or $-v_L$ (R, L), which obey

$$\begin{cases} v_R \cdot \nabla_{x_0} \left[\epsilon_R(x_0) T_R(x_0)\right]^{\pm} + \alpha_L \left[\epsilon_L(x_0) T_L(x_0)\right]^{\pm} - \alpha_L \left[\epsilon_R(x_0) T_R(x_0)\right]^{\pm} = -\epsilon^{\pm}_R(x_0) \\ -v_L \cdot \nabla_{x_0} \left[\epsilon_L(x_0) T_L(x_0)\right]^{\pm} + \alpha_R \left[\epsilon_R(x_0) t_R(x_0)\right]^{\pm} - \alpha_R \left[\epsilon_L(x_0) T_L(x_0)\right]^{\pm} = -\epsilon^{\pm}_L(x_0) \\ \left[\epsilon_R T_R\right]^{\pm}(L) = 0, \quad \left[\epsilon_L T_L\right]^{\pm}(-L) = 0 \end{cases} \tag{S39}$$

where we utilize the fact that $T^{+}_{R}(x = L) = 0$ and $T^{-}_{L}(x = -L) = 0$ to find the exact analytical solutions. Since the initial velocity is random, we use $T = [T_R + T_L]/2$ at the initial position $x_0 = 0$ to assess the mean RT in the IIM, while $T^{\pm} = [T^{\pm}_R + T^{\pm}_L]/2$ at $x_0 = 0$ correspond to the RTs in the correct and wrong decisions (fig. S14(b)):

$$\begin{cases} \epsilon^-(x=0) = \dfrac{\alpha_L v_L e^{\frac{2\alpha_L L}{v_R}} - \frac{1}{2}(\alpha_L v_L + \alpha_R v_R) e^{\frac{\alpha_L L}{v_R} + \frac{\alpha_R L}{v_L}}}{\alpha_L v_L e^{\frac{2\alpha_L L}{v_R}} - \alpha_R v_R e^{\frac{2\alpha_R L}{v_L}}} \\[2ex] T(x=0) = \dfrac{(2L(\alpha_L+\alpha_R)+v_L+v_R)\left(e^{\frac{\alpha_R L}{v_L}} - e^{\frac{\alpha_L L}{v_R}}\right)\left(\alpha_R v_R e^{\frac{\alpha_R L}{v_L}} - \alpha_L v_L e^{\frac{\alpha_L L}{v_R}}\right)}{2(\alpha_L v_L - \alpha_R v_R)\left(\alpha_L v_L e^{\frac{2\alpha_L L}{v_R}} - \alpha_R v_R e^{\frac{2\alpha_R L}{v_L}}\right)} \\[2ex] T^+(x=0) = \dfrac{e^{\frac{\alpha_R L}{v_L} - \frac{\alpha_L L}{v_R}}\left(\alpha_L v_L e^{\frac{2\alpha_L L}{v_R} - \frac{2\alpha_R L}{v_L}} - \alpha_R v_R\right)}{2v_R(\alpha_L v_L - \alpha_R v_R)\left(\alpha_L v_L e^{\frac{2\alpha_L L}{v_R}} - \alpha_R v_R e^{\frac{2\alpha_R L}{v_L}}\right)^2} \left\{ \dfrac{e^{\frac{\alpha_R L}{v_L}}}{\alpha_L v_L e^{\frac{\alpha_L L}{v_R}} - \alpha_R v_R e^{\frac{\alpha_R L}{v_L}}} \left[ \alpha_L \alpha_R v_R e^{2L\left(\frac{\alpha_L}{v_R} + \frac{\alpha_R}{v_L}\right)} \times \right.\right. \\ \quad \times \left(\alpha_L L v_L (v_L + 2v_R) + \alpha_R L v_R (2v_L + v_R) + 2v_L v_R (v_L + v_R)\right) - \alpha_R^2 L v_R^3 (\alpha_L + \alpha_R) e^{\frac{\alpha_L L}{v_R} + \frac{3\alpha_R L}{v_L}} + \\ \quad \left. \alpha_L^2 L v_L e^{\frac{4\alpha_L L}{v_R}} \left(\alpha_L v_L^2 + \alpha_R v_R^2\right) - \alpha_L \alpha_R v_R e^{\frac{3\alpha_L L}{v_R} + \frac{\alpha_R L}{v_L}} \left(\alpha_L L v_L (2v_L + v_R) + \alpha_R L v_R (v_L + 2v_R) + 2v_L v_R (v_L + v_R)\right) \right] \\ \quad + \dfrac{e^{\frac{\alpha_L L}{v_R}}}{v_L\left(e^{\frac{\alpha_L L}{v_R} - \frac{\alpha_R L}{v_L}} - 1\right)} \left[ \alpha_R v_R e^{\frac{\alpha_L L}{v_R} + \frac{2\alpha_R L}{v_L}} \left(\alpha_L L v_L (v_L + 2v_R) + \alpha_R L v_R (2v_L + v_R) + v_L v_R (v_L + v_R)\right) + \right. \\ \quad + \alpha_L v_L e^{\frac{3\alpha_L L}{v_R}} \left(\alpha_L L v_L^2 + v_R(\alpha_R L v_R + v_L(v_L + v_R))\right) - \alpha_R v_L v_R^2 e^{\frac{3\alpha_R L}{v_L}} \left(L(\alpha_L + \alpha_R) + v_L + v_R\right) - \\ \quad \left.\left. - \alpha_L v_L e^{\frac{2\alpha_L L}{v_R} + \frac{\alpha_R L}{v_L}} \left(\alpha_L L v_L (2v_L + v_R) + \alpha_R L v_R (v_L + 2v_R) + v_L v_R (v_L + v_R)\right) \right] \right\} \\[2ex] T^-(x=0) = \dfrac{1}{2(\alpha_L v_L - \alpha_R v_R)} \left[ 2\alpha_L L + 2\alpha_R L + v_L + v_R + \dfrac{L(v_L+v_R) e^{\frac{\alpha_R L}{v_L}} (\alpha_L v_L + \alpha_R v_R)}{v_L v_R \left(e^{\frac{\alpha_R L}{v_L}} - e^{\frac{\alpha_L L}{v_R}}\right)} + \right. \\ \quad \left. + \dfrac{\alpha_R (v_L+v_R) e^{\frac{\alpha_R L}{v_L}} (\alpha_L L v_L + \alpha_R L v_R + 2v_L v_R)}{v_L \left(\alpha_R v_R e^{\frac{\alpha_R L}{v_L}} - \alpha_L v_L e^{\frac{\alpha_L L}{v_R}}\right)} + \dfrac{4\alpha_R (v_L+v_R) e^{\frac{2\alpha_R L}{v_L}} (\alpha_L L v_L + v_R(\alpha_R L + v_L))}{v_L \left(\alpha_L v_L e^{\frac{2\alpha_L L}{v_R}} - \alpha_R v_R e^{\frac{2\alpha_R L}{v_L}}\right)} \right] \end{cases} \tag{S40}$$

In the case of small bias, we can describe or system with the symmetric RnT process with equal velocities: $v_L = v_R = v$. Then, the analytical solutions are as follows (fig. S14(c))

$$\begin{cases} \epsilon^-(x=0) = \frac{1}{2} \dfrac{(\alpha_L+\alpha_R) e^{\frac{L(\alpha_L-\alpha_R)}{v}} - 2\alpha_L e^{\frac{2L(\alpha_L-\alpha_R)}{v}}}{\alpha_R - \alpha_L e^{\frac{2L(\alpha_L-\alpha_R)}{v}}} \\[2ex] T(x=0) = \dfrac{\left(1 - e^{\frac{L(\alpha_L-\alpha_R)}{v}}\right)\left(\alpha_R - \alpha_L e^{\frac{L(\alpha_L-\alpha_R)}{v}}\right)(L(\alpha_L+\alpha_R)+v)}{v(\alpha_R-\alpha_L)\left(\alpha_R - \alpha_L e^{\frac{2L(\alpha_L-\alpha_R)}{v}}\right)} \\[2ex] T^-(x=0) = T^+(x=0) = \dfrac{1}{v(\alpha_L-\alpha_R)} \left[ \dfrac{\alpha_R(L(\alpha_L+\alpha_R)+2v)}{\alpha_R - \alpha_L e^{\frac{L(\alpha_L-\alpha_R)}{v}}} - \dfrac{L(\alpha_L+\alpha_R)}{e^{\frac{L(\alpha_L-\alpha_R)}{v}} - 1} - \dfrac{4\alpha_R(L(\alpha_L+\alpha_R)+v)}{\alpha_R - \alpha_L e^{\frac{2L(\alpha_L-\alpha_R)}{v}}} + L(\alpha_L+\alpha_R) + v \right] \end{cases} \tag{S41}$$

We analyze the results by plotting the solutions for the error rate ($\epsilon^-$), the conditioned and the unconditioned MFPTs ($T$, $T^{\pm}$) for both symmetric and asymmetric RnT processes (fig. S14(b), (c)).

If a particle starts in the middle of the interval and moves according to the general asymmetric RnT process with unequal velocities $v_R \neq v_L$, the conditioned times are not equal for any tumble rates ($T^- \neq T^+ \neq T$). In the symmetric RnT process with $v_R = v_L$, the conditioned times are equal ($T^- = T^+$), even for the unequal tumble rates ($\alpha_R \neq \alpha_L$), similarly to the DDM. Also, the conditioned times $T^{\pm}$ are equal to the unconditioned time $T$ only in the limit of diffusion [16] (large equal tumble rates $\alpha_{R,L} = \alpha \to \infty$ and a large speed $v_{R,L} = v \to \infty$ at a constant diffusivity $D = v^2/\alpha$).

In the case of unequal velocities with the drift towards the positive threshold ($v_R > v_L$) the conditioned time to reach $x = L$ is lower ($T^+ < T^-$), for any relation between the tumble rates $\alpha_{R,L}$. These results show the same tendency as the ratio $\mathrm{RT_c/RT_w}$ in IIM does at low temperatures in zone II (fig. S10(c)2), where the particle moves ballistically or makes a few tumbles (with low tumble rates), or at high temperatures in zone II, where the tumble rates are high and the regime is close to the drift-diffusion. However, the analytical predictions for the regular RnT motion without stops cannot explain the peak above 1 in the middle of zone II.

#### *3. Run-and-tumble with stops*

In order to describe the correspondence between the IIM and RnT in zone II qualitatively, we modify the RnT process by adding stops at the spin-flip events (as described in section S3 D 1 and shown in fig. S15(c)) in order to describe qualitatively the correspondence between the IIM and RnT in zone III (fig. S10(c)2). In other words, we consider the one-dimensional Levy walk with constant pauses [17].

Looking at the numerical simulations of the symmetric RnT with stops ($v_R = v_L = v$, fig. S14(d)), we find that the mean RT in the correct decisions surpasses the RT in the wrong trajectories if the particle starts in the middle of the interval.

We explain these observations as follows. As the temperature and inhibition decrease, the tumble rates also decrease, allowing fewer tumbles to occur. Therefore, at low temperatures of zone II, the set of trajectories consists of linear trajectories and trajectories with a few tumbles. Meantime, due to the positive bias $\epsilon_1$ in the IIM, we get $a_R > a_L$, which allows more tumbles to occur towards the positive direction. Therefore, the particle has longer trajectories (on average) while reaching the positive threshold $x = L$ compared to the negative threshold $x = -L$. This result becomes even more prominent as we take into account the process of acceleration and deceleration at each tumble, as it increases the duration of long trajectories at each flipping event. Thus, at moderate temperatures of zone II, the effect of long trajectories prevails, and the RT ratio reaches its maximum above 1 for both IIM and RnT processes (fig. S10(c)2). The simulations necessarily under-sample long trajectories that reach the negative threshold, as these become rare, and therefore, the simulations do not recover the theoretical prediction of equal conditioned exit times, even for pure RnT.

### S4. ISING DECISION MAKER WITH GLOBAL INHIBITION

This section discusses the Ising Decision Maker (IDM) and its properties [9] (see fig. 3B in the main text).

The IDM describes the same spin system as the Integrated Ising model (IIM) does (see sec. “Theoretical model” in the main text). The difference in the models is in the way of integration and the decision rule. The IIM’s DV integrates the relative firing activity in the two spin groups (with instantaneous velocity $V = n_1 - n_2$) similar to the drift-diffusion model [8, 18].

At the same time, in the IDM, the decision variable (DV) is represented by two components which are the instantaneous firing activity in the two groups ($n_{1,2}$). The decision process starts at zero when both groups are inhibited and continues until one group reaches a high-activity state, while the second group is inhibited. Therefore, the decision-making process in the IDM resembles the gradient descent on the two-dimensional energy surface that we show in fig. S17(a)1. The decision thresholds are marked as boxed around the energy minima [9] (green lines in fig. S17(a)1). We also show the evolution of the firing activities in the two groups (fig. S17(a)2,3) and the velocity and DV, defined as in the IIM ($V = n_1 - n_2$, $DV = \int V(t)dt$), for the direct comparison of the two models (fig. S17(a)4,5).

One can consider the IDM as a limit of the IIM with a low threshold, which does not allow switching between the two stable states (the MF solutions $V^{\pm}_{\mathrm{MF}}$ of eq. (7) in the main text). However, this interpretation is relevant for the ordered phase of the IIM. In the disordered phase, the neural firing activity is limited, and the spin system remains in the state of “indecision”. As a result, the trajectories do not reach the fixed decision thresholds (fig. S17(b)). Therefore, the phase space of the IDM is confined by the second-order transition line.

The error rate and the RT in the IDM (fig. S18) behave similarly to the IIM in the ordered phase (fig. 2B-D in the main text). At fixed bias, the error rate is smaller near the second-order phase transition line, while the RT grows with temperature $T$ and inhibition $\eta$. However, the RT ratio in the correct and wrong decisions behaves differently, reaching its minimum (below 1) near the tricritical point. Also, the ratio $\mathrm{RT_c/RT_w}$ in the IDM does not have a maximum above 1 as the IIM exhibits (zone II in fig. S10(c)2).

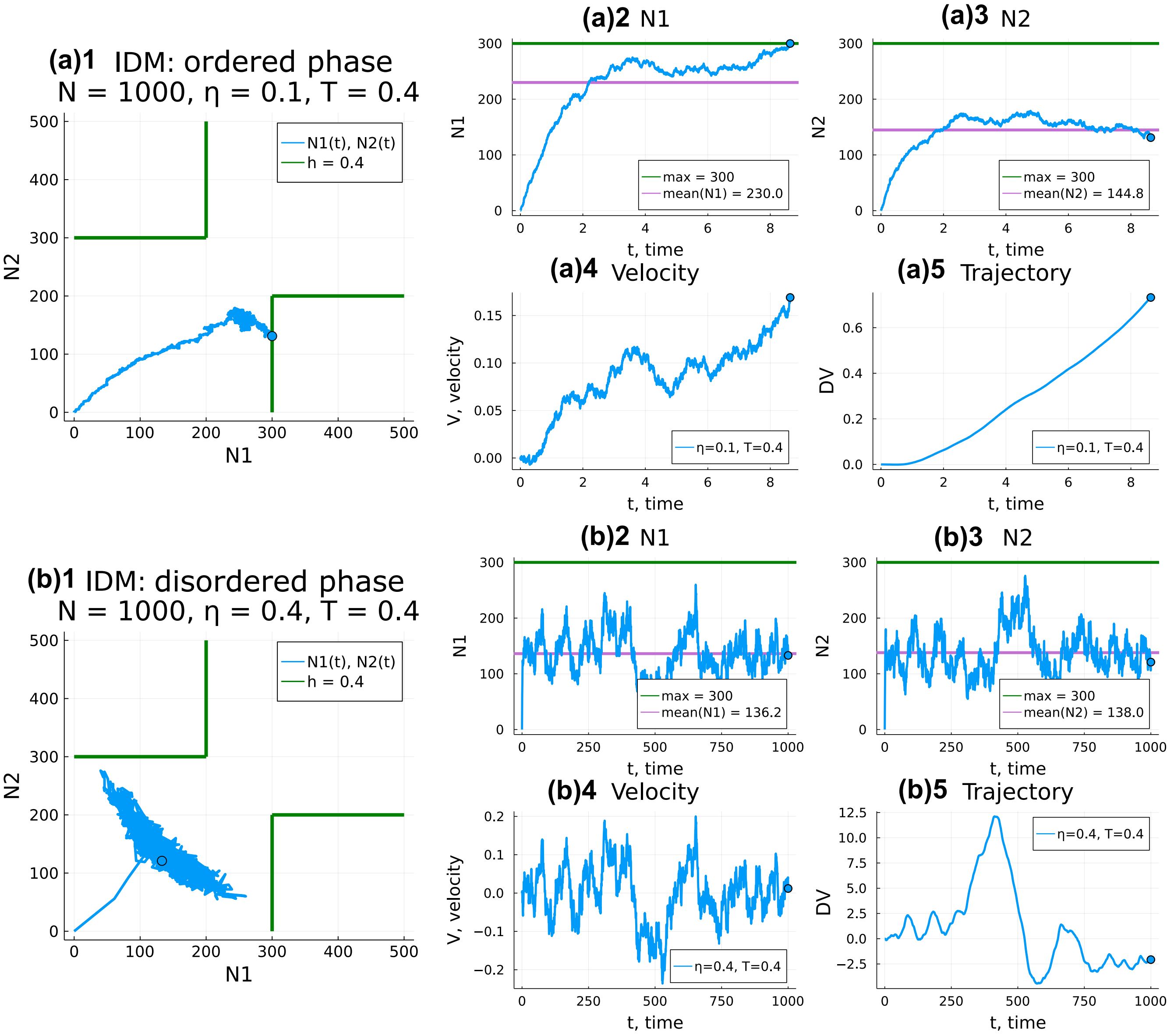


Figure S17: IDM: typical trajectories in the ordered and disordered phases. (a) Ordered phase: $T = 0.4$, $\eta = 0.1$, $\epsilon_1 = 0$. (a)1 Activity in the two spin groups. The total number of spins: $N = 1000$. The number of spins in the groups: $N^I = N^{II} = 500$. The thresholds are represented as boxes of the size $hN^I \times hN^{II}$, where $h = 0.4$. (a)2 Activity in group $I$ as a function of time. The green line indicates the lower side of the box around the energy minimum. The purple line is the average firing activity. (a)3 Activity in group $II$ as a function of time. (a)4 Velocity in the decision process, defined as in the IIM ($V = n_1 - n_2$). (a)5 Decision variable (DV) in the decision process, defined as in the IIM ($DV = \int V(t)dt$). (b) Disordered phase: $T = 0.4$, $\eta = 0.4$, $\epsilon_1 = 0$.

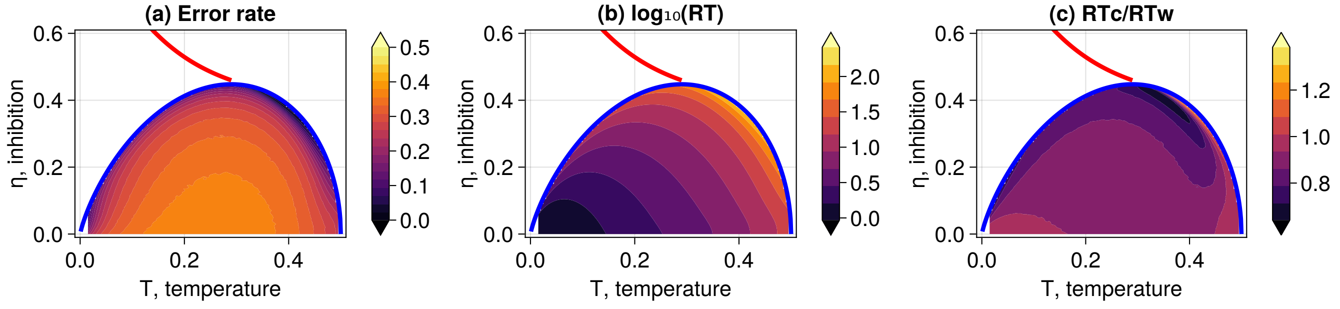


Figure S18: IDM [9]. (a) Error rate, (b) Reaction time (RT), (c) The RT ratio in the correct and wrong decisions as functions of the system's parameters ($\eta$, $T$) at fixed bias $\epsilon_1 = 0.01$ for the zero IC, presented as the heatmaps (the color bars indicate the values). The decision thresholds are defined as shown in fig. S17(a)1. The red and blue lines on the heatmaps denote the first and second-order transitions in the IIM, respectively.

### S5. SPIN ACTIVITY IN THE IIM.

Motivated by the experimental results shown below, we now explore the properties of the IIM with respect to learning, memory decay, and neuronal activity as a function of the global inhibition. We focus on the region near the tricritical point by fixing the temperature at $T = 0.3$ and varying the inhibition (fig. S19).

Within our model, the bias represents the result of a learning process by which the subject learns from experience over repeated trials which of the options is correct. The bias in our model represents the strength of this conviction regarding the correct option and thereby determines the accuracy of the decisions. Irrespective of the precise representation of the learning process, we can explore the properties of the IIM for different biases. Since the accuracy of the decisions is a measurable quantity, unlike the bias, we can compare the behavior of the IIM at different inhibitions if we maintain the same accuracy.

For example, in fig. S19(a)1,(b)1, we consider the following learning process: starting with low bias that corresponds to a fixed level of high error (0.37, blue line), we find the high biases that correspond to a low error, which represent the end of the learning process (0.08, green line). In fig. S19(a)2, we plot the ratio of the biases during this learning process and find that this is maximal near the transition line. This observation indicates that in this region, it takes the largest relative increase in bias to achieve the same improvement in accuracy.

The flip-side of this property is demonstrated in fig. S19(a)1,(b)2: here we use the high biases achieved at the end of the previous learning process (green line), and consider a process of memory decay which is represented by the decay of the bias by a fixed arbitrary factor of 4 (orange line in fig. S19(a)). We find that the relative increase in error due to this bias decay is minimal near the transition line (fig. S19(b)2).

Related to this maximal increase in relative bias (fig. S19(a)2), we find that the relative increase in the speed of decision making is maximal at the transition line, as observed by the largest decrease in the RT (fig. S19(c)).

In the IIM, we can relate the spin states to the neuronal firing activity and see how it depends on the global inhibition. Therefore, we can make some predictions using our model with respect to the neuronal activity during decision making, which is measurable [19].

In fig. S19(d)1, we plot the overall activity of the neurons, which we define to be simply related to the fraction of active spins: $A = (N_1^I + N_1^{II})/N$, as a function of the global inhibition and for the different biases given in fig. S19(a)1. We find that near the transition, there is a maximum in the relative increase in activity (fig. S19(d)2). This result, together with fig. S19(a)2, indicates that in this region, the relative change in bias and activity required to improve the accuracy of decision making is maximal.

The theoretical basis for the appearance of the maxima and minima in fig. S19 remains a challenge that will be addressed in future work. It may be that these quantities are related to the susceptibility of the system to an external field, which we know is maximal near a 2nd-order phase transition [20].

We also note that the region close to the tricritical point is least affected by fluctuations in the strength of the cross-inhibition between the spin groups, as shown in SI section S1. This may, therefore, be an additional advantage of this region of parameter space, allowing the decision-making process to be more robust against fluctuations in the interactions between the neurons.

The learning process in this region is most "costly", involving the largest relative increase in bias and neuronal activity, but it has the advantage of giving the largest relative increase in the decision speed, while the accuracy of the decisions is least sensitive to the decay in the bias.

[t!]

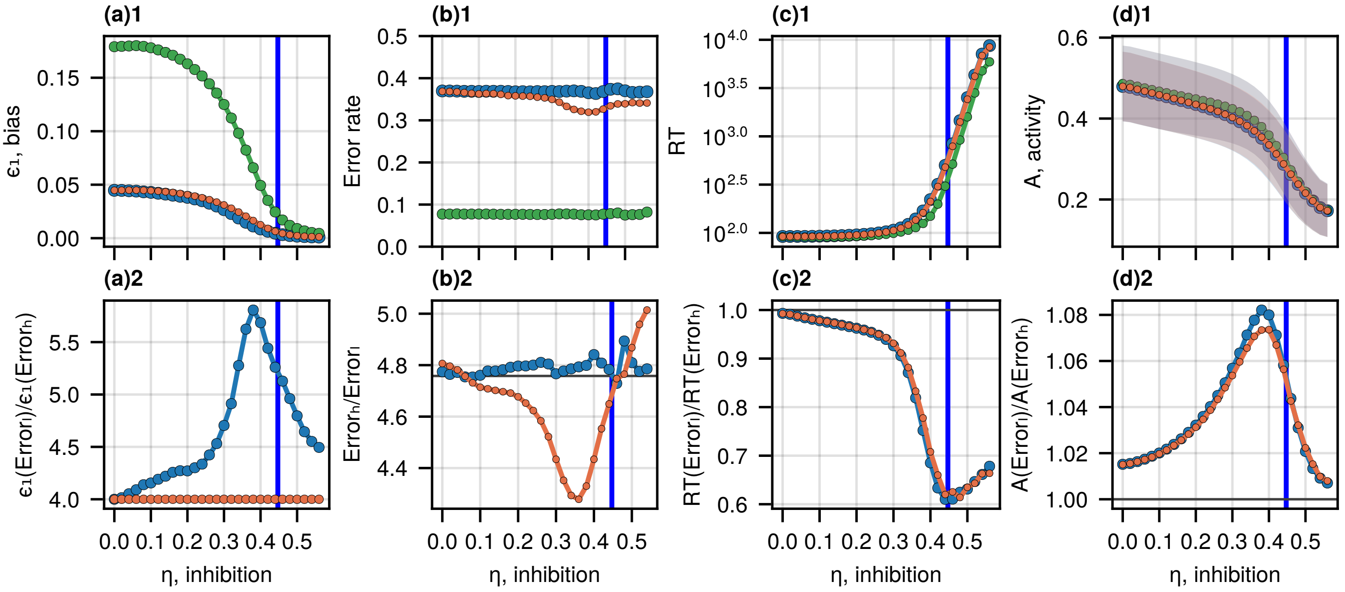


Figure S19: Spin activity in the IIM. (a)1 The biases corresponding to the fixed error rates: $0.08 \pm 0.01$ (green line) and $0.37 \pm 0.02$ (blue line) as functions of the global inhibition $\eta$ at a fixed temperature $T = 0.3$. The orange line denotes the green line divided by 4. The blue vertical line indicates the second-order phase transition. (a)2 The ratio of the green/blue (blue line) and of the green/orange biases (orange line) from (a)1, as a function of the global inhibition $\eta$. (b)1 The fixed error rates of $0.08 \pm 0.01$ (green line) and $0.37 \pm 0.02$ (blue line), corresponding to the biases with the same colors in (a)1. The orange line gives the errors corresponding to the orange bias line in (a)1. (b)2 The ratio of the green/blue (blue line) and the green/orange errors (orange line) from (b)1, as a function of the global inhibition $\eta$. (c)1 The average RTs for the three biases in (a)1 as functions of the global inhibition $\eta$. (c)2 The ratio of the RTs (green/blue and green/orange) from (c)1. (d)1 The total firing activity of spins for the different biases shown in (b)1, which is the total fraction of active spins in the two groups ($A = (N_1^I + N_1^{II})/N$), as functions of the global inhibition $\eta$ (mean $\pm$ STD). (d)2 The ratio of the firing activities (green/blue and green/orange) from (d)1.

### S6. EXPERIMENTAL SETUP I

#### A. Data measurements

We consider the process of reinforced learning in the context of a probabilistic game, where the subject learns over repeated trials which of the presented options is the correct one. The correct option is the one that gives a reward at a higher probability.

In this experimental setup, the game is represented by a sequence of two-choice decision tasks with two pairs of two options (Chinese characters), each pair refers to the gain or loss conditions. One option in each pair gives a score of 1 (gain) or -1 (loss) with a probability of 70% or 0 with a probability of 30%, and the other option gives a reward of ±1 (gain or loss condition) or 0 with the probabilities of 30% and 70%, respectively. In the gain condition, the first option, which gives +1 in 70% of trials, is the correct one since, on average, it maximizes the total score. In the loss condition, the correct option is the second option, which gives a zero reward with a probability of 70%.

In the experiment, 20 volunteers accomplished a series of two-choice tasks under uncertainty: 60 trials per condition (120 decision tasks). The choice and reaction time (RT, in ms) were registered during each trial (fig. S20(a)). The data comprises choices and the corresponding decision time in the trials per participant under the gain and loss conditions. A few trials showed significantly low RTs (3-4 ms) and therefore, were removed in the following analysis (possibly explained by technical issues in the registration process).

The initial trials imply the learning process, where the volunteers explore the game conditions and the hidden probabilities. Therefore, we consider trials 1-33 as the learning process (blue vertical line in fig. S20(b), (c)), after which the average proportion of correct choices saturates (fig. S20(b)). The number of initial trials (33) is chosen such that a random sequence of binary choices can give 30% mistakes with less than 1% probability. In other words, if the number of mistakes in the given sequence of binary choices is lower than 30%, it is highly likely that the sequence is not random (with a significance of 1%). Therefore, the following analysis excludes the choices and RTs in these learning trials.

We excluded all the participants (in both gain and loss games) who, for the given condition, did not explore both options. We required the participants to choose each option in the pair at least once during the entire game. One participant chose the wrong option in all trials after the learning period, and we also excluded these results. Overall, we analyze the results of 16 participants.

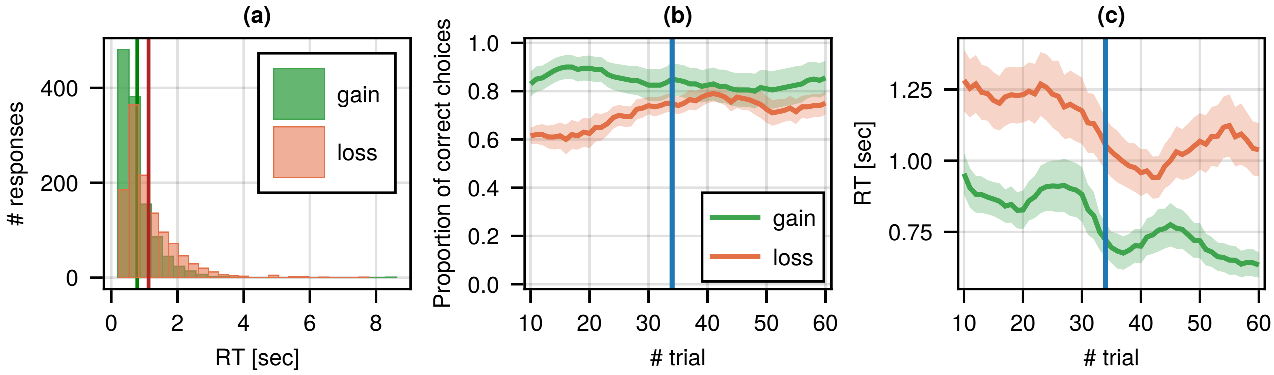


Figure S20: (a) RT distribution of all responses in trials 1-60 of all 20 participants in the gain and loss conditions. The green and red vertical lines indicate the average RTs (± STD, in sec): $0.77 \pm 0.23$ (gain), $1.13 \pm 0.37$ (loss). (b) Proportion of correct choices. (c) The average RTs. The green and orange lines represent the probability of choosing the correct option and the average RT in the 70%/30% gain (green) and loss (orange) conditions, averaged over a 10-trial moving window for all participants (mean ± SE, standard error of the mean). The blue vertical lines indicate the learning process (trials 1-33).

#### B. Data analysis

In the data set described above, we measure the error rate as the fraction of wrong choices made by each participant under the gain or loss conditions and the mean RT per participant over all trials after the learning period (fig. S21(a)). We then average the error rates and the RTs over the participants in the gain and loss conditions (the green and red lines in histograms in fig. S21(a)). We also measure the mean RT in the correct decisions ($\mathrm{RT_c}$) and the mean RT in the wrong decisions ($\mathrm{RT_w}$) per participant under each condition. Then, we take the ratio $\mathrm{RT_c/RT_w}$ and average over the participants (fig. S21(b)).

The resulting error rate, the mean ratio of the RTs in the gain and loss conditions ($\mathrm{RT_{gain}/RT_{loss}}$), and the mean RT ratio in the correct and wrong decisions ($\mathrm{RT_c/RT_w}$) under the gain and loss conditions are presented in table S3. Note that the ratios are first calculated per participant and then averaged. The quantities are calculated over 16 participants, except for the ratio $\mathrm{RT_c/RT_w}$ under the gain condition, where only 9 participants made wrong choices.

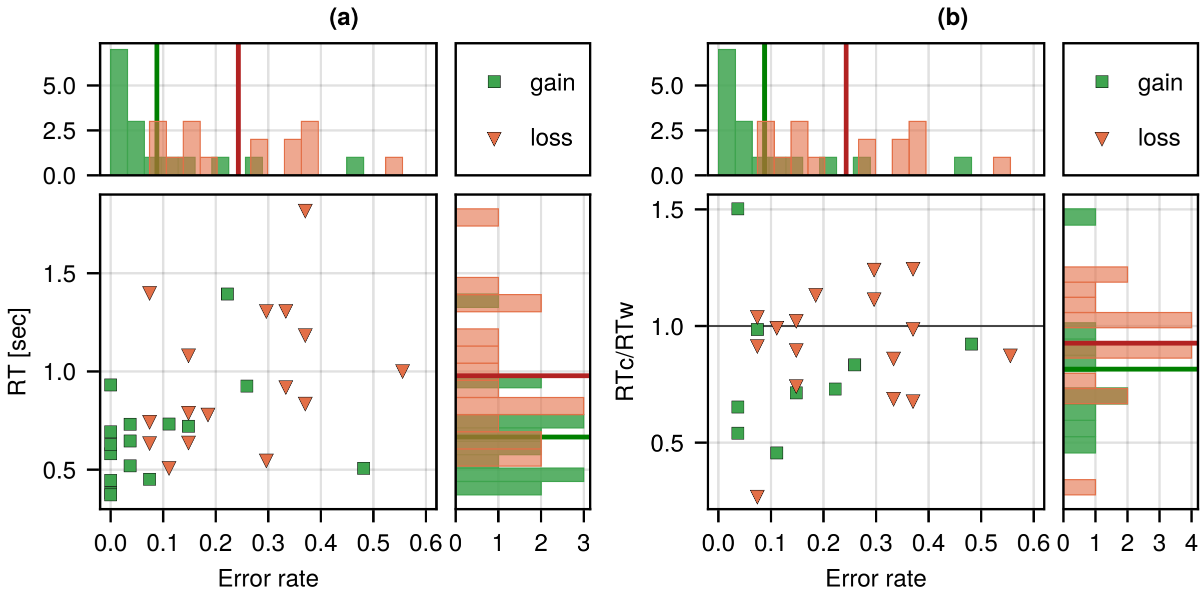


Figure S21: (a) The average RT vs. the error rate in the gain and loss condition after the learning process (bottom right). (b) The average ratio $\mathrm{RT_c/RT_w}$ vs. the error rate in the gain and loss condition after the learning process (bottom right). Each square (gain) and triangle (loss) indicate a result of one game for a single participant. Top: histogram of the error rate per participant. The green and red vertical lines indicate the average error rates of the participants (table S3). Right: histogram of the average RTs and the average ratios $\mathrm{RT_c/RT_w}$ per participant. The green and red horizontal lines indicate the average RTs and ratios (table S3). The black horizontal line in (b) is the ratio of 1.

Table S3: Experimental setup I: intermixed trials

| **Parameters** | **mean ± STD** | **mean ± SE** | **# participants** |
|---|---|---|---|
| Error rate, gain | $0.088 \pm 0.134$ | $0.088 \pm 0.033$ | 16 |
| Error rate, loss | $0.243 \pm 0.142$ | $0.243 \pm 0.035$ | 16 |
| RT (gain) [sec] | $0.666 \pm 0.258$ | $0.666 \pm 0.065$ | 16 |
| RT (loss) [sec] | $0.978 \pm 0.360$ | $0.978 \pm 0.090$ | 16 |
| $\mathrm{RT_{gain}/RT_{loss}}$ | $0.704 \pm 0.180$ | $0.704 \pm 0.045$ | 16 |
| $\mathrm{RT_c/RT_w}$ (gain) | $0.815 \pm 0.309$ | $0.815 \pm 0.103$ | 9 |
| $\mathrm{RT_c/RT_w}$ (loss) | $0.926 \pm 0.245$ | $0.926 \pm 0.061$ | 16 |

Experimental results, calculated for 16 volunteers, participated in 60 gain and 60 loss intermixed trials of two-choice tasks under uncertainty with mixed gain and loss conditions. The mean values are calculated in trials $34 - 60$ after the learning period.

To analyze the results, we assume the normal distribution of the values and conduct the two-tailed t-test for the mean values. Since the RT distributions are bounded by zero (positively skewed, fig. S20(a)), we also compare the medians to 1 and conduct the Wilcoxon signed-rank test.

The error rate in the gain conditions is significantly lower than the error rate in the loss conditions ($Error_{\mathrm{gain}} - Error_{\mathrm{loss}} \neq 1$; unequal variance t-test: $df = 30$, $t = -3.182$, $p = 0.0034$; Wilcoxon-test: $W = 10$, $p = 0.0029$).

The RT in the gain conditions is significantly lower than the RT in the loss conditions ($\mathrm{RT_{gain}/RT_{loss}} \neq 1$; t-test: $df = 15$, $t = -6.599$, $p < 0.0001$; Wilcoxon-test: $W = 0$, $p < 0.0001$).

The RT ratio in the correct and wrong decisions in the gain condition does not show a significant difference from 1 ($\mathrm{RT_c/RT_w}$(gain) $\neq 1$; t-test: $df = 8$, $t = -1.798$, $p = 0.1098$; Wilcoxon test: $W = 8$, $p = 0.0977$).

The RT ratio in the correct and wrong decisions in the loss condition does not show a significant difference from 1 ($\mathrm{RT_c/RT_w}$(loss) $\neq 1$; t-test: $df = 15$, $t = -1.2$, $p = 0.2489$; Wilcoxon test: $W = 50$, $p = 0.3755$).

The RT ratios in the correct and wrong decisions in the gain and loss conditions do not show a significant difference ($\mathrm{RT_c/RT_w}$(gain) $\neq RT_{c,l}/$ $\mathrm{RT_c/RT_w}$(loss); unequal variance t-test: $df = 13.8$, $t = -0.93$, $p = 0.3681$).

### S7. FITTING THE MODEL TO THE EXPERIMENTAL OBSERVATIONS

In our IIM, there are three main parameters (temperature $T$, global inhibition $\eta$, bias $\epsilon_1$) that significantly affect the outcomes of the decision processes (error rate, RT, the RT ratio in the correct and wrong decisions). We showed earlier the dependency of the outcomes as functions of each parameter while fixing two other parameters (fig. S7, SI section S2). It turns out that the error rate decreases if we increase any parameter ($\eta$, $T$, $\epsilon_1$), while the RT decreases only if the bias increases and increases if the temperature or global inhibition increase (fig. S7). In contrast, if we increase cross-inhibition $J_{out}$, the error rate increases, while the RT decreases (fig. S8).

Also, the temperature in the IIM ($T$) represents the noise in the neural network, while the global inhibition ($\eta$) and cross-inhibition ($J_{out}$) relate to the activity of inhibitory neurons and neural interactions and, therefore, are better adjustable by the brain. The experimental observations in setup I (table I in the main text, SI section S6) where both the error rate and the RT at the loss condition were smaller than at the gain condition support using the global inhibition, which shows a similar tendency. Thus, we are motivated to use global inhibition as the primary control parameter in the model.

Now, we want to fit the IIM's outcomes to the experimental observations (error rate, RT, and the ratio $RT_c/RT_w$). For all sets of parameters ($\eta$, $T$, $\epsilon_1$), we run numerical simulations for the decision trajectories (up to $2 \times 10^5$ per set) and define the error rate as the proportion of the trajectories that reached the negative threshold ($L = -40$) and the RT as the average duration of the trajectories. We also find the RT in the correct and wrong decisions by taking the trajectories that reached either the positive or the negative threshold. We run simulations for each set ($\eta$, $T$, $\epsilon_1$) for two types of initial conditions: zero (initially, all spins are inactive) and random (initially, the distribution of "on " and "off" spin states is random), see SI section S2 for the comparison.

Then, we fix a point on the phase space ($T$, $\eta$), and for the given value of the error rate, we extract the corresponding bias $\epsilon_1$ (we show how the bias changes as a function of the global inhibition at a fixed error rate denoted by color in fig. S22(a)). Since the error rate is measured with error bars, the extracted bias lies in some range, and we take the average value (the red horizontal line and the shaded area in fig. S22(b)). After that, we are able to calculate the RT and $RT_c/RT_w$, which relate to the chosen error rate for each point on the phase diagram.

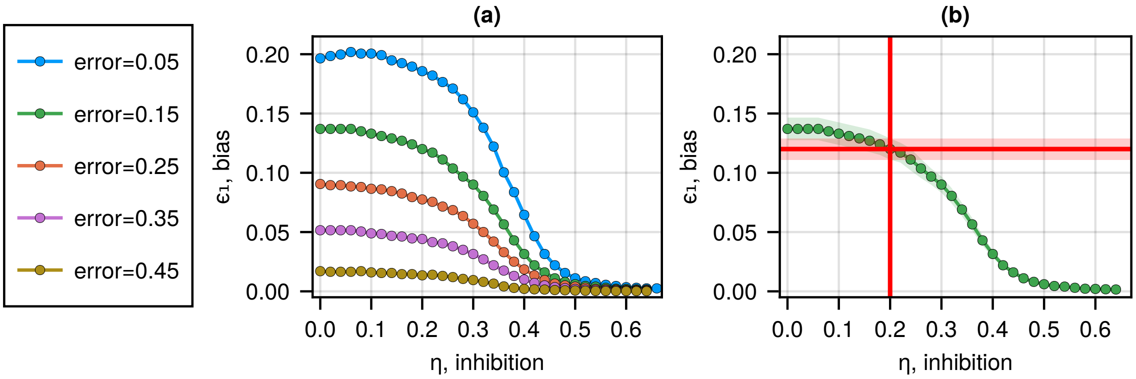


Figure S22: (a) Bias, required for a certain error rate (denoted by color), as a function of inhibition $\eta$ at a fixed temperature $T = 0.3$. (b) Bias ($\pm$ STD), required for the fixed error rate of $0.15 \pm 0.03$, as a function of inhibition $\eta$ at a fixed temperature $T = 0.3$. The vertical red line indicates the level of constant inhibition $\eta = 0.2$. The horizontal red line represents the required bias ($0.12 \pm 0.01$), which gives the desired error probability (within the error bars) at the fixed inhibition.

In the first experimental setup (section S6), the two error rates at the gain and loss condition are given together with the ratio of the two average RTs ($RT_{gain}/RT_{loss}$). For each point of the phase space ($T$, $\eta$), we find the two biases ($\epsilon_{1,gain} \neq \epsilon_{1,loss}$) corresponding to the two error rates. Then, we calculate the RT ratio ($RT_{gain}/RT_{loss}$) for each point on the phase space and the two extracted biases. We plot the Z-score of the RT ratio on the phase diagram (fig. 5C in the main text). The smallest values (green) indicate the area that fits the two error rates and is the closest to the observed $RT_{gain}/RT_{loss}$. Note that in this setup we do not fit the RT ratio in the correct and wrong decisions ($RT_c/RT_w$).

In the second experimental setup (section S9), we are given two error rates for the green and blue groups, the corresponding normalized RTs (in the biased and unbiased conditions: $RT_{65/35}/RT_{50/50}$; which is below 1 in the green group and above 1 in the blue group, see fig. 6A in the main text), and the two RT ratios in the correct and wrong decisions under the biased condition ($RT_c/RT_w$). In this scenario, we fit the error rate for the green group and find the corresponding average bias as described above. Then, we extract the normalized RT (we take $\epsilon_1 = 0$ for the $RT_{50/50}$ in the unbiased condition) and the ratio $RT_c/RT_w$ for each point on the phase space ($T$, $\eta$). We take into the following consideration only those points ($T$, $\eta$) that give the ratios $RT_{65/35}/RT_{50/50}$ and $RT_c/RT_w$ that fit the observed results for the green group. Thus, we obtain a curve near the tricritical point on the phase diagram (the green circles in fig. 6B in the main text).

Motivated by the experiment, we assume that the global inhibition in the blue group is higher than in the green group. Therefore, we shift the green curve toward higher inhibition levels by multiplying $\eta$ by a constant factor which is the ratio of the concentration of GABA molecules for the two groups, found in the experiment (see sec. “Setup II” in the main text for the details). Then, for each point $(T,\ \eta)$, we again extract the average biases that fit the error rate observed in the blue group. We find the ratio of the RT in the green and blue groups for each temperature (fig. 6C(iv) in the main text) and compare it with the ratio in the experiment ($\mathrm{RT}_{65/35}(\mathrm{green})/\,\mathrm{RT}_{65/35}(\mathrm{blue})$, indicated by the black line). It turns out that the predicted RT ratio of the average RTs in the two groups lies within the error bars of the experimental observation for the temperature near the tricritical point. Also, the curves in this setup lie close to the contour found in the previous part for the first setup (fig. 6B(iii) in the main text).

### S8. FITTING THE DDM AND RUN-AND-TUMBLE MODELS TO EXPERIMENT I

This section compares the results of the experimental observations in setup I, which are the error rates ($Error_{\rm gain}$, $Error_{\rm loss}$), the ratio of the reaction times (RT) in the gain and loss conditions ($\rm RT_{gain}/RT_{loss}$), and the RT ratio in the correct and wrong decisions ($\rm RT_c/RT_w$, in the gain or loss conditions), with the exact analytical expressions of the proportion of wrong choices ($\epsilon_g^-$, $\epsilon_l^-$), the ratio of the unconditioned mean exit times ($T_g/T_l$), and the ratios of the conditioned exit times ($T^+/T^-$, in the gain or loss conditions) for the drift-diffusion model (DDM) and the run-and-tumble motion. All the analytical derivations are presented in SI section S3. Note that we call experimental observations *Error* and *RT*, while $\epsilon^-$, $T$, and $T^\pm$ denote the analytical solutions of the DDM and the run-and-tumble motion. Also, in both models, we assume that the positive and negative thresholds are placed at equal distances $L$ from the initial position ($x_0 = 0$) of the stochastic decision processes.

#### A. RT distributions

We first aim to compare the RT distributions in the experiments with the IIM's simulations for different areas of the phase space. We choose three points in the ordered, intermittent, and disordered phases (fig. S23(a)1, fig. S23(b)1) and find the biases that satisfy the experimental error rates in the gain/loss conditions and the RT ratio in the gain and loss conditions $\rm RT_{gain}/RT_{loss}$ (as described in sec. "Setup I" in the main text and section S7). We then plot the normalized RT distributions in the experiments and in the IIM's simulations (divided by the mean RT in each distribution, fig. S23(a)2, fig. S23(b)2) and the typical decision trajectories corresponding to the diffusion process in the disordered phase and run-and-tumble or ballistic in the ordered and intermittent phases (fig. S23(a)3, fig. S23(b)3).

The differences between the experimental data (x-axis) and the three distributions (y-axis) are quantified using the quantile-quantile plot [21, 22] (fig. S23(a)4, fig. S23(b)4) and the distribution parameters (table S4). Overall, the distributions for both the disordered and intermittent phases are significantly different from the data, while the ordered phase (close to the transition) is similar to the observed data.

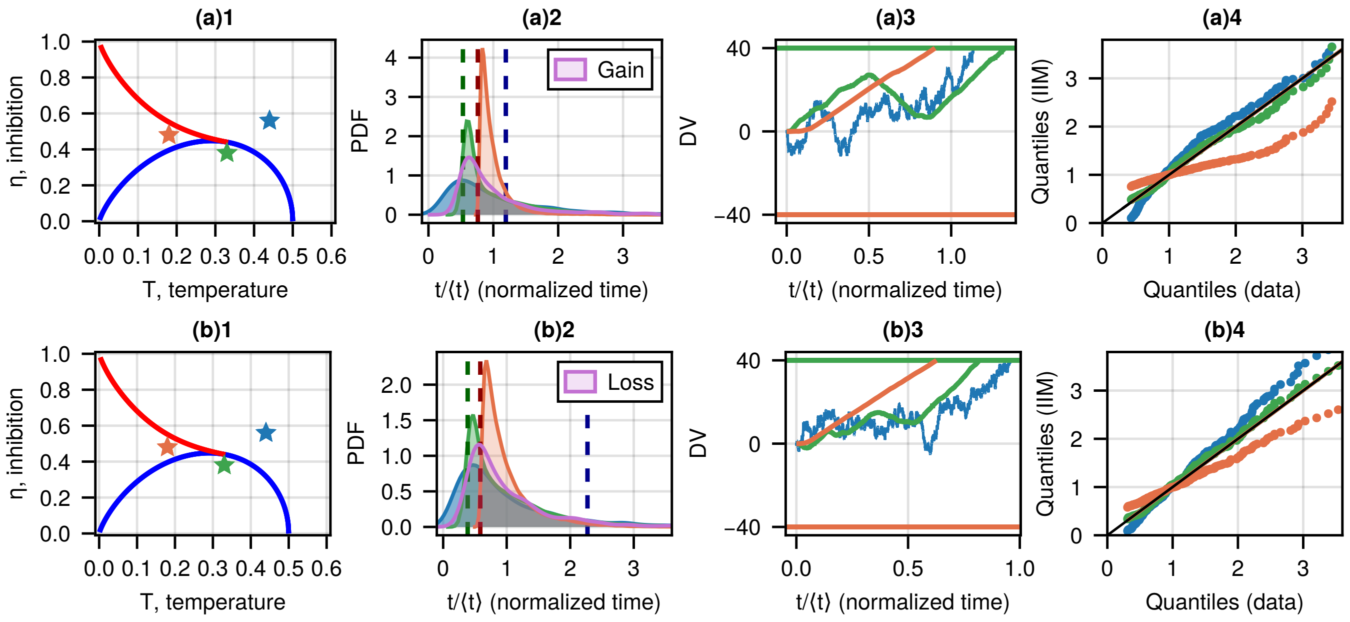


Figure S23: (a)1 Phase diagram of the IIM. The red and blue lines on the heatmaps denote the first and second-order transitions, respectively. The stars indicate the parameters ($\eta$, $T$) in the three phases from which we sample the IIM's RT distributions to compare with the experimental data. (a)2 The RT distributions (normalized by the mean RT) for the three regimes (denoted in (a)1) and the observed RT distribution in the trials after the learning period in the gain condition (purple area). The bias for each point is chosen such that the predicted error rate for this point satisfies the observed error rate for the gain condition and $\rm RT_{gain}/RT_{loss}$ (see table S4). The vertical dashed lines indicate the theoretical RT given by $RT^+_{bal}$ (eq. (8) in the main text). (a)3 Typical trajectories during the decision-making process corresponding to the IIM's RT distributions shown in (a)2. (a)4 Comparison of the IIM's RT distributions with the observed data (black identity line) using a quantile-quantile representation. (b) Comparison for the loss condition. The parameters $\eta$ and $T$ are the same as in the gain condition.

Table S4: Comparison of the parameters of the normalized RT distributions, given in different phases, vs. experimental data (setup I).

| **Parameters** | **Mean** | **Median** | **STD** | **Skewness** | **Kurtosis** | $L/V^+_{\mathbf{MF}}$ | **# simulations** |
|---|---|---|---|---|---|---|---|
| Intermittent: $T = 0.18, \eta = 0.48, \epsilon_1 = 0.05$ | 1 | 0.92 | 0.26 | 3.41 | 16.99 | 0.76 | $10^5$ |
| Ordered: $T = 0.33, \eta = 0.38, \epsilon_1 = 0.038$ | 1 | 0.75 | 0.59 | 2.29 | 7.29 | 0.53 | $2 \times 10^4$ |
| Disordered: $T = 0.44, \eta = 0.56, \epsilon_1 = 0.005$ | 1 | 0.75 | 0.79 | 2.05 | 6.22 | 1.09 | $10^3$ |
| Gain (data) | 1 | 0.78 | 0.61 | 2.19 | 5.35 | – | 432 |
| Intermittent: $T = 0.18, \eta = 0.48, \epsilon_1 = 0.019$ | 1 | 0.86 | 0.43 | 2.46 | 9.57 | 0.58 | $10^5$ |
| Ordered: $T = 0.33, \eta = 0.38, \epsilon_1 = 0.016$ | 1 | 0.75 | 0.72 | 2.15 | 6.69 | 0.38 | $2 \times 10^4$ |
| Disordered: $T = 0.44, \eta = 0.56, \epsilon_1 = 0.002$ | 1 | 0.73 | 0.82 | 1.87 | 4.75 | 2.25 | $10^3$ |
| Loss (data) | 1 | 0.76 | 0.71 | 2.69 | 11.27 | – | 432 |

The parameters of IIM ($\eta$, $T$) are marked in fig. S23. $RT^+_{bal} = L/V^+_{\mathrm{MF}}$ (eq. (8)) indicates the theoretical prediction for the mean RT using the MF velocity.

### B. DDM

Our IIM in the disordered regime is similar to the DDM though it has time correlations for the velocity dynamics because the spin system changes its configuration and velocity via a sequence of spin flips compared to the uncorrelated changes in the DDM. In the case of large decision thresholds, where the decision trajectory consists of a large number of spin-flipping events (with a large RT), we neglect this time correlation and assume that the IIM behavior can be approximated with the regular DDM. We now compare the results of the DDM with the observed data.

For the symmetric DDM (with equal thresholds $L$ and initial position at $x_0 = 0$), we derived the error rate and the unconditioned and conditioned mean exit times as functions of the system's parameters ($v$ is a constant drift, $D$ is diffusivity, $\mathrm{Pe} = vL/D$ is the Péclet number, see SI section S3 for the details):

$$\begin{cases} \epsilon^-(x_0 = 0) = \frac{1}{e^{\mathrm{Pe}}+1} \\ T(x_0 = 0) = T^+(x_0 = 0) = T^-(x_0 = 0) = \frac{L}{v} \tanh\left(\frac{\mathrm{Pe}}{2}\right) \end{cases} \tag{S42}$$

We fix the diffusion coefficient ($D$) and the threshold ($L$) and define the drift velocity ($v$) as the only free parameter in the system, which specifies the preference for one of the options in the decision task. From the analytical solutions (eq. (S42)), we write the drift velocity as a function of the error rate:

$$\epsilon^-(x_0 = 0) = \frac{1}{1 + e^{\mathrm{Pe}}} \Rightarrow \mathrm{Pe}(\epsilon^-) = \ln\left(\frac{1}{\epsilon^-} - 1\right) \overset{\mathrm{Pe}=\frac{vL}{D}}{\Rightarrow} v(\epsilon^-) = \frac{D}{L} \ln\left(\frac{1}{\epsilon^-} - 1\right) \tag{S43}$$

Then, we derive the unconditioned mean first-passage time $T$ as a function of the error rate ($\epsilon^-$):

$$T(\epsilon^-|x_0 = 0) \overset{(S42),(S43)}{=} \frac{L^2}{D} \frac{(1 - 2\epsilon^-)}{\ln\left(\frac{1}{\epsilon^-} - 1\right)} \tag{S44}$$

Then, the analytical RT ratio in the gain and loss conditions with the fixed threshold $L$, diffusivity $D$, and variable drift $v$ is written as follows:

$$\frac{T_g}{T_l} \overset{(S42)}{=} \frac{\frac{L}{v_g} \tanh\left(\frac{\mathrm{Pe}_g}{2}\right)}{\frac{L}{v_l} \tanh\left(\frac{\mathrm{Pe}_l}{2}\right)} = \frac{\mathrm{Pe}_l \tanh\left(\frac{\mathrm{Pe}_g}{2}\right)}{\mathrm{Pe}_g \tanh\left(\frac{\mathrm{Pe}_l}{2}\right)} = \frac{(1 - 2\epsilon_g) \ln\left(\frac{1}{\epsilon_l} - 1\right)}{(1 - 2\epsilon_l) \ln\left(\frac{1}{\epsilon_g} - 1\right)} \tag{S45}$$

where $\epsilon_{g,l}$ indicate the proportion of wrong choices under the gain and loss conditions predicted by the DDM. Similarly to the error rate, the RT ratio $T_g/T_l$ is determined by only one parameter Pe.

In the experiment, $Error_{\mathrm{gain}} = 0.09 \pm 0.03$, $Error_{\mathrm{loss}} = 0.24 \pm 0.04$, $\mathrm{RT_{gain}}/\mathrm{RT_{loss}} = 0.7 \pm 0.04$, (red line in fig. S24). In the disordered region of the IIM, $T_g/T_l = 0.76 \pm 0.05$ (blue line in fig. S24, see also fig. 5B(ii) in the main text above the transition lines). When we plug the given error rates into the analytical solution of the DDM (eq. (S45)), we get the following ratio: $T_g/T_l(\epsilon_g, \epsilon_l) = 0.78 \pm 0.11$ (green line in fig. S24). It turns out that the analytical expression from the DDM coincides with the ratio, given by the IIM, as $0.76 \pm 0.05$ indeed lies in the predicted range of the DDM. The DDM's predicted range overlaps with the experimental observation within the error bars ($0.7 \pm 0.04 \cap 0.78 \pm 0.11$). Also, the DDM's approximation fits the experimental data within the error bars ($\mathrm{RT_{gain}}/\mathrm{RT_{loss}} \neq 0.78$; t-test: $df = 15$, $t = -1.676$, $p = 0.1144$; Wilcoxon-test: $W = 39$, $p = 0.1439$), see fig. S24.

In the DDM, the conditioned times are equal [7], $T^+ = T^-$, meaning that this ratio is independent of the system's parameter Pe, and the DDM with symmetric thresholds and constant drift does not explain the cases where the RT ratio in the correct and wrong decisions differs from 1 (so-called fast or slow errors [8, 9]).

### C. Run-and-tumble

In the IIM, the bias that defines the error rate and the RT in the decision processes is different in the gain and loss conditions, and therefore, it introduces an asymmetry in the run-and-tumble motion in the ordered phase below the second-order phase transition line (see the details in SI section S3). It results in variability in all four parameters of

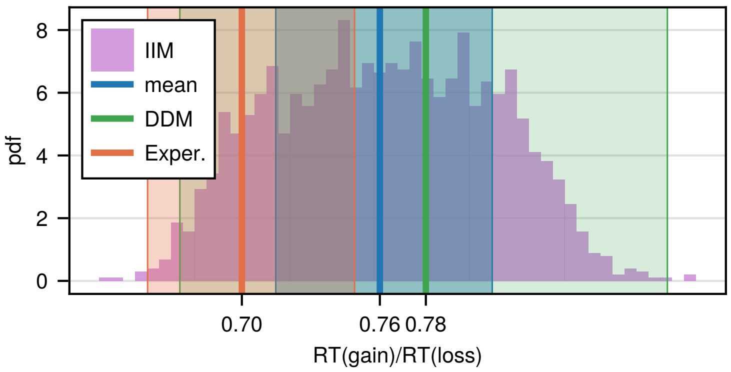


Figure S24: Comparison of the ratio $\mathrm{RT_{gain}/RT_{loss}}$ in the IIM in the disordered regime, in the traditional DDM, and in the experiment. The purple histogram indicates the distribution of the RT ratio in the gain and loss conditions obtained in the IIM in the disordered phase (see fig. 5B(ii) in the main text above the transition lines). For that, we ran simulations for each point ($\eta$, $T$) of the phase diagram above the transition line and extracted the ratio $\mathrm{RT_{gain}/RT_{loss}}$ for the points which satisfied the two errors within the error bars (table S3). The blue line with the blue-shaded region is the mean ratio ($\mathrm{RT_{gain}/RT_{loss}} \pm$ STD) in the IIM in the disordered phase ($0.76 \pm 0.05$). The green line and the green-shaded rectangle indicate the mean ratio predicted by the pure DDM ($T_g/T_l = 0.78 \pm 0.11$, see section S8 B). The orange line and the green-shaded rectangle are the mean RT ratio in the experiment ($\mathrm{RT_{gain}/RT_{loss}} = 0.7 \pm 0.04$, table S3)). This plot shows that the IIM in the disordered phase is indeed close to the analytical results of the DDM. Both the IIM and the DDM predictions for the ratio $\mathrm{RT_{gain}/RT_{loss}}$ at the two fixed errors overlap with the experiment within the error bars, indicating a marginal fit.

the RnT process ($\alpha_{R,L}, v_{R,L}$) at fixed thresholds which is more than in the IIM ($T$, $\eta$, $\epsilon_1$) and DDM ($v$, $D$).

Similarly to the DDM, we want to compare the error rates, the ratio of the RTs in the gain and loss conditions $\mathrm{RT_{gain}/RT_{loss}}$, and the RT ratio in the correct and wrong decisions $\mathrm{RT_c/RT_w}$ from the experimental observations with the exact analytical expressions of the proportion of wrong choices $\epsilon_-$ and the mean exit times $T$, $T^{\pm}$. We assume again equal thresholds and the zero initial position ($x_0 = 0$).

#### 1. *Asymmetric Run-and-tumble*

Let us demonstrate how to estimate the RT ratio in the gain and loss conditions ($T_g/T_l$) in the asymmetric run-and-tumble process by fixing three of four parameters: the tumble rate $\alpha_R$ (from left to right) and the velocity amplitudes $v_{R,L}$. The parameters are estimated at a fixed point on the phase space as described in section S3 D 1, fig. S16. In the particular case that we show below, we consider the point $T = 0.36$, $\eta = 0.38$ and bias $\epsilon_1 = 0.009$, which fits the error rate observed in the loss condition ($0.24 \pm 0.04$). We obtain $\alpha_R = 0.019$, $v_R = 0.21$, $v_L = 0.19$.

Then, we represent the bias towards the positive threshold $x = +L$ in terms of the variable ratio of the tumble rates $\tau = \alpha_L/\alpha_R$ and the fixed unequal velocities $v_R > v_L$. We plug the variable tumble rate $\alpha_L = \tau\alpha_R$ (from right to left) into the analytical solutions for the error rate $\epsilon_-$ and the mean exit time $T$ in the run-and-tumble process with equal thresholds (eq. (S40)) and obtain $\epsilon_-(\tau)$ and $T(\tau)$. Then, we find numerically two ratios $\tau_{g,l}$ as functions of the error rates $\epsilon_{g,l}$ and after that, we calculate the corresponding mean passage times $T_{g,l}$ as functions of $\epsilon_{g,l}$. Therefore, we can numerically find $T_g/T_l(\epsilon_g, \epsilon_l)$ (the procedure is similar to the DDM fitting, see eq. (S45)).

We calculate $T_g/T_l$ for all possible errors ($0 \ldots 0.5$) and display the result as a heatmap (the background in fig. S25(a)), marking the region with the experimental ratio $0.70 \pm 0.04$ (black lines) and the error rates $0.09 \pm 0.03$ (gain, green lines) and $0.24 \pm 0.04$ (loss, blue lines). As a result, in this simplified approximation, the asymmetric run-and-tumble model does fit the experimental data within the chosen error bars.

In the previous section (SI section S3), we showed that in the asymmetric run-and-tumble process without stops, the RT ratio in the correct and wrong decisions ($T^+/T^-$) is always below 1 if the velocity in the positive direction is larger than in the negative direction ($v_R > v_L$).

#### 2. *Symmetric Run-and-tumble*

We also show a more simplified version of the run-and-tumble process with equal velocity amplitudes ($v_R = v_L = v$). Extracting the run-and-tumble parameters from the IIM ($\alpha_R = 0.019$, $v = 0.21$) and considering the ratio of the tumble $\tau$ rates as the only bias in the system, we again assess the error rate and the RT ratio in the gain and loss conditions analytically to fit the experimental observations.

Similarly to the asymmetric run-and-tumble process, we find numerically the ratios of the tumble rates as functions of the error rates, $\tau_g(\epsilon_g)$ and $\tau_l(\epsilon_l)$, and derive the mean exit times $T_{g,l}$ as functions of errors $\epsilon_g$, $\epsilon_l$. We calculate $T_g/T_l$ for all possible errors ($0 \ldots\ 0.5$) and display the result as a heatmap (the background in fig. S25(b)), marking the region with the experimental ratio $0.70 \pm 0.04$ (black lines) and the error rates $0.09 \pm 0.03$ (gain, green lines) and

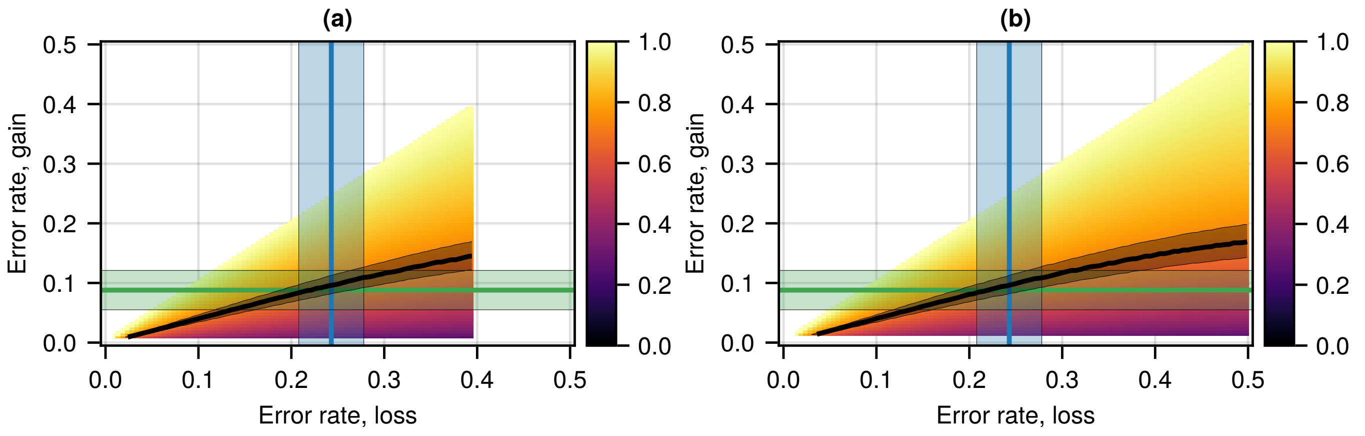


Figure S25: (a) Asymmetric run-and-tumble motion. The heatmap shows the analytical ratio of the mean first-passage times in the gain and loss condition $T_g/T_l$ as a function of the error rates in the gain and loss conditions. The green and blue lines and the shaded rectangles indicate the error rate in the gain and loss conditions from the experiment: $0.09 \pm 0.03$ and $0.24 \pm 0.04$ (table S3). The black lines indicate the RT ratio in gain and loss conditions: $0.70 \pm 0.04$. The ratio of the tumble rates $\tau = \alpha_L/\alpha_R$ plays the role of the bias in the run-and-tumble process. The threshold, the tumble rate from left to right, and the velocities are fixed: $L = 40$, $\alpha_R = 0.019$, $v_R = 0.21$, $v_L = 0.19$. The parameters are estimated from the IIM at $T = 0.36$, $\eta = 0.38$, and $\epsilon_1 = 0.009$, at the model which fits the error rate observed in the loss condition ($0.24 \pm 0.04$). (b) Symmetric run-and-tumble motion with parameters $L = 40$, $\alpha_R = 0.019$, $v = 0.21$.

$0.24 \pm 0.04$ (loss, blue lines). As a result, in this approximation, the symmetric RnT model also fits the experimental data within the error bars.

However, similar to the DDM, the symmetric run-and-tumble process with equal thresholds does not explain the case of unequal conditioned mean exit times (derived in SI section S3).

### S9. EXPERIMENTAL SETUP II

#### A. Data measurements

Similar to the first experimental setup (SI section S6), we consider the process of reinforced learning in the context of a probabilistic game. In this experimental setup, the participants completed sequences of binary decision tasks under four different conditions (see more details about the experiment and the data set in [23]).

In the gain condition, one of the two options (represented by Chinese characters) increases the total score by 1 with the fixed probability of $p$ (unknown for the participants) or does not change the total score with the probability of $1-p$. The second option increases the total score by 1 with the probability of $1-p$ and does not change it with the probability of $p$. In the loss condition, the two options (represented by other Chinese characters) decrease the total score by 1 with the probabilities $p$ or $1-p$.

The other two conditions determine the probabilities. In the unbiased case, the options are equivalent and increase (gain) or decrease (loss) the total score with the probability of $p=0.5$. In the biased case, one of the options increases (gain) or decreases (loss) the total score with the probability of $p=0.65$, while the probability for the second option is $1-p=0.35$.

In the experiment, 107 volunteers played four separate games of 50 trials in each of the following combinations: gain and loss, with probabilities of 65/35 and 50/50. The choice and the reaction time (RT, in sec) were registered during each trial (fig. S26). A few trials showed significantly low RTs (below 110 ms) and, therefore, were removed in the following analysis (possibly explained by technical issues in the registration process).

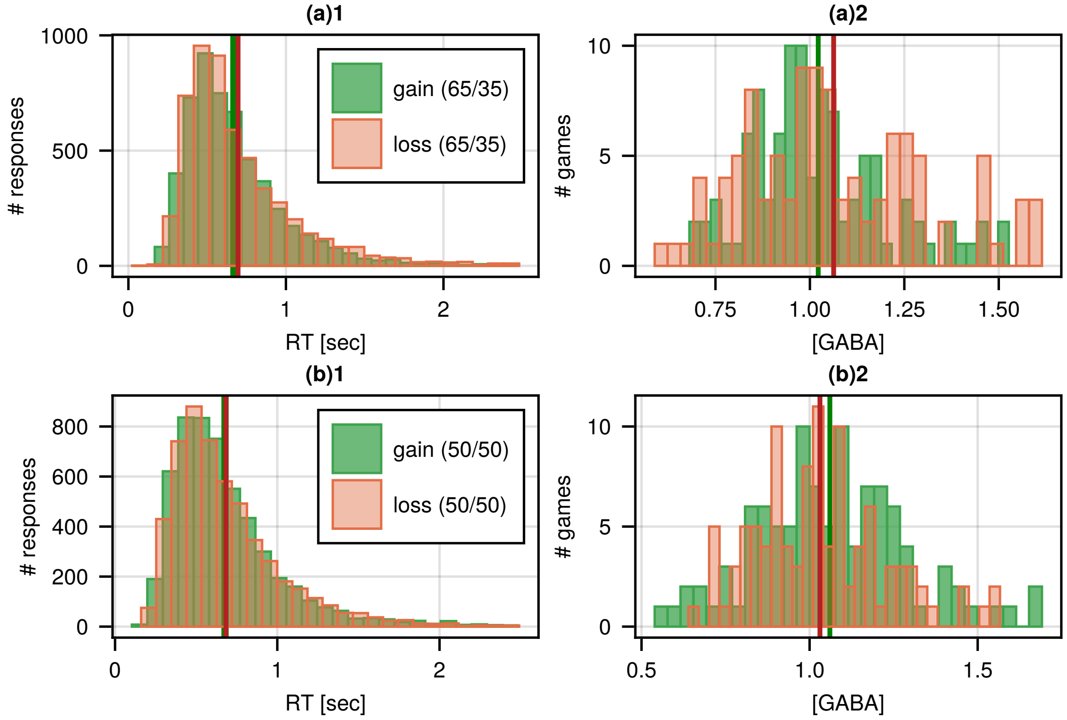


Figure S26: (a)1 RT distributions of all responses in trials 1-50 of all 107 participants in the gain/loss biased conditions (65/35). The green and red vertical lines indicate the average RTs (± STD, in sec): $0.67 \pm 0.31$ (gain), $0.70 \pm 0.35$ (loss). (a)2 Distributions of the GABA concentrations averaged over the trials in one game for all 107 participants in the gain/loss biased conditions (65/35). The green and red vertical lines indicate the average GABA concentrations (± STD): $1.02 \pm 0.19$ (gain), $1.06 \pm 0.24$ (loss). (b)1 RT distributions of all responses in trials 1-50 of all 107 participants in the gain/loss unbiased conditions (50/50). The green and red vertical lines indicate the average RTs (± STD, in sec): $0.68 \pm 0.34$ (gain), $0.68 \pm 0.33$ (loss). (b)2 Distributions of the GABA concentrations averaged over the trials in one game for all 107 participants in the gain/loss unbiased conditions (50/50). The green and red vertical lines indicate the average GABA concentrations (± STD): $1.06 \pm 0.23$ (gain), $1.03 \pm 0.20$ (loss).

Also, for each participant, the concentration of inhibitory neurotransmitter ($\gamma$-aminobutyric-acid, GABA) was measured during the game (averaged over the trials, fig. S26). The GABA and Glutamate concentrations were quantified from the dorsal anterior cingulate cortex (dACC), using Proton Magnetic Resonance Spectroscopy ($^1$H-MRS) at 7T [24].

#### B. Data analysis

The initial trials imply the learning process, where the volunteers explore the game conditions and the hidden probabilities. Therefore, we consider trials 1-28 as the learning process, after which the mean values saturate (fig. S27(a)).

The number of initial trials (28) is chosen such that a random sequence of binary choices can give 35% mistakes with less than 5% probability. In other words, if the number of mistakes in the given sequence of binary choices is lower than 35%, it is highly likely that the sequence is not random (with a significance of 5%). Therefore, the following analysis excludes the choices, RTs, and GABA concentrations in these learning trials. We also excluded all the participants who, for the given condition, did not explore both options. We required that the participants choose each option in the pair at least three times during the entire game.

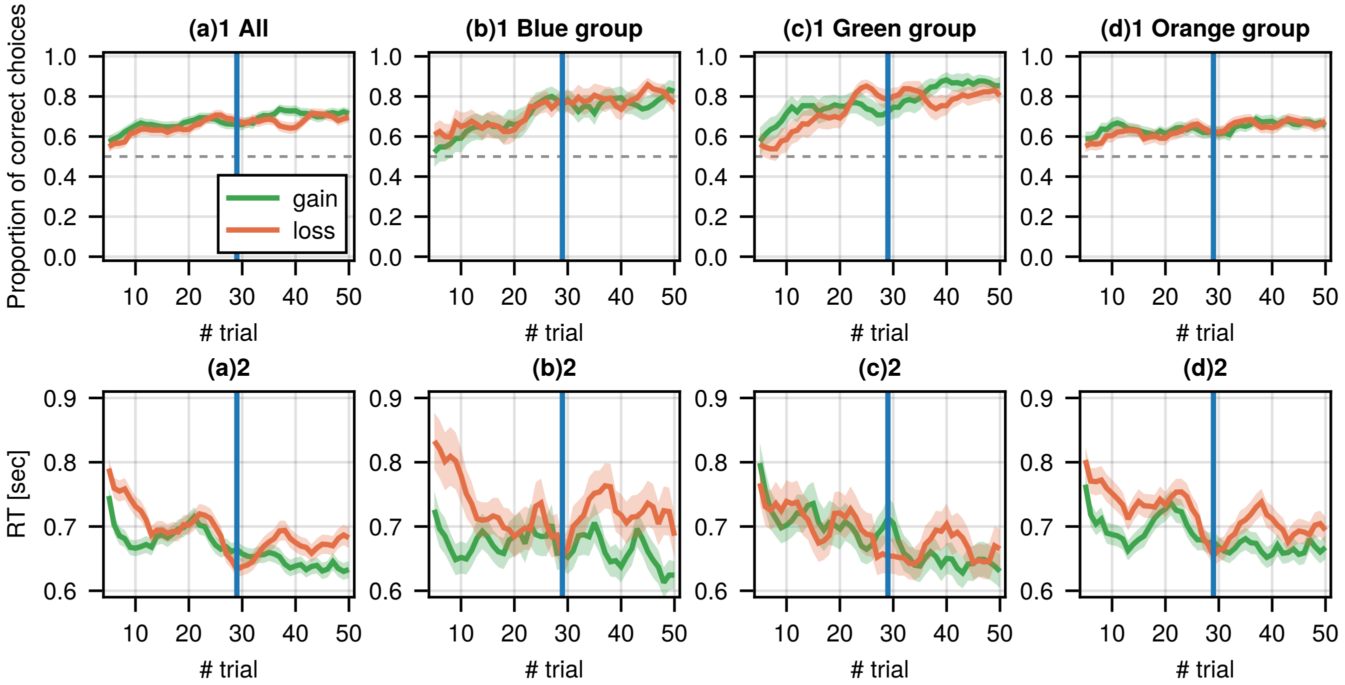


Figure S27: The proportion of correct choices (1, top) and the RTs (2, bottom) averaged over a 5-trial moving window (mean ± shaded SE, standard error of the mean). (a) All participants. (b) Participants in the blue group (see fig. 6A in the main text for the details about the division into the groups). (c) Participants in the green group. (d) Participants in the orange group. The green and red lines represent the probability of choosing the correct option in the 65/35 gain and loss conditions. The blue vertical lines indicate the learning process: trials 1-28. The grey dashed horizontal line is the probability of 0.5.

Under the unbiased conditions, when the probabilities of the zero and non-zero rewards for both options were 50%, there was no correct option so only the RTs were measured as the baseline (labeled $RT_{50/50}$). We consider the results of only those participants who did not prefer one option to the other. Thus, we remove all recordings with the error rate $< 0.23$ or $> 0.77$. These numbers are chosen such that a random sequence of choices can result in an error of 0.23 and lower with a significance of 0.01. In other words, if a sequence demonstrates an error of 0.23 or lower (similarly, 0.77 or higher), it is highly likely that this sequence is not random.

Under the biased condition (65/35), we keep the results of only those participants who learned correctly which of the options is better, so we remove all data with the error rate $\geq 0.5$. In the biased conditions (gain and loss), we also excluded one game, where the ratio $RT_c/RT_w$ was above 3.5 (the average ratio ± STD was $0.973 \pm 0.220$ for 190 observations without this outlier).

In the data set described above, we measure the error rate as the fraction of wrong choices made by each participant under each condition, the mean RT over all trials after the learning period (29-50), and the mean GABA concentrations during each game. We then average the RTs and the GABA concentrations over the participants. We also measure the mean RT in the correct decisions and the mean RT in the wrong decisions per participant under the biased conditions, take the ratio $RT_c/RT_w$ and then, average over the participants (all the measurements are presented in table S5 and fig. S28).

The gain and loss trials did not show significant differences in their error rates, RTs, and GABA concentrations (green vs. orange graphs in fig. S26, fig. S27, fig. S28; note that in table S5, the numbers of gain and loss observations are not equal due to data processing, described above). We show that by conducting the equal variance two-tailed t-test for the measurements in the biased and unbiased conditions after the learning period and presented in table S6. We, therefore, combine the gain and loss observations for the following analysis.

After the described procedure, we also find the ratio of the mean RTs in the biased and unbiased conditions (the normalized reaction time $RT_{65/35}/RT_{50/50}$) per participant. Some participants did explore both options during the games under some conditions (out of 4), while they did not explore under the other conditions. In this case, while calculating the normalized RTs in the gain and loss trials, we take the biased $RT_{65/35}$ if the game is not excluded for this participant. The unbiased $RT_{50/50}$ is calculated for both gain and loss conditions together per participant if both results of the games are maintained or for either the gain or loss condition if the other unbiased condition is excluded.

Table S5: Experimental setup II: separate trials

| **Condition** | **Parameters** | **mean ± STD** | **mean ± SE** | **# participants** |
|---|---|---|---|---|
| Gain, 65/35 | Error rate | 0.235 ± 0.145 | 0.235 ± 0.016 | 80 |
| | RT [sec] | 0.654 ± 0.193 | 0.654 ± 0.022 | 80 |
| | $RT_c/RT_w$ | 0.935 ± 0.220 | 0.935 ± 0.026 | 70 |
| | GABA | 1.021 ± 0.181 | 1.021 ± 0.021 | 74 |
| Loss, 65/35 | Error rate | 0.237 ± 0.137 | 0.237 ± 0.016 | 78 |
| | RT [sec] | 0.688 ± 0.232 | 0.688 ± 0.026 | 78 |
| | $RT_c/RT_w$ | 0.978 ± 0.211 | 0.978 ± 0.025 | 72 |
| | GABA | 1.076 ± 0.223 | 1.076 ± 0.025 | 77 |
| Gain, 50/50 | Error rate | 0.496 ± 0.139 | 0.496 ± 0.015 | 88 |
| | RT [sec] | 0.655 ± 0.234 | 0.655 ± 0.025 | 88 |
| | GABA | 1.047 ± 0.218 | 1.047 ± 0.024 | 85 |
| Loss, 50/50 | Error rate | 0.517 ± 0.123 | 0.517 ± 0.013 | 89 |
| | RT [sec] | 0.677 ± 0.205 | 0.677 ± 0.022 | 89 |
| | GABA | 1.027 ± 0.200 | 1.027 ± 0.021 | 87 |

Experimental results were calculated for 107 volunteers who participated in four games of 50 trials of two-choice tasks under uncertainty under the gain/loss and biased/unbiased conditions. The values are calculated in the trials after the learning period (trials 29-50). In the unbiased case (50/50), when the two options are equivalent, we use the error rate to denote the proportion of choosing one of the options over the other.

Table S6: Statistical analysis for the averaged quantities, presented in table S5.

| **Hypotheses for 65/35** | **t-test** |
|---|---|
| Error(gain) ≠ Error(loss) | df = 156, t = -0.0561, p = 0.9553 |
| RT(gain) ≠ RT(loss) | df = 156, t = -0.9983, p = 0.3197 |
| GABA(gain) ≠ GABA(loss) | df = 149, t = -1.6462, p = 0.1018 |
| $RT_c/RT_w$(gain) ≠ $RT_c/RT_w$(loss) | df = 140, t = -1.1774, p = 0.241 |
| $RT_c/RT_w$(gain) ≠ 1 | df = 69, t = -2.4682, **p = 0.0161** |
| $RT_c/RT_w$(loss) ≠ 1 | df = 71, t = -0.9025, p = 0.3699 |
| **Hypotheses for 50/50** | **t-test** |
| Error(gain) ≠ Error(loss) | df = 175, t = -1.0984, p = 0.2736 |
| RT(gain) ≠ RT(loss) | df = 175, t = -0.6671, p = 0.5056 |
| GABA(gain) ≠ GABA(loss) | df = 170, t = 0.6497, p = 0.5168 |

We conduct the equal variance two-tailed t-test for comparison and show that the error rate, the RTs, the GABA concentrations, and the ratios $RT_c/RT_w$ are not significantly different in the gain and loss conditions for both biased (65/35) and unbiased (50/50) conditions. In the unbiased case, when the two options are equivalent, we use the error rate to denote the proportion of choosing one of the options. We also compare the ratios $RT_c/RT_w$ to 1 and find that it is significantly lower than 1 in the gain condition.

### C. Fitting the IIM to experiment II

To show the robustness and self-consistency of the IIM, we repeat the analysis shown in sec. "Setup II" in the main text for two other ways of dividing the data into three groups with two lower error thresholds (0.1 and 0.15 instead of 0.2, used in the main text).

The observations (under both gain and loss conditions) in the biased case (65/35) are divided into three groups such that in the green group, the error rate per participant in a single game is lower than the error threshold (0.2, 0.1, or 0.15), and the normalized RT ($RT_{65/35}/RT_{50/50}$) is below 1. In the blue group, the error rate is also lower than the error threshold, and the normalized RT is above 1. In the orange group, the error rate is above 0.2.

We show the learning curves and the RTs for each group in fig. S27. The summary of the experimental data is shown in table S7, table S8 and plotted in fig. 6A(ii) in the main text (for the error threshold of 0.2), fig. S29(a)(ii) (0.1), and fig. S30(a)(ii) (0.15).

These cases with the lower error thresholds (0.1, 0.15) show similar results to what we find in the main text (0.2). First, the average GABA concentration in the biased case in the blue group is larger than in the green group for all error thresholds, while in the unbiased case, the two concentrations are similar (table S7). We conduct the unequal variance two-tailed t-test for comparison (table S9).

We then mark the region on the IIM's phase diagram that satisfies the experimental observations (data: table S8, results: fig. S29, fig. S30). The detailed procedure is described in sec. "Setup II" in the main text. It turns out that for all three approaches with different error thresholds, we obtain similar regions on the phase diagram near the tricritical point, which supports the robustness of our IIM.

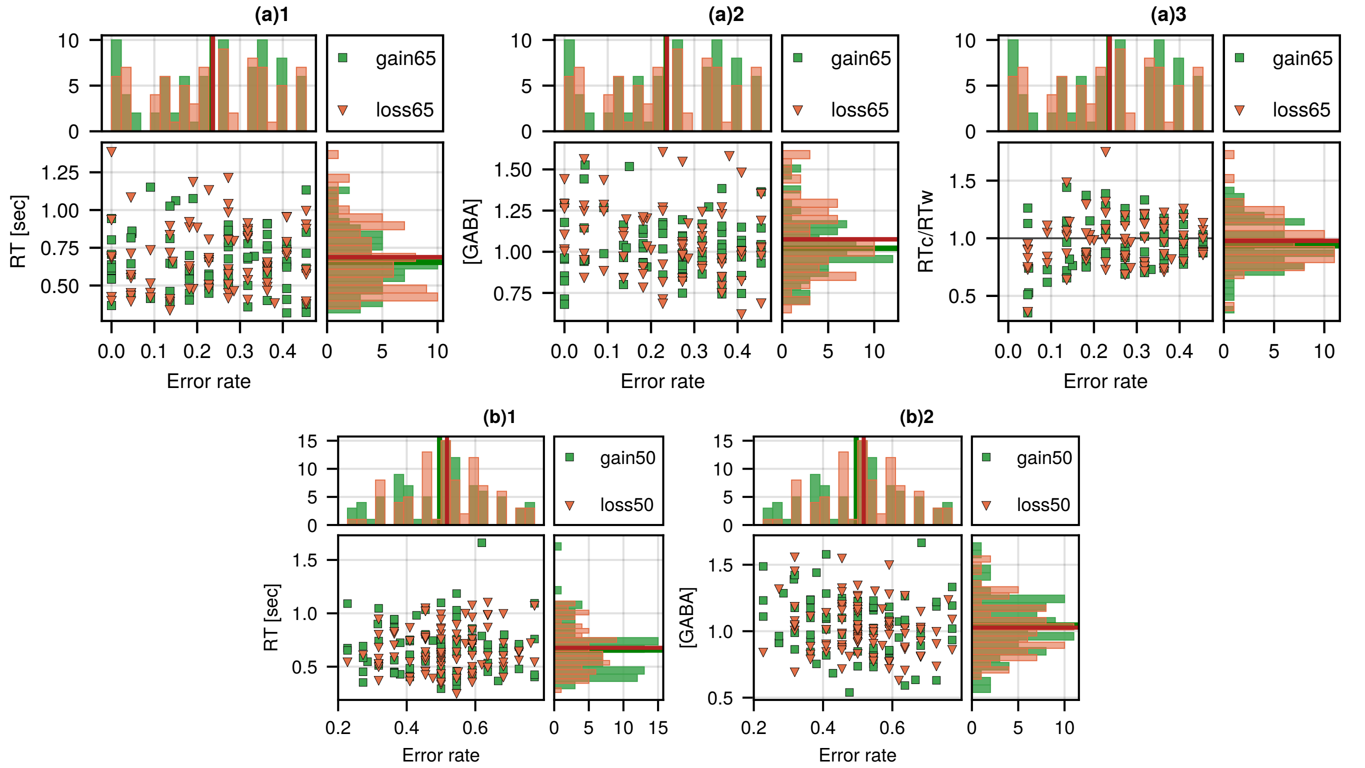


Figure S28: (a) Measurements in the biased (65/35) gain and loss conditions after the learning process (trials 29-50). (a)1 The average RT vs. the error rate. (a)2 The average GABA concentration vs. the error rate. (a)3 The average ratio $RT_c/RT_w$ vs. the error rate (b) Measurements in the unbiased (50/50) gain and loss conditions after the learning process (trials 29-50). (b)1 The average RT vs. the error rate. (b)2 The average GABA concentration vs. the error rate. Each green square (gain) and orange triangle (loss) indicate the result for a single participant. Top histogram in each subfigure: the histogram of the error rate per participant. The green and red vertical lines indicate the average error rates of the participants (table S5). Right histogram in each subfigure: the histogram of the average RTs, GABA concentrations, and the average ratios $RT_c/RT_w$ per participant. The green and red horizontal lines indicate the average values over the participants (table S5). The black horizontal line in (a)3 is the ratio of 1.

Table S7: GABA concentrations, measured in experimental setup II

| Error Threshold | Parameter | Mean ± STD | Mean ± SE | # observations |
|---|---|---|---|---|
| 0.2 | $GABA_{65/35}$(green) | 1.016 ± 0.168 | 1.016 ± 0.034 | 25 |
| | $GABA_{50/50}$(green) | 1.057 ± 0.207 | 1.057 ± 0.040 | 27 |
| | $GABA_{65/35}$(blue) | 1.127 ± 0.249 | 1.127 ± 0.050 | 25 |
| | $GABA_{50/50}$(blue) | 1.091 ± 0.243 | 1.091 ± 0.049 | 25 |
| 0.1 | $GABA_{65/35}$(green) | 1.014 ± 0.205 | 1.014 ± 0.057 | 13 |
| | $GABA_{50/50}$(green) | 1.024 ± 0.242 | 1.024 ± 0.065 | 14 |
| | $GABA_{65/35}$(blue) | 1.2 ± 0.285 | 1.2 ± 0.082 | 12 |
| | $GABA_{50/50}$(blue) | 1.117 ± 0.266 | 1.117 ± 0.077 | 12 |
| 0.15 | $GABA_{65/35}$(green) | 0.998 ± 0.182 | 0.998 ± 0.042 | 19 |
| | $GABA_{50/50}$(green) | 1.053 ± 0.23 | 1.053 ± 0.05 | 21 |
| | $GABA_{65/35}$(blue) | 1.168 ± 0.26 | 1.168 ± 0.065 | 16 |
| | $GABA_{50/50}$(blue) | 1.078 ± 0.25 | 1.078 ± 0.062 | 16 |

GABA concentrations averaged over participants in biased (65/35) and unbiased (50/50) conditions in trials 29-50 after the learning period (combined gain and loss conditions). The observations are divided into groups such that in the green group, the error rate per participant in a single game is lower than the error threshold (0.2, 0.1, or 0.15), and the RT is below 1. In the blue group, the error rate is also lower than the error threshold, and the RT is above 1. The data points are presented in fig. 6A(ii) in the main text, fig. S29(a)(ii), and fig. S30(a)(ii).

### D. RT distributions

Similarly to setup I (section S8), we compare the RT in the experiments to the IIM's simulations.

We first aim to compare the RT distributions in the experiments with the IIM's simulations for different areas of the phase space. We take the same three points in the ordered, intermittent, and disordered phases (marked in (fig. S31(a)1, (b)1)) and find the biases that satisfy the experimental error rate and the normalized RT ratio ($RT_{65/35}/RT_{50/50}$) for the green group and only the error rate for the blue group (see section S7). We then plot the

Table S8: Summarized data for the experimental setup II for three different error thresholds

| **Error Threshold** | **Parameter** | **Mean ± SE** |
|---|---|---|
| 0.2 | Error rate(green, 65/35) | 0.09 ± 0.014 |
| | $RT_{65/35}/RT_{50/50}$(green) | 0.807 ± 0.023 |
| | $RT_c/RT_w$(green, 65/35) | 0.927 ± 0.054 |
| | Error rate(blue, 65/35) | 0.105 ± 0.014 |
| | $RT_c/RT_w$(blue, 65/35) | 0.91 ± 0.058 |
| | $GABA_{65/35}$(blue)/$GABA_{65/35}$(green) | 1.1089 ± 0.0857 |
| | $RT_{65/35}$(blue)/$RT_{65/35}$(green) | 1.3079 ± 0.1676 |
| 0.1 | Error rate(green, 65/35) | 0.026 ± 0.009 |
| | $RT_{65/35}/RT_{50/50}$(green) | 0.811 ± 0.03 |
| | $RT_c/RT_w$(green, 65/35) | 0.843 ± 0.067 |
| | Error rate(blue, 65/35) | 0.034 ± 0.01 |
| | $RT_c/RT_w$(blue, 65/35) | 0.709 ± 0.097 |
| | $GABA_{65/35}$(blue)/$GABA_{65/35}$(green) | 1.1826 ± 0.1474 |
| | $RT_{65/35}$(blue)/$RT_{65/35}$(green) | 1.4389 ± 0.2683 |
| 0.15 | Error rate(green, 65/35) | 0.063 ± 0.013 |
| | $RT_{65/35}/RT_{50/50}$(green) | 0.809 ± 0.025 |
| | $RT_c/RT_w$(green, 65/35) | 0.912 ± 0.065 |
| | Error rate(blue, 65/35) | 0.065 ± 0.014 |
| | $RT_c/RT_w$(blue, 65/35) | 0.864 ± 0.094 |
| | $GABA_{65/35}$(blue)/$GABA_{65/35}$(green) | 1.1698 ± 0.1142 |
| | $RT_{65/35}$(blue)/$RT_{65/35}$(green) | 1.3305 ± 0.2048 |

Summarized data for the experimental setup II for three different error thresholds, used for the IIM fitting (fig. S29, fig. S30). The group names relate to the colors in fig. S29(a)(i), fig. S30(a)(i).

Table S9: Statistical analysis for the averaged GABA concentrations in experimental setup II

| **Error Threshold** | **Hypotheses** | **t-test** |
|---|---|---|
| 0.2 | $GABA_{50/50}$(green) ≠ $GABA_{50/50}$(blue) | df = 47.43, t = -0.5404, p = 0.5914 |
| | $GABA_{65/35}$(green) ≠ $GABA_{65/35}$(blue) | df = 42.16, t = -1.8419, **p = 0.0725** |
| | $GABA_{65/35}$(green) ≠ $GABA_{50/50}$(green) | df = 49.18, t = 0.7772, p = 0.4408 |
| | $GABA_{65/35}$(blue) ≠ $GABA_{50/50}$(blue) | df = 47.97, t = -0.5198, p = 0.6056 |
| 0.1 | $GABA_{50/50}$(green) ≠ $GABA_{50/50}$(blue) | df = 22.55, t = -0.9322, p = 0.3611 |
| | $GABA_{65/35}$(green) ≠ $GABA_{65/35}$(blue) | df = 19.83, t = -1.8523, **p = 0.0789** |
| | $GABA_{65/35}$(green) ≠ $GABA_{50/50}$(green) | df = 24.79, t = 0.1072, p = 0.9155 |
| | $GABA_{65/35}$(blue) ≠ $GABA_{50/50}$(blue) | df = 21.9, t = -0.7309, p = 0.4726 |
| 0.15 | $GABA_{50/50}$(green) ≠ $GABA_{50/50}$(blue) | df = 30.96, t = -0.3239, p = 0.7482 |
| | $GABA_{65/35}$(green) ≠ $GABA_{65/35}$(blue) | df = 26.19, t = -2.1913, **p = 0.0375** |
| | $GABA_{65/35}$(green) ≠ $GABA_{50/50}$(green) | df = 37.37, t = 0.8343, p = 0.4094 |
| | $GABA_{65/35}$(blue) ≠ $GABA_{50/50}$(blue) | df = 29.95, t = -0.9873, p = 0.3314 |

Statistical analysis for the averaged GABA concentrations in the blue and green groups of participants in the biased (65/35) and unbiased (50/50) conditions, presented in table S7. We conduct the unequal variance two-tailed t-test for comparison and show that the GABA concentrations are not significantly different in the biased (65/35) and unbiased (50/50) conditions for both green and blue groups, while the GABA concentrations are marginally different in the two groups in the biased case (bold font).

normalized RT distributions in the experiments and in the IIM's simulations (divided by the mean RT, fig. S31(a)2, (b)2). The typical decision trajectories correspond to the diffusion process in the disordered phase and run-and-tumble or ballistic in the ordered and intermittent phases (fig. S31(a)3, (b)3).

The differences between the experimental data in setup II (x-axis) and the three distributions (y-axis) are quantified using the quantile-quantile plot [21, 22] (fig. S31(a)4, (b)4) and the distribution parameters (table S10).

Overall, the distributions in all phases are different from the data in the green group, while the intermittent phase shows a similar distribution to the observed data in the blue group. This is different from the result for setup I, where the observed distributions were similar to the RT distribution in the ordered phase (section S8, fig. S23).

#### E. Fitting the DDM to experiment II

As we show in fig. 6B in the main text, the disordered phase of the IIM does not fit to the experimental data, as we now demonstrate using the analytical solutions of the DDM for the normalized RT ($RT_{65/35}/RT_{50/50}$). We assume

Table S10: Comparison of the parameters of the normalized RT distributions, given in different phases, vs. experimental data (setup II).

| **Parameters** | **Mean** | **Median** | **STD** | **Skewness** | **Kurtosis** | $L/V^{+}_{\mathbf{MF}}$ | **# simulations** |
|---|---|---|---|---|---|---|---|
| Intermittent: $T = 0.18, \eta = 0.45, \epsilon_1 = 0.063$ | 1 | 0.97 | 0.13 | 6.35 | 69.53 | 0.88 | $10^5$ |
| Ordered: $T = 0.3, \eta = 0.36, \epsilon_1 = 0.077$ | 1 | 0.86 | 0.38 | 3.23 | 12.36 | 0.8 | $2 \times 10^4$ |
| Disordered: $T = 0.44, \eta = 0.56, \epsilon_1 = 0.0045$ | 1 | 0.8 | 0.75 | 1.65 | 3.6 | 1.21 | $10^3$ |
| Green (data) | 1 | 0.88 | 0.46 | 1.88 | 4.2 | – | 592 |
| Intermittent: $T = 0.18, \eta = 0.5, \epsilon_1 = 0.026$ | 1 | 0.81 | 0.57 | 2.14 | 6.74 | 0.44 | $10^5$ |
| Ordered: $T = 0.3, \eta = 0.4, \epsilon_1 = 0.0395$ | 1 | 0.75 | 0.54 | 2.47 | 8.1 | 0.61 | $2 \times 10^4$ |
| Disordered: $T = 0.44, \eta = 0.56, \epsilon_1 = 0.004$ | 1 | 0.77 | 0.79 | 1.79 | 4.35 | 1.28 | $10^3$ |
| Blue (data) | 1 | 0.86 | 0.46 | 1.37 | 2.28 | – | 562 |

The parameters of IIM ($\eta$, $T$) are marked in fig. S31. $RT^{+}_{bal} = L/V^{+}_{\mathrm{MF}}$ (eq. (8)) indicates the theoretical prediction for the mean RT using the MF velocity.

equal thresholds for the two options and write the quantities in terms of the dimensionless Péclet number $\mathrm{Pe} = vL/D$, which characterizes the ratio between the diffusion and convection time scales (see also SI section S3 C):

$$\begin{cases} Error = \frac{1}{e^{\mathrm{Pe}}+1} \\ \mathrm{RT} = \mathrm{RT_c} = \mathrm{RT_w} = \frac{L}{v} \tanh\left(\frac{\mathrm{Pe}}{2}\right) \end{cases} \tag{S46}$$

where the $Error$ is the probability of reaching the negative decision threshold $-L$, $v$ is a constant drift, and $D$ is the diffusion coefficient.

We assume that the constant drift $v$ plays a role of bias in the system, so we fix the diffusion coefficient $D$ and the threshold $L$. Then, we express the unbiased $\mathrm{RT}_{50/50}$ at the zero drift:

$$\mathrm{RT}_{50/50} = \lim_{v \to 0} \frac{L}{v} \tanh\left(\frac{vL}{2D}\right) = \frac{L^2}{2D} \tag{S47}$$

And the RT as a function of the error rate is

$$\mathrm{RT}_{65/35} = \frac{L^2\left(1 - 2\,Error_{65/35}\right)}{D \ln\left(\frac{1}{Error_{65/35}} - 1\right)} \tag{S48}$$

Then, the normalized RT can be written in the DDM as follows:

$$\frac{\mathrm{RT}_{65/35}}{\mathrm{RT}_{50/50}} = \frac{2\left(1 - 2\,Error_{65/35}\right)}{\ln\left(\frac{1}{Error_{65/35}} - 1\right)} \tag{S49}$$

Plugging the experimental values (table II in the main text) into the analytical solutions, we find the DDM's RT ratio for any error rate in the biased case (65/35), presented in fig. S32. It turns out that indeed, the DDM's analytical solution does not fit the experimental results for both green and blue groups, which supports the finding in the IIM. At the same time, the orange group exhibits a marginal fit.

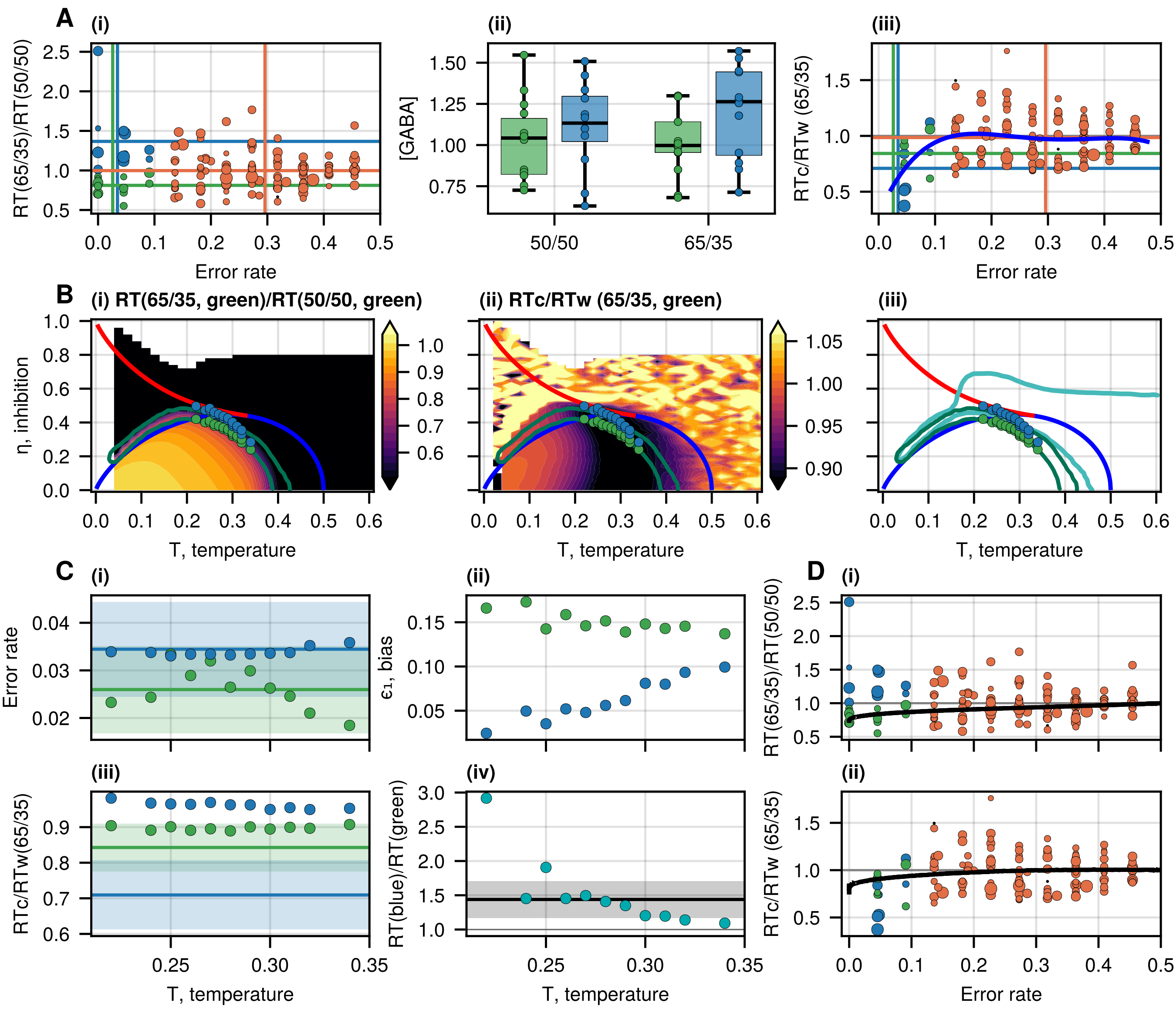


Figure S29: The error threshold is 0.1. (a)(i) Normalized RT ($RT_{65/35}/RT_{50/50}$) as a function of the error rate. Each RT for the gain or loss trials (with the probabilities of 65%-35% per choice) is normalized by the participant's RT in the unbiased condition (with the reward probability of 50% per choice). Each point represents a result from a single participant, for gain and loss separately. The size of the points is related to the average GABA concentration measured for each participant during the task. The data is divided into three groups: the green group indicates the volunteers whose error rate $\leq 0.1$ and the normalized RT $\leq 1$, the blue group has error rate $\leq 0.1$ and the RT $> 1$, while the orange group exhibits error rate $> 0.1$. The vertical and horizontal lines denote the average error rate and normalized RT for each group. (a)(ii) The GABA concentration, quantified from the dorsal anterior cingulate cortex (dACC) during the task for the unbiased and biased trials for the green and blue groups. We find that the concentration of $GABA_{50/50}$(green) is not different from $GABA_{50/50}$(blue) (unequal variance t-test: p = 0.3611, see in table S9), though for the biased conditions, the concentration $GABA_{65/35}$(green) shows a marginal difference with $GABA_{65/35}$(blue) (unequal variance t-test: p = 0.0789). (a)(iii) The ratio $RT_c/RT_w$ for the same groups of (a)(i) at the biased conditions (both gain and loss). Each point represents a result from a single volunteer. The size of the points is related to the average GABA concentration of each participant during the task. The blue line indicates a 4th-degree polynomial fitted to the data as a guide to the eye. (b)(i) The normalized RT ratio (between the biased and unbiased conditions) for the average error rate of the green group (table S8), as given by the IIM. For each point of the phase diagram, we find the bias that satisfies the error rate of the green group ($0.03 \pm 0.01$), and this gives us the biased $RT_{65/35}$, while the unbiased $RT_{50/50}$ is calculated for $\epsilon_1 = 0$. The green circles correspond to the green group's average normalized RT and $RT_c/RT_w$ (table S8). The blue circles denote a shift of the green circles by increasing the global inhibition by a factor of 1.18, which is the ratio of the measured average GABA concentrations in the two groups for the biased conditions (table S8). The red and blue lines on the heatmaps denote the first and second-order transitions, respectively. The dark green contour denotes the region that fits the green group's RT ratio (without constraining the ratio of $RT_c/RT_w$) for the error threshold 0.2 (see fig. 6B in the main text). (b)(ii) Heatmap of the $RT_c/RT_w$ given by the IIM for the average error rate of the green group at the biased conditions (table S8). (b)(iii) The comparison of the areas of the phase space that fit the experimental data in setups I and II. The turquoise contour is a guide to the eye, which indicates the area of the phase space that best matched the experimental data of setup I (fig. 5C in the main text). (c)(i-ii) The error rate and biases of the green and blue circles from (b), as a function of temperature $T$. The calculated error rates agree with the mean values of the experimental observations (denoted by the horizontal lines and shading). The higher error rate for the blue circles corresponds to lower biases. (c)(iii) The ratio $RT_c/RT_w$ for the green and blue circles in (b). The green circles agree well with the experimental observation (denoted by the horizontal lines and shading), while the blue circles do not match the observed data. (c)(iv) The ratio of the RTs for the green and blue circles in (b) as a function of temperature $T$. The black line and the shaded area indicate the ratio of the average biased RTs between the blue and green groups in the experiment: $1.44 \pm 0.27$. (d)(i) The experimental data for the normalized RT (as in fig. S29(a)(i)) compared to the IIM model (the black line). We plot the normalized RT for one of the green points of (b) ($T = 0.28$, $\eta = 0.36$) by varying the bias from a small value (where the error rate approaches 0.5) to a large value (where the error rate approaches zero). Similarly, this gives us the $RT_c/RT_w$ as shown in (d)(ii).

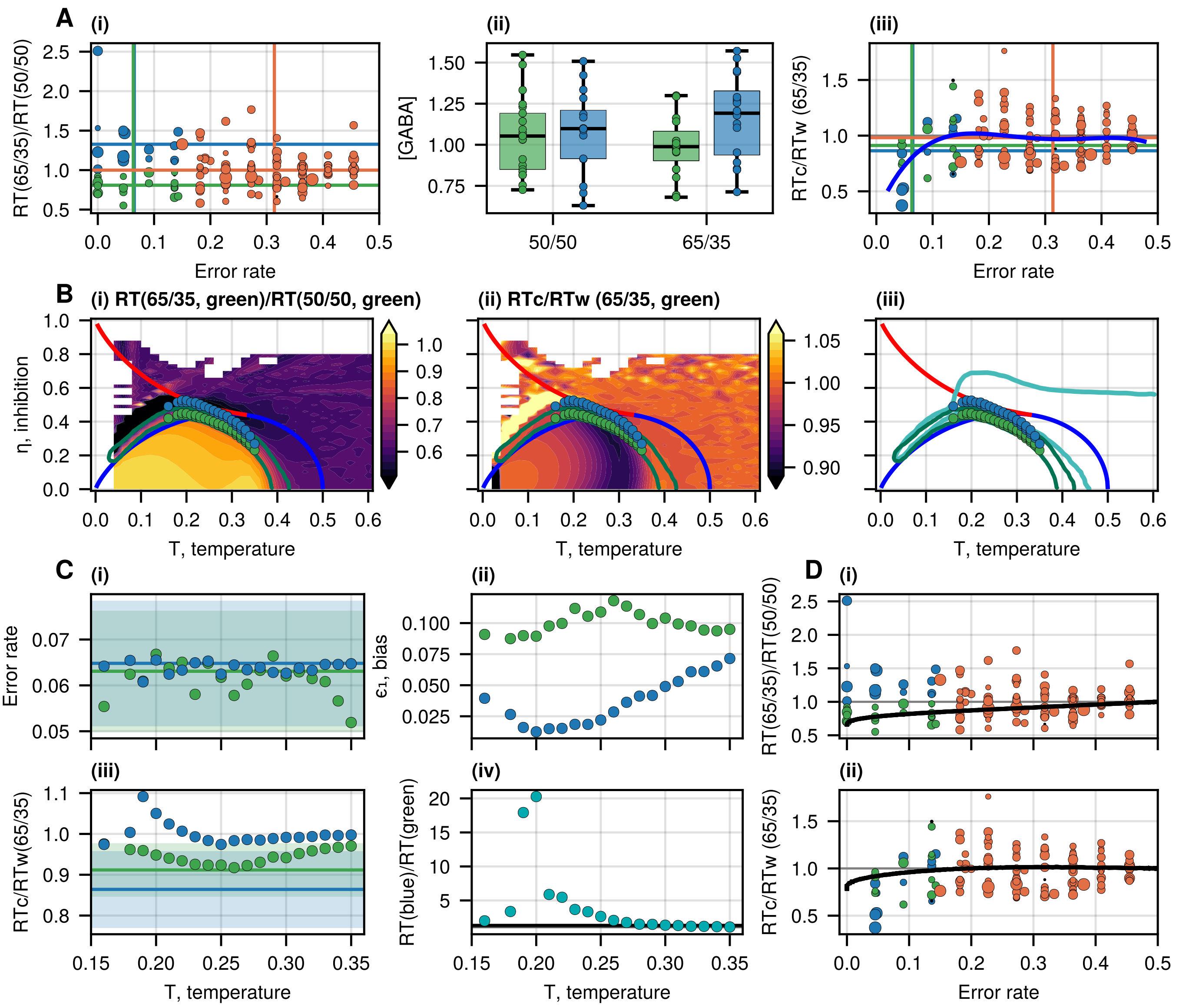


Figure S30: The error threshold is 0.15. (a)(i) Normalized RT ($RT_{65/35}/RT_{50/50}$) as a function of the error rate. Each RT for the gain or loss trials (with the probabilities of 65%-35% per choice) is normalized by the participant's RT in the unbiased condition (with the reward probability of 50% per choice). Each point represents a result from a single participant, for gain and loss separately. The size of the points is related to the average GABA concentration measured for each participant during the task. The data is divided into three groups: the green group indicates the volunteers whose error rate $\leq 0.15$ and the normalized RT $\leq 1$, the blue group has error rate $\leq 0.15$ and the RT $> 1$, while the orange group exhibits error rate $> 0.15$. The vertical and horizontal lines denote the average error rate and normalized RT for each group. (a)(ii) The GABA concentration, quantified from the dorsal anterior cingulate cortex (dACC) during the task for the unbiased and biased trials for the green and blue groups. We find that the concentration of $GABA_{50/50}$(green) is not different from $GABA_{50/50}$(blue) (unequal variance t-test: $p = 0.7482$, see in table S9), though for the biased conditions, the concentration $GABA_{65/35}$(green) shows a marginal difference with $GABA_{65/35}$(blue) (unequal variance t-test: $p = 0.0375$). (a)(iii) The ratio $RT_c/RT_w$ for the same groups of (a)(i) at the biased conditions (both gain and loss). Each point represents a result from a single volunteer. The size of the points is related to the average GABA concentration of each participant during the task. The blue line indicates a 4th-degree polynomial fitted to the data as a guide to the eye. (b)(i) The normalized RT ratio (between the biased and unbiased conditions) for the average error rate of the green group (table S8), as given by the IIM. For each point of the phase diagram, we find the bias that satisfies the error rate of the green group ($0.06 \pm 0.01$), and this gives us the biased $RT_{65/35}$, while the unbiased $RT_{50/50}$ is calculated for $\epsilon_1 = 0$. The green circles correspond to the green group's average normalized RT and $RT_c/RT_w$ (table S8). The blue circles denote a shift of the green circles by increasing the global inhibition by a factor of 1.17, which is the ratio of the measured average GABA concentrations in the two groups for the biased conditions (table S8). The red and blue lines on the heatmaps denote the first and second-order transitions, respectively. The dark green contour denotes the region that fits the green group's RT ratio (without constraining the ratio of $RT_c/RT_w$) for the error threshold 0.2 (see fig. 6B in the main text). (b)(ii) Heatmap of the $RT_c/RT_w$ given by the IIM for the average error rate of the green group at the biased conditions (table S8). (b)(iii) The comparison of the areas of the phase space that fit the experimental data in setups I and II. The turquoise contour is a guide to the eye, which indicates the area of the phase space that best matched the experimental data of setup I (fig. 5C in the main text). (c)(i-ii) The error rate and biases of the green and blue circles from (b), as a function of temperature $T$. The calculated error rates agree with the mean values of the experimental observations (denoted by the horizontal lines and shading). The higher error rate for the blue circles corresponds to lower biases. (c)(iii) The ratio $RT_c/RT_w$ for the green and blue circles in (b). The green circles agree well with the experimental observation (denoted by the horizontal lines and shading), while the blue circles do not match the observed data. (c)(iv) The ratio of the RTs for the green and blue circles in (b) as a function of temperature $T$. The black line and the shaded area indicate the ratio of the average biased RTs between the blue and green groups in the experiment: $1.33 \pm 0.2$. (d)(i) The experimental data for the normalized RT (as in fig. S30(a)(i)) compared to the IIM model (the black line). We plot the normalized RT for one of the green points of (b) ($T = 0.28$, $\eta = 0.38$) by varying the bias from a small value (where the error rate approaches 0.5) to a large value (where the error rate approaches zero). Similarly, this gives us the $RT_c/RT_w$ as shown in (d)2.

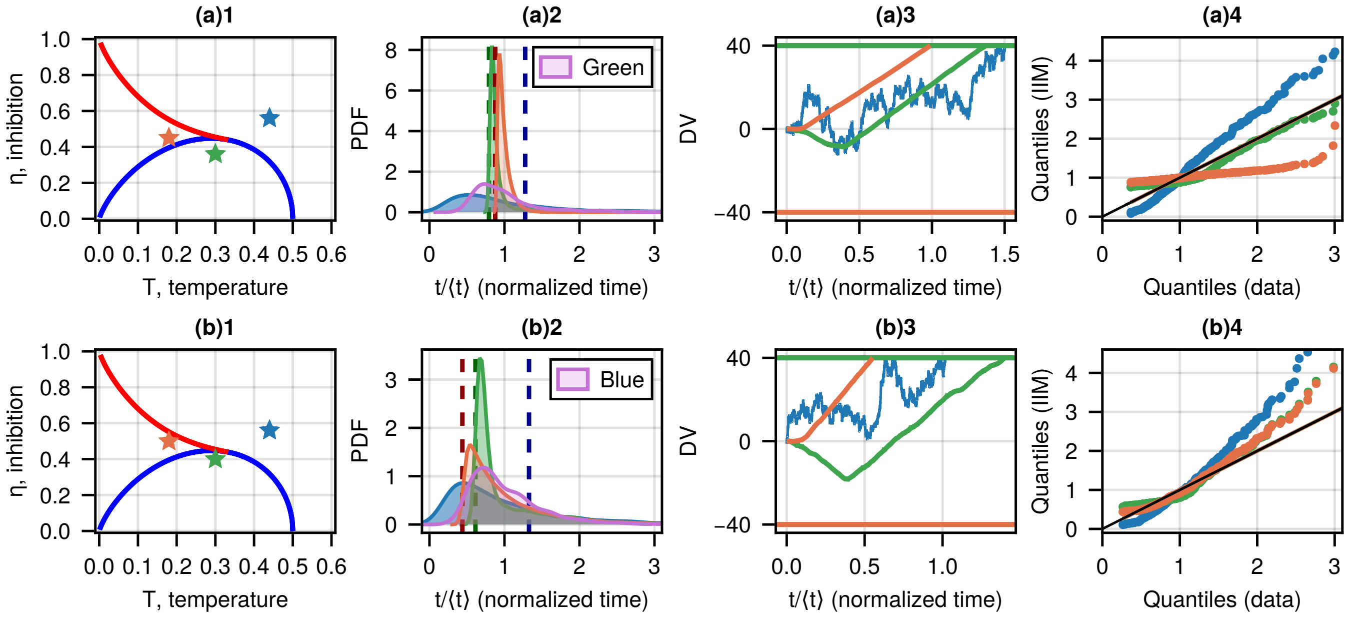


Figure S31: (a)1 Phase diagram of the IIM. The red and blue lines on the heatmaps denote the first and second-order transitions, respectively. The stars indicate the parameters ($\eta$, $T$) in the three phases from which we sample the IIM's RT distributions to compare with the experimental data. (a)2 The RT distributions (normalized by the mean RT) for the three regimes (denoted in (a)1) and the observed RT distribution in the trials after the learning period for the green group (purple area). The bias for each point is chosen such that the predicted error rate for this point satisfies the observed error rate for the green group and the mean normalized RT in the experiment for the green group ($\mathrm{RT}_{65/35}/\mathrm{RT}_{50/50}$) (see table S10). The vertical dashed lines indicate the theoretical RT given by $RT^{+}_{bal}$ (eq. (8) in the main text). (a)3 Typical trajectories during the decision-making process corresponding to the IIM's RT distributions shown in (a)2. (a)4 Comparison of the IIM's RT distributions with the observed data (black identity line) using a quantile-quantile representation. (b) Comparison for the blue group. The bias is chosen such that the error rate in the IIM simulations satisfies the observed error rate in the blue group (see table S10).

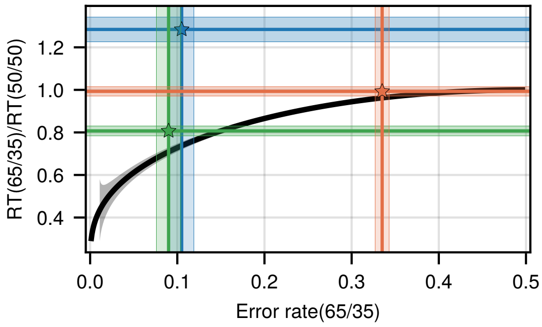


Figure S32: DDM vs. setup II. Analytical ratio of the mean RTs in the biased and unbiased conditions ($\mathrm{RT}_{65/35}/\mathrm{RT}_{50/50}$) for the DDM model as a function of the error in the biased condition (eq. (S49)), represented by the black line. The green, blue, and orange lines, the shaded rectangles, and the stars indicate the error rates and the RT ratios in the experiment (table II in the main text).

[1] V. H. Sridhar, L. Li, D. Gorbonos, M. Nagy, B. R. Schell, T. Sorochkin, N. S. Gov, and I. D. Couzin, Proceedings of the National Academy of Sciences **118**, e2102157118 (2021), publisher: Proceedings of the National Academy of Sciences.

[2] J. Bezanson, A. Edelman, S. Karpinski, and V. B. Shah, SIAM Review **59**, 65 (2017), publisher: Society for Industrial and Applied Mathematics.

[3] R. Bogacz, E. Brown, J. Moehlis, P. Holmes, and J. D. Cohen, Psychological Review **113**, 700 (2006), place: US Publisher: American Psychological Association.

[4] G. Volpe and G. Volpe, American Journal of Physics **81**, 224 (2013).

[5] C. W. Gardiner, *Handbook of Stochastic Methods: For Physics, Chemistry and Natural Sciences*, 2nd ed., Springer series in synergetics No. 13 (Springer, Berlin Heidelberg, 1985).

[6] S. Redner, *A Guide to First-Passage Processes* (Cambridge University Press, Cambridge, 2008).

[7] E. Roldán, I. Neri, M. Dörpinghaus, H. Meyr, and F. Jülicher, Physical Review Letters **115**, 250602 (2015), publisher: American Physical Society.

[8] R. Ratcliff and J. N. Rouder, Psychological Science **9**, 347 (1998), publisher: SAGE Publications Inc.

[9] S. Verdonck and F. Tuerlinckx, Psychological Review **121**, 422 (2014).

[10] K. Malakar, V. Jemseena, A. Kundu, K. V. Kumar, S. Sabhapandit, S. N. Majumdar, S. Redner, and A. Dhar, Journal of Statistical Mechanics: Theory and Experiment **2018**, 043215 (2018), publisher: IOP Publishing and SISSA.

[11] H. C. Berg, *Random Walks in Biology: New and Expanded Edition*, first princeton paperback printing, expanded edition ed., Princeton paperbacks Biology (Princeton University Press, Princeton, NJ Chichester, 1993).

[23] T. Finkelman, E. Furman-Haran, K. C. Aberg, R. Paz, and A. Tal, Inhibitory mechanisms in the prefrontal-cortex differentially mediate Putamen activity during valence-based learning (2024), pages: 2024.07.29.605168 Section: New Results.

[24] T. Finkelman, E. Furman-Haran, R. Paz, and A. Tal, NeuroImage **247**, 118810 (2022).
